# Plasmodesmata act as unconventional membrane contact sites regulating inter-cellular molecular exchange in plants

**DOI:** 10.1101/2023.12.18.572149

**Authors:** Jessica Pérez-Sancho, Marija Smokvarska, Gwennogan Dubois, Marie Glavier, Sujith Sritharan, Tatiana Souza Moraes, Hortense Moreau, Victor Dietrich, Matthieu Pierre Platre, Andrea Paterlini, Ziqiang Patrick Li, Laetitia Fouillen, Magali S. Grison, Pepe Cana-Quijada, Françoise Immel, Valerie Wattelet, Mathieu Ducros, Lysiane Brocard, Clément Chambaud, Yongming Luo, Priya Ramakrishna, Vincent Bayle, Linnka Lefebvre-Legendre, Stéphane Claverol, Matej Zabrady, Wolfgang Busch, Marie Barberon, Jens Tilsner, Yrjö Helariutta, Eugenia Russinova, Antoine Taly, Yvon Jaillais, Emmanuelle M. Bayer

**Affiliations:** Laboratoire de Biogenèse Membranaire, UMR5200, CNRS, Université de Bordeaux, Villenave d’Ornon, France; Laboratoire Reproduction et Développement des Plantes, Université de Lyon, ENS de Lyon, UCB Lyon 1, CNRS, INRA, F-69342 Lyon, France; Laboratoire de Biochimie Théorique, UPR9080, CNRS, Université Paris Cité, Paris, France; Salk Institute For Biological Studies, Plant Molecular and Cellular Biology Laboratory, 10010 North Torrey Pines Road, La Jolla, CA 92037, USA; The Sainsbury Laboratory, University of Cambridge, Cambridge, UK; Institute of Molecular Plant Sciences, School of Biological Sciences, University of Edinburgh, Edinburgh EH9 3BF, UK; Bordeaux Imaging Centre, Plant Imaging Platform, UMS 3420, INRA-CNRS-INSERM-University of Bordeaux, Villenave-d’Ornon, France; Department of Plant Biotechnology and Bioinformatics, Ghent University, 9052 Ghent, Belgium; Center for Plant Systems Biology, VIB, 9052 Ghent, Belgium; Department of Plant Sciences, University of Geneva, 1211 Geneva, Switzerland; Laboratory for Biological Geochemistry, School of Architecture, Civil and Environmental Engineering, Ecole Polytechnique Fédérale de Lausanne (EPFL), CH-1015, Lausanne, Switzerland; Univ. Bordeaux, Bordeaux Proteome, Bordeaux, France; Biomedical Sciences Research Complex, University of St Andrews, Fife KY16 9ST, UK; Cell and Molecular Sciences, The James Hutton Institute, Dundee DD2 5DA, UK; Institute of Biotechnology, HiLIFE/Organismal and Evolutionary Biology Research Programme, Faculty of Biological and Environmental Sciences, Viikki Plant Science Centre, University of Helsinki, Helsinki, Finland

## Abstract

Membrane contact sites (MCS) are fundamental for intracellular communication, but their role in intercellular communication remains unexplored. We show that in plants, plasmodesmata communication bridges function as atypical endoplasmic reticulum (ER)-plasma membrane (PM) tubular MCS, operating at cell-cell interfaces. Similar to other MCS, ER-PM apposition is controlled by a protein-lipid tethering complex, but uniquely, this serves intercellular communication. Combining high-resolution microscopy, molecular dynamics, pharmacological and genetic approaches, we show that cell-cell trafficking is modulated through the combined action of Multiple C2 domains and transmembrane domain proteins (MCTP) 3, 4, and 6 ER-PM tethers, and phosphatidylinositol-4-phosphate (PI4P) lipid. Graded PI4P amounts regulate MCTP docking to the PM, their plasmodesmata localization and cell-cell permeability. SAC7, an ER-localized PI4P-phosphatase, regulates MCTP4 accumulation at plasmodesmata and modulates cell-cell trafficking capacity in a cell-type specific manner. Our findings expand MCS’s functions in information transmission, from intracellular to intercellular cellular activities.

**In brief:** Plant intercellular communication is regulated via tubular membrane contact through PI4P binding-ER-PM tether MCTP proteins

**Highlights:**

1. Plasmodesmata are unconventional ER/PM tubular contact sites located at cell-cell interface
2. Plasmodesmata operate as control valves, modulating ER-PM contacts to regulate transport
3. MCTP3, MCTP4, MCTP6 and PI4P tethering elements act as valve regulators
4. SAC7 PI4P phosphatase controls plasmodesmata MCS in a cell-type-specific manner

## INTRODUCTION

Recent years have seen a paradigm shift in our understanding of intracellular communication with the discovery of membrane contact sites (MCS) (Scorrano et al., 2019; Prinz et al., 2020). By mediating close contacts between two membranes, MCS act as specialized platforms for inter-organellar exchanges, enabling transfer of ions and other molecules (Soboloff et al., 2006; Mesmin et al., 2013; Schauder et al., 2014; Chung et al., 2015; von Filseck et al., 2015; Eden et al., 2016; Wong et al., 2016; Wilhelm et al., 2017; Balla, 2018; Bian et al., 2018; Kumagai and Hanada, 2019; Ruiz-Lopez et al., 2021; Qian et al., 2022; Radulovic et al., 2022; Guillén-Samander and De Camilli, 2023; Naón et al., 2023; Wozny et al., 2023). This direct molecular transfer between organelles involves specialized protein machinery and membrane proximity regulation (Scorrano et al., 2019). Interest in MCS has grown dramatically in the last decade, with a shift of focus from describing their molecular composition to understanding their functions. Besides inter-organellar communication, MCS were shown to be involved in organelle biogenesis, fission and trafficking, cargo sorting, cellular stress responses, and autophagy; highlighting the functional diversity of these eukaryotic structures (Eden et al., 2010; Friedman et al., 2011; Hamasaki et al., 2013; Rowland et al., 2014; Pérez-Sancho et al., 2015a; Dong et al., 2016; Hirabayashi et al., 2017; Lees et al., 2017; Collado et al., 2019; Hernández-Alvarez et al., 2019; Hoffmann et al., 2019; Lee et al., 2019a; Wang et al., 2019; Ruiz-Lopez et al., 2021). It is now clear that every organelle inside a single cell forms functional contacts, and that MCS activities are not only broad but also vital for the cell. While MCS are now well-established and essential contributors to intracellular coordination, many multicellular eukaryotic organisms require effective communication that extends beyond organelles and includes communication between individual cells. So far, our understanding of MCS function is restricted to intracellular activities, and there is currently no evidence supporting the notion that MCS facilitate direct molecular exchange between cells.

With this shift in mind, we addressed the possible functions of MCS in inter-cellular (as opposed to inter-organellar) molecular exchange. Among eukaryotic cells, the endoplasmic reticulum (ER) serves as a central hub for MCS, forming multiple contacts with various organelles(Salvador-Gallego et al., 2017; Wu et al., 2018a). In most eukaryotic cells the ER is confined to a single cell. However, in land plants and some green algae, the ER network extends from cell-to-cell through communication bridges known as plasmodesmata (Nicolas et al., 2017b) (**Figure 1A**). Unlike desmosome cell-cell junctions or any other cell-cell contact dependent communication structures, both the ER and the PM traverse cell-cell boundaries through plasmodesmata. Inside plasmodesmata the ER takes the form of a condensed membrane tube known as the desmotubule which comes in close apposition with the PM, creating an atypical tubular MCS-like structure (Ding et al., 1992; Tilsner et al., 2016; Nicolas et al., 2017a). Millions of plasmodesmata bridges weave through the plant body, allowing direct molecular exchange of nutrients and signaling molecules in the space between the ER and the PM, called the cytoplasmic sleeve (Deinum et al., 2019) (**Figure 1A**). These structures are pivotal for plant intercellular communication, allowing cells to coordinate their activities, exchange resources, and respond to environmental and developmental signals in a synchronized manner (Nakajima et al., 2001; Guseman et al., 2010; Benitez-Alfonso et al., 2013; Faulkner et al., 2013; Stahl et al., 2013; Wang et al., 2013; Daum et al., 2014; Vaddepalli et al., 2014; Otero et al., 2016; Stahl and Faulkner, 2016; Gaudioso-Pedraza et al., 2018; Tylewicz et al., 2018; Miyashima et al., 2019; Song et al., 2019; Tran et al., 2019; Gao et al., 2020; Mellor et al., 2020; Kitagawa et al., 2022; Mehra et al., 2022; Tee et al., 2022; Wang et al., 2023). Yet the molecular mechanisms regulating cell-cell molecular flow continue to evade complete understanding.

**Figure 1.**
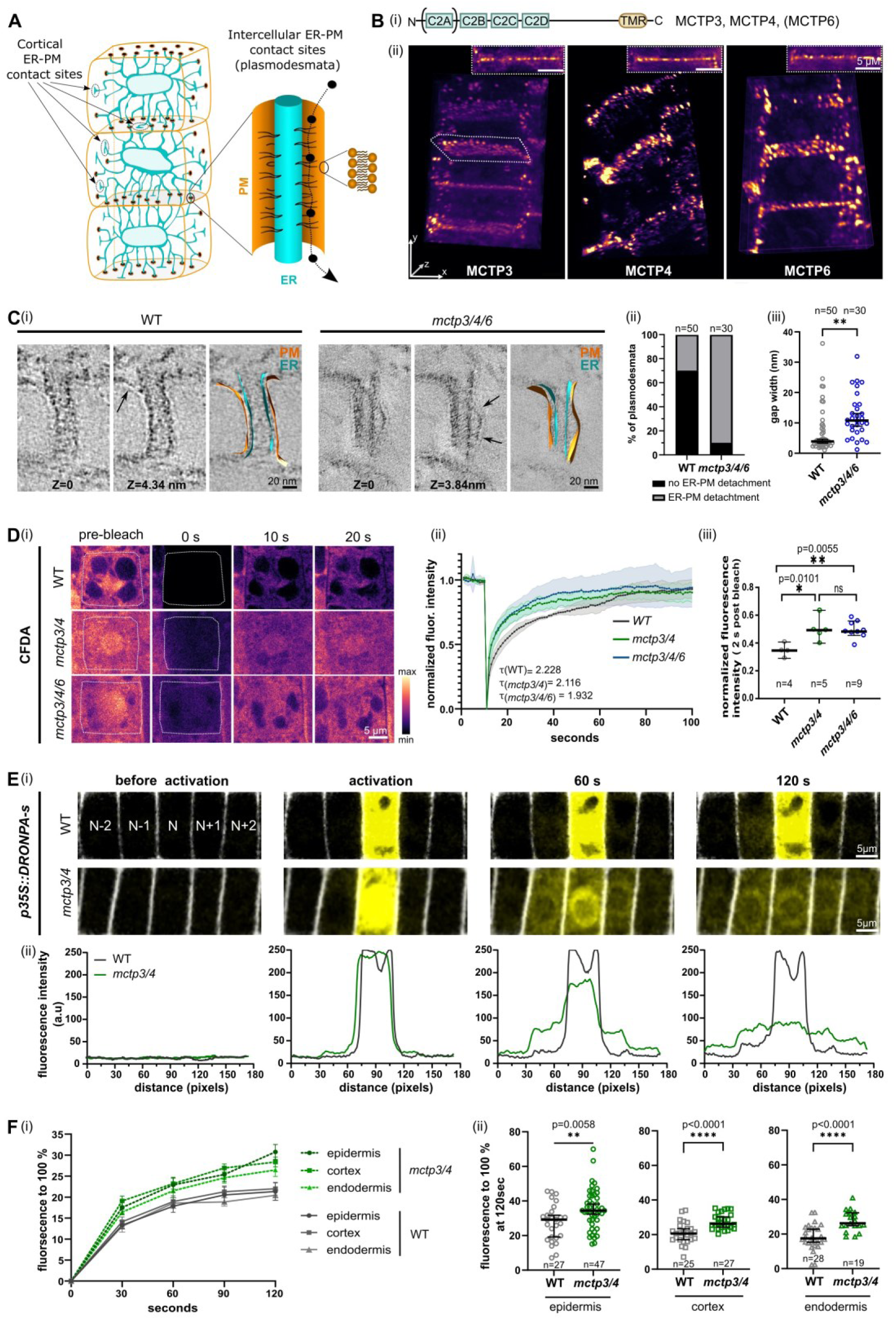
MCTPs function as ER-PM tethers within plasmodesmata and regulate its diffusion capacity. (A) Schematic representation of ER-PM contact sites in plants. (B) (i) Schematic representation of plant ER-PM contact site tethers MCTPs depicting phospholipid-binding C2 domains and ER-anchored transmembrane regions (TMR). The size of the domains and the distances between them is not to scale. (ii) 3D view of *mctp3/4 pUBQ10::eYFP MCTP3*, *mctp3/4 pUBQ10::eYFP-MCTP4* and *mctp3/4 pUBQ10::MCTP6-mVenus* acquired with lattice light-sheet microscopy. Insets represent one focal plane from the lattice light-sheet set. In this panel and throughout all the figures in the paper, *mctp3/4* stands for *mctp3-2/mctp4-1* and *mctp3/4/6* stands for *mctp3-1/mctp4-1/mctp6-1*. (C) (i) Individual tomographic slices and segmented 3D reconstructions of plasmodesmata in wild-type (WT) and *mctp3/4/6*. Arrows pointing to ER-PM detachment. Z are two different planes (ii) Contingency plot of the percentage of plasmodesmata showing ER-PM detachment. (iii) Quantification of the distance between ER and PM in WT and *mctp3/4/6* plasmodesmata at the point where the gap is the widest. (D) CFDA mobility assay measured by FRAP in root epidermal meristematic cells in WT, *mctp3/4* and *mctp3/4/6*. (i) Representative images before and the indicated time after bleach (the cells used for quantification are surrounded by dotted squares). (ii) Normalized fluorescence intensity over a period of 90 seconds after bleaching. Half-life (τ) of CFDA recovery for each genotype is shown. (iii) Normalized fluorescence intensity 2 seconds after photobleaching. Statistical analysis was done with ANOVA followed by Tukey’s test. Experiment was repeated twice with similar results. (E-F) DRONPA-s mobility assay measured as the diffusion of DRONPA-s signal from one single activated cell by two-photon into the neighboring cells in the longitudinal axis. (E) (i) Representative images of the DRONPA-s signal in WT and *mctp3/4* before activation, at activation and 60 and 120 second post activation. N, activated cell; N±1, direct neighbors; N±2, second level neighbors. Propidium iodide (PI, white) was used to mark cell peripheries. (ii) Signal plot showing raw intensity values from the images in (i). F (i) Normalized fluorescence over time of DRONPA-s averaged from N±1 in meristematic epidermis, cortex and endodermis cells imaged every 15 seconds. Lines indicate mean. Error bars indicate SEM. (ii) Percentage of DRONPA-s normalized fluorescence in the N±1 cells at 120 seconds post activation from (i). At least 6 roots were used per cell type per genotype per experiment. Experiment was repeated 3 times and results were pooled for quantification. When not differently stated, statistical analysis was done with Student t-text. Lines indicate median. Error bars indicate 95% confidence interval (CI).

While plasmodesmata comply with the general definition of MCS: (1) area of close proximity between two membranes, (2) absence of membrane fusion, (3) site of specialized function and (4) defined proteome and lipidome (Li et al., 2021), we still do not know if they function as MCS. Their unique membrane organization has been known for decades and yet it is still not known whether plasmodesmata-driven cell-cell communication requires regulation of ER-PM contacts and if dysregulation of these contacts impacts the functionality of plasmodesmata. Until now, the local synthesis and degradation of a beta-1,3-glucan polymer (callose) in the wall around plasmodesmata, is the only characterized mechanism to regulate plasmodesmata permeability (Zavaliev et al., 2011; Amsbury et al., 2017; Wu et al., 2018b). Additionally, plant viruses also encode movement proteins, which can also act as exogenous regulators of plasmodesmata permeability to transfer the viral genome through plasmodesmata (Crawford and Zambryski, 2001; Guenoune-Gelbart et al., 2008; Amari et al., 2010; Tilsner et al., 2013; Tilsner et al., 2014).

Given their position at cell-to-cell interfaces, we hypothesize that regulating ER-PM contact sites at plasmodesmata may function as a mechanism to modulate the diffusion of cytoplasmic components between neighboring cells. This mechanism would stand as the first example in eucaryotes of MCS driven-intercellular communication. In this work, we undertook to identify the molecular actors by which plasmodesmata may control the exchange of information among cells by regulating MCS, with a focus on the model plant *Arabidopsis thaliana*.

## RESULTS

### MCTP3, MCTP4 and MCTP6 form a tethering complex at plasmodesmata and redundantly impact plant growth and development

Plasmodesmata are dynamic nanoscopic structures with the capacity to dilate and contract, acting as checkpoints controlling cell-to-cell trafficking. Sitting at the cell-cell interface, they have the potential to combine membrane contact regulation with intercellular functions. In this scenario, MCS would act as a valve, controlling molecular flow by adjusting the ER-PM gap and/or changing the cytoplasmic sleeve property through steric hindrance (**Figure 1A**). To test this hypothesis, we first targeted the molecular machinery controlling ER-PM contact. MCS are regulated by protein tethers which physically bridge the two membranes (Scorrano et al., 2019). Previously, we identified members of the Multiple C2 domains and transmembrane domain proteins (MCTPs) family as putative plasmodesmata tethers (Brault et al., 2019). Besides, Miras et al. (Miras et al., 2022b) identified MCTPs among 20 core plasmodesmata proteins by cross-referencing the proteomes of *Arabidopsis thaliana*, *Populus trichocarpa*, and *Nicotiana benthamiana*. Johnston *et al*. (Johnston et al., 2023) also classified them as one of three conserved plasmodesmata families through phyloproteomics.

We therefore selected MCTPs as ER-PM tether candidates and evaluated their function in cell-cell trafficking in the model plant *Arabidopsis thaliana*. MCTPs are structurally related to the MCS tethers Extended-synaptotagmins (E-Syts) in animal and Synaptotagmins (SYTs) in plant (Giordano et al., 2013; Pérez-Sancho et al., 2015a; Brault et al., 2019; Ruiz-Lopez et al., 2021). They consist of a C-terminal ER-anchor and three to four C2 lipid binding domains (**Figure 1B**). Nearly all plant cells are interconnected by plasmodesmata, so a primary component is expected to be widely expressed and consistently associated with them, regardless of tissue type or developmental stage. In Arabidopsis, the MCTP family comprises 16 members, among which MCTP3, MCTP4, and MCTP6 were identified in plasmodesmata fractions in a systematic manner (Fernandez-Calvino et al., 2011; Kraner et al., 2017; Brault et al., 2019; Kirk et al., 2022; Miras et al., 2022a; Gombos et al., 2023; Johnston et al., 2023). Analysis of publicly available gene expression profiling databases revealed that all three genes were widely and consistently expressed in both shoot and root tissues, with MCTP4 exhibiting the strongest expression levels (**Figure S1A and S1B**). In all examined tissues, the proteins displayed a dotty localization at cell-cell interfaces, reminiscent of plasmodesmata (**Figure 1B; Figure S1C-E; Movie S1-3**). All three members show plasmodesmata association when expressed under their endogenous promoters as confirmed by Airyscan imaging and co-localization with callose, a well-established plasmodesmata marker (**Figure S1F;** Brault et al. 2019). MCTP3, 4 and 6 can physically interact when co-expressed as shown by co-immunoprecipitation when transiently expressed (**Figure S2A-B)**. The interaction at plasmodesmata was confirmed using Förster Resonance Energy Transfer Fluorescence Lifetime Imaging Microscopy (FRET-FLIM) analysis with functional pMCTP4::TagRFP-MCTP4 co-expressed with pMCTP3::eYFP-MCTP3 in the complemented triple mutant background *mctp3-1/mctp4-1/mctp6-1* (**Figure S2C**).

Single loss-of-function mutants (*mctp3-1*, *mctp3-2, mctp4-1, mctp4-2* or *mctp6-1*) presented no strong phenotypic anomaly (**Figure S2D-E, S3)**. However, the double mutants involving MCTP4 loss-of-function (*mctp3-1/mctp4-1, mctp3-2/mctp4-1, mctp4-1/mctp6-1*) were severely affected in both shoot and root growth and development (**Figure S2D-E, S3**). The triple *mctp3-1/mctp4-1/mctp6-1* mutant was not viable when grown on soil and died after two weeks (**Figure S3)**. The strong pleiotropic phenotype is consistent with a role in the coordination of cellular activity at the multicellular level.

Altogether, these data indicate that MCTP3, MCTP4, and MCTP6 are core plasmodesmata constituents making them suitable targets for investigating plasmodesmata MCS function.

### MCTPs act as ER/PM tethers inside plasmodesmata bridges, controlling membrane contact

In previous work, we showed that overexpressing MCTP4 in a delta-ER-PM tether yeast mutant partially restored cortical ER, suggesting a potential function as membrane tether (Brault et al., 2019). To test whether MCTP3, MCTP4 and MCTP6 act as ER-PM linkers inside plasmodesmata, we checked whether their genetic deletion would loosen the ER-PM gap within the bridges.

Functional redundancy has been shown for E-Syts and SYTs (Saheki et al., 2016; Lee et al., 2020; Ruiz-Lopez et al., 2021) with only high order mutants showing clear ER-PM detachment (Giordano et al., 2013). Therefore, we focused on *mctp3-1/mctp4-1/mctp6-1* mutant to check for potential plasmodesmata structural defects. We used primary root, which allows high-resolution microscopy imaging of plasmodesmata by electron tomography on high-pressure frozen, freeze-substituted samples (Nicolas et al., 2017a; Yan et al., 2019). Electron tomography is the only technology capable of achieving the nanometer-scale resolution necessary to visualize plasmodesmata, which are only 20-40 nm in diameter. We focused on the root epidermis division zone, where plasmodesmata structure is well preserved after cryo-fixation, and on apical-basal walls where plasmodesmata are highly active in transport (Nicolas et al., 2017a). We adapted our previous protocol (Petit et al.; Nicolas et al., 2017a) to reduce staining and increase substructural resolution (see material and methods for details). This new protocol allowed us to visualize the ER-PM boundary within plasmodesmata and discern potential membrane detachment with more precision, while maintaining excellent cell fixation and structural membrane preservation (**Figure S4A**). Altogether we acquired over 80 tomograms of plasmodesmata. While in wild-type plants, a large proportion of plasmodesmata presented tight ER-PM contacts throughout the entire length of plasmodesmata (∼70 % plasmodesmata; n=50) as previously described (Nicolas et al., 2017a) (**Figure 1C, Movie S4),** this configuration was lower in the *mctp3-1/mctp4-1/mctp6-1* mutant (∼10 % plasmodesmata; n=30). Instead, the majority of plasmodesmata in the *mctp3-1/mctp4-1/mctp6-1* mutant showed clear and visible ER-PM membrane detachment (∼90% of plasmodesmata in *mctp3-1/mctp4-1/mctp6-1* mutant as opposed to ∼30% in the wild-type) (**Figure 1C (i) (ii), Figure S4B, Movies S5-6**). In some cases (about ∼10%), membrane separation in *mctp3-1/mctp4-1/mctp6-1* was nearly complete, as illustrated in **Figure S4B (iii)** (**Movie S6**). Furthermore, when measuring the ER-PM distance at detachment points, we observed larger values in the triple *mctp* mutant compared to wild-type (**Figure 1C (iii))**. Average plasmodesmata diameter (PM-PM) was slightly higher *mctp* mutant compared to wild-type (**Figure S4C**) likely due to more frequent ER-PM detachments. These data indicate that MCTP3/MCTP4/MCTP6 act as ER-PM tethers inside the plasmodesmata bridges and that their loss-of-function leads to looser membrane junctions.

### Loss of MCTP ER-PM tethers increases intercellular molecular flow through plasmodesmata

We next tested the impact of loss of MCTP tethers on intercellular trafficking. We first used fluorescence recovery after photobleaching (FRAP) on the carboxyfluorescein diacetate (CFDA) cytosolic dye. CFDA is a cell-permeant compound which once loaded in the cells is cleaved by intracellular esterases, creating a membrane-impermeable fluorescent probe that moves from cell-to-cell through plasmodesmata (Nicolas et al., 2017a). After photobleaching in the root division zone, we noted a significantly faster recovery in both *mctp3-2/mctp4-1* and *mctp3-1/mctp4-1/mctp6-1* mutants compared to wild-type (**Figure 1D**). No differences were noted in terms of recovery rate between the *mctp3-2/mctp4-1* and *mctp3-1/mctp4-1/mctp6-1* mutants. This could be attributed to MCTP6 down regulation in the *mctp3-2/mctp4-1* mutant background (Brault et al., 2019).To validate that loose ER-PM contacts within plasmodesmata is indeed leading to an increase in molecular flow, we next employed the genetically encoded DRONPA-s, a reversibly switchable photoactivatable fluorescent marker. DRONPA-s is bigger than CFDA with an estimated Stoke radius of 2.8 nm against 0.61 nm for CFDA, allowing us to assess how molecules of different sizes behave. Cytosolic DRONPA-s was selectively activated in a single cell via two-photon excitation microscopy, and movement into neighboring cells monitored over time (Gerlitz et al., 2018). This enabled us to precisely evaluate variations in molecular flow in the apico-basal walls of the root epidermis division zone, where the tomography study was conducted. We selected both the double *mctp3-2/mctp4-1* and the triple *mctp3-1/mctp4-1/mctp6-1* mutants to express cytosolic DRONPA-s. Despite repetitive attempts, we only obtained sufficient DRONPA-s expression levels in the double *mctp3-2/mctp4-1* mutant. After activation of DRONPA-s in one cell (N), its cell-cell transport can be monitored as the loss of DRONPA-s signal in the activated cell or the gain of DRONPA-s in the neighbors (N+1, N+2 and N-1, N-2) over time (**Figure 1E-F**). Loss-of-function in *mctp3-2/mctp4-1* resulted in much faster DRONPA-s loss in the activated cell and significantly increased movement in the N±1 neighboring cells (**Figure 1E-F**; **Movie S7**), indicating that plasmodesmata efflux capacity was more permissive than in wild-type plants. This permissive molecular flow was not specific to the epidermis, as both the endodermis and cortex presented a similar pattern (**Figure 1F**). Together these data show that modifying the ER-PM contacts by means of MCTP3/MCTP4/MCTP6 tether complex, modifies plasmodesmata transport capacity.

### Heightened cell-to-cell diffusion in *mctp* mutants is independent of callose deposition

In the text-book model, plasmodesmata cell-cell trafficking is primarily regulated by callose, a beta 1-3 extracellular glucan that is specifically enriched around plasmodesmata and metabolized by plasmodesmata-located enzymes. Callose accumulation reduces cell-cell transport while its degradation increases transport (Amsbury et al., 2017). While the exact mechanism of callose action remains elusive, it is proposed to modulate the size of the cytoplasmic sleeve, by increasing or relaxing pressure on the PM.

We therefore enquired if the enhanced cell-cell molecular flow in *mctp3-2/mctp4-1* and *mctp3-1/mctp4-1/mctp6-1* mutants was caused by decreased callose levels. We first checked if the expression of genes associated with callose regulation at plasmodesmata was modified by performing RNAseq analysis of root extracts. The data indicate that none of the callose-related proteins were mis-regulated in either the single *mctp3-2* or *mctp4-1* mutants, or in the *mctp3-2/mctp4-1* double mutant when compared to wild-type plants (**Figure 2A; Figure S5**). We then performed comparative proteomic analysis to see if callose-related proteins were mis-regulated at the protein level. Total protein extract from wild-type, double and triple *mctp* mutant seedlings were simultaneously analyzed by liquid chromatography–tandem mass spectrometry (LC-MS/MS). For each protein identified, its relative enrichment or depletion in the *mctp* mutants versus wild-type was determined (see M&M for details). We selected the same set of proteins as for the callose-related RNAseq analysis (**Figure 2A**). We observed no significant changes in the callose-related proteins that were detected (**Figure 2B**). We next directly quantified callose levels *in situ* in the root division zone by whole-mount immunolocalization using callose-specific monoclonal antibodies, in the same area used for DRONPA-s cell-cell trafficking assay (**Figure 2C)**. The *mctp* double and triple mutants presented similar accumulation of callose as wild-type plants (**Figure 2D**). From these data, we concluded that the increase of cell-cell trafficking in *mctp* mutants is not caused by a reduction of callose levels.

**Figure 2.**
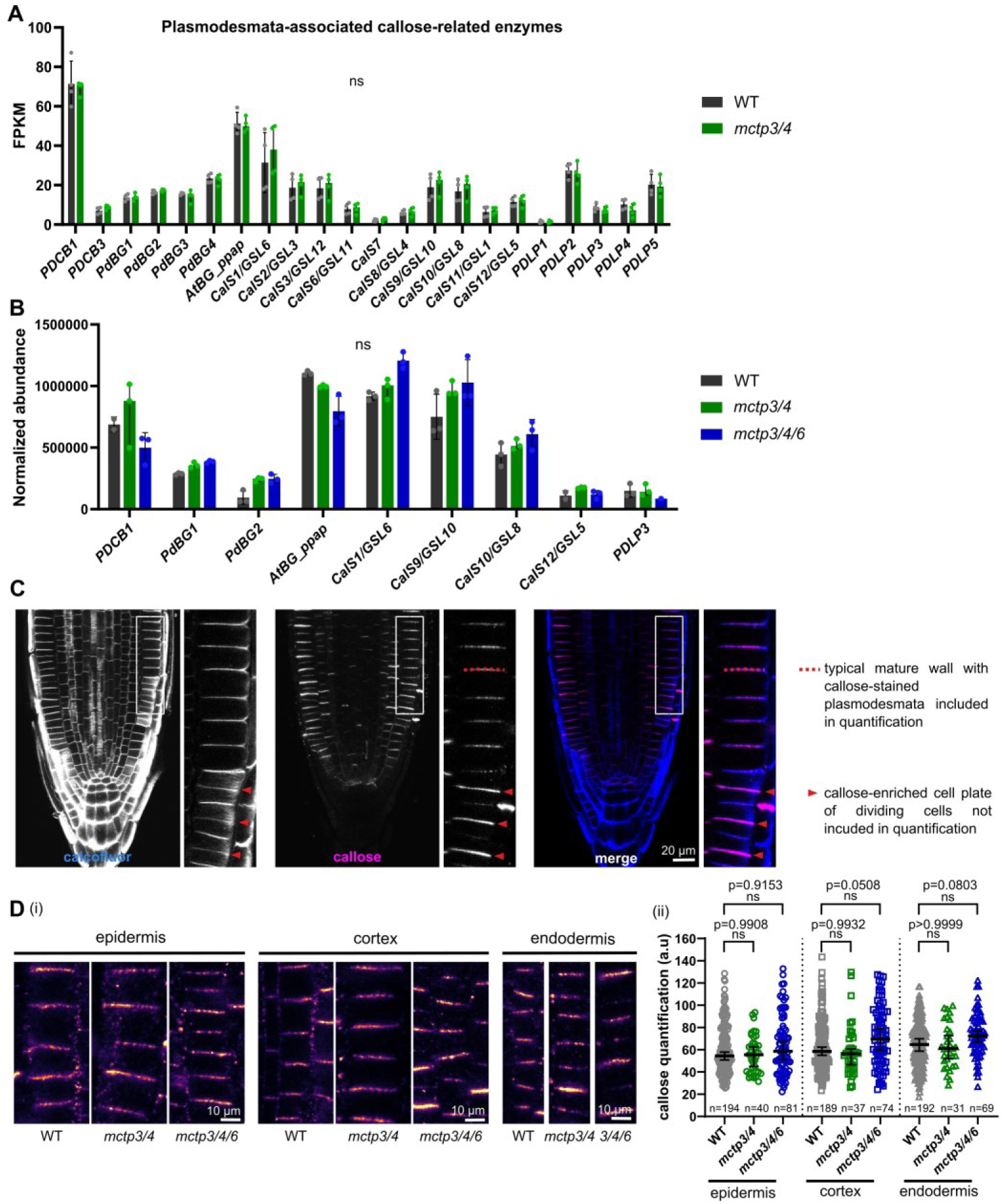
Loss of function of MCTP does not lead to defects in callose deposition. (A) Normalized expression of callose-related enzymes based on mRNA-seq of WT and *mctp3/4*. FPKM, Fragments per kilobase mapped. Bars reach the value of the median. Error bars represent 95% CI. Statistical analysis was done with multiple paired t-test. Experiment includes 4 biological replicates, each of them being a pool of 40 mg of 5 days old roots cut at 1 cm from the tip. (B) Normalized protein abundance of callose related enzymes based on mass spectrometry of WT, *mctp3/4* and *mctp3/4/6*. Bars reach the value of the median. Error bars represent 95 % CI. Statistical analysis was done with background-based ANOVA. Experiment includes 3 biological replicates, each of them being a pool of 80 mg of 5 days old roots cut at 1 cm from the tip. (C) Representative images of whole-mount anti-callose immunostaining (magenta) of WT showing the root apical meristem. Calcofluor (blue) was used to stain cell-walls. The white rectangle indicates the area generally used for quantification. For each mature cell-wall, a segmented line was drawn (dashed-red line as an example) and the mean intensity value along the line was used for quantification. Red arrowheads point to cell-walls that are not included in the analysis because they are just formed, as indicated by a very intense anti-callose signal and weak calcofluor signal. (D) Callose accumulation in the epidermis, cortex and endodermis meristematic cells of WT, *mctp3/4* and *mctp3/4/6*. (i) Representative confocal images of anti-callose immunostaining. (ii) Quantification of callose as explained in (C). Lines indicate median. Error bars indicate 95 % CI. Statistical analysis was done with ANOVA followed by Tukey’s test. A minimum of 4 roots were used per genotype, per experiment. Experiment was repeated twice with similar results and data were pooled together.

### Defective ER-PM tethering impairs the dynamic control of intercellular molecular flow

Intercellular communication is dynamically controlled, and a key feature of plasmodesmata lies in their ability to respond to various environmental or developmental stimuli by reducing or increasing intercellular molecular flow (Gisel et al., 1999; Faulkner et al., 2013; Han et al., 2014; Cui and Lee, 2016; Cheval and Faulkner, 2018; Gaudioso-Pedraza et al., 2018; Lexy et al., 2018; Tylewicz et al., 2018; Grison et al., 2019; Sager et al., 2020; Mehra et al., 2022; Tee et al., 2022; Wang et al., 2023). We therefore tested whether dynamic regulation of trafficking persisted in the absence of a fully functional tether machinery.

We first tested how plasmodesmata responded to brassinosteroid levels, which are known to modify cell-cell transport and callose deposition (Wang et al., 2023). We used brassinazole, a brassinosteroid biosynthesis inhibitor, which drastically reduces callose levels and promotes cell-cell trafficking (Wang et al., 2023). Upon exposure to brassinazole (24h, 1 µM), wild-type roots exhibited almost complete suppression of callose accumulation (**Figure 3A**). A similar effect was observed in both *mctp3-2/mctp4-1* and *mctp3-1/mctp4-1/mctp6-1* mutants, reaffirming that callose response remains unaffected in the *mctp* mutants (**Figure 3A**). As expected, in wild-type plants, DRONPA-s spread faster into neighboring cells after callose removal (**Figure 3B**). In this condition, cell-cell trafficking in the wild-type increased to the same magnitude as in untreated *mctp3-2/mctp4-1* plants. However, callose removal in the *mctp3-2/mctp4-1* mutant did not change macromolecular flux, which remained identical to control conditions (**Figure 3B**).

**Figure 3.**
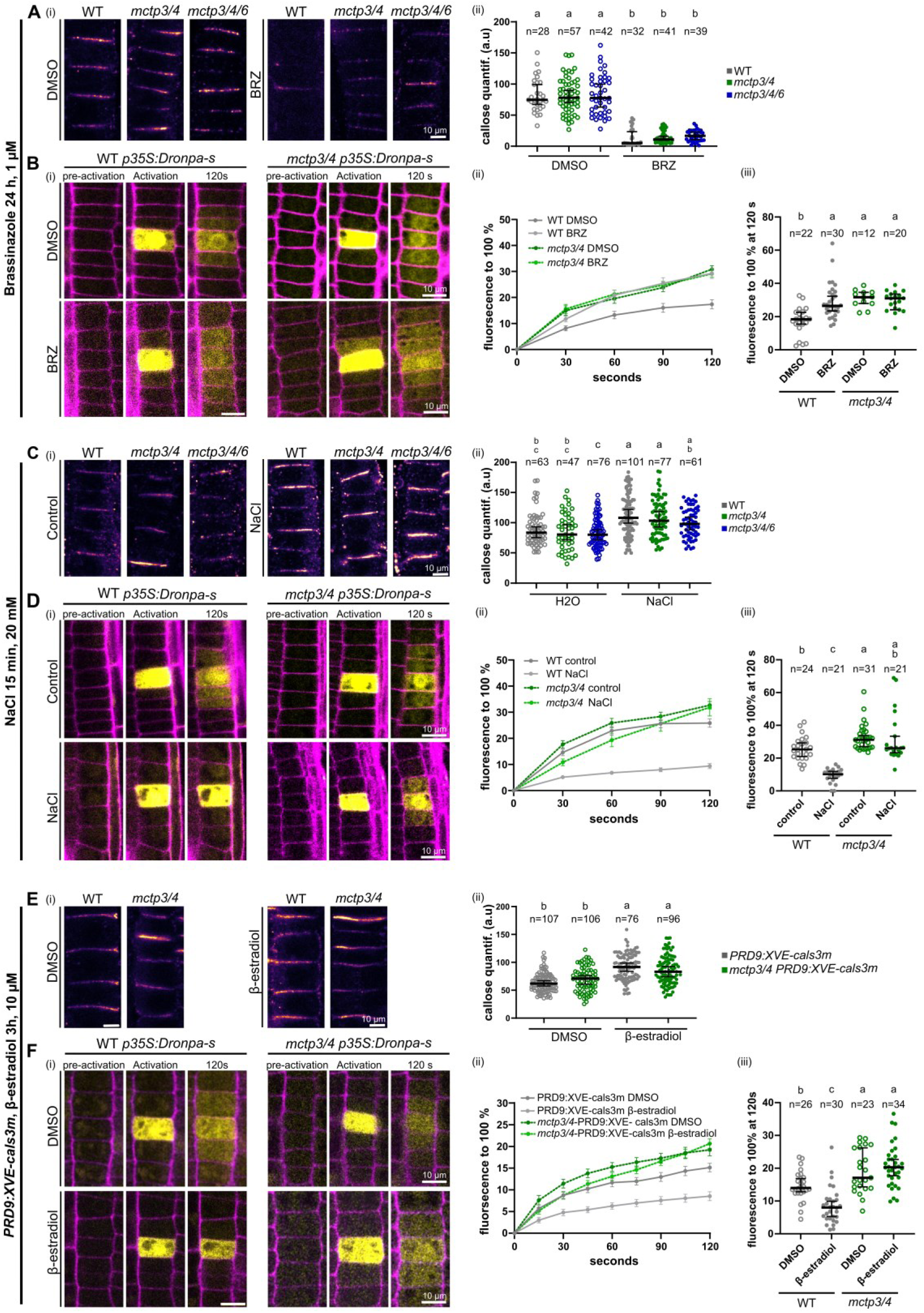
Callose regulation in loss of function MCTPs has minor effects on the cell-cell diffusion. (A-B) Brassinazole treatment; DMSO or 1 μM Brassinazole (BRZ) for 24 h. (A) Anti-callose immunostaining in treated WT, *mctp3/4* and *mctp3/4/6*. (i). Representative confocal images of callose immunostaining. (ii) Quantification of callose. (B) DRONPA-s mobility assay in treated WT and *mctp3/4*. (i) Representative images of DRONPA-s diffusion. (ii) Normalized DRONPA-s fluorescence over time. Line show mean. Error bars show SEM. (iii). Percentage of DRONPA-s normalized fluorescence in the N±1 cells at 120 s post activation from (ii). (C-D) Salt short-term treatment; H2O (control) or 20mM NaCl for 15 min. (C) Anti-callose immunostaining in treated WT, *mctp3/4* and *mctp3/4/6* (i) Representative confocal images of callose immunostaining. (ii) Quantification of callose. (D) DRONPA-s mobility assay in treated WT and *mctp3/4*. (i) Representative images of DRONPA-s diffusion. (ii) Normalized DONRPA-s fluorescence over time. Line show mean. Error bars show SEM. (iii) Mean percentage of DRONPA-s normalized fluorescence in the N±1 cells at 120 s post activation from (ii). (E-F) Callose-inducible *pPRD9::XVE-cals3m*; DMSO or 10μM b-estradiol for 3 h. (E) Anti-callose immunostaining in treated WT *pPRD9::XVE-cals3m* and *mctp3/4 pPRD9::XVE-cals3m*. (i) Representative confocal images of callose immunostaining. (ii). Quantification of callose. (F) DRONPA-s mobility assay in treated WT *pPRD9::XVE-cals3m* and *mctp3/4 pPRD9::XVE-cals3m*. (i) Representative images of DRONPA-s diffusion. (ii) Normalized DRONPA-s fluorescence over time. Line show mean. Error bars show SEM. (iii) Percentage of DRONPA-s normalized fluorescence in the N±1 cells at 120 s post activation from (ii). When not specified differently, lines indicate median. Error bars indicate 95 % CI. Statistical analysis was done with ANOVA followed by Tukey’s test. Different letters indicate p<0.05. A minimum of 5 roots were used per genotype, per condition, per experiment. Experiments were repeated three times with similar results and data were pooled together.

Next, we tested if down-regulation of cell-cell trafficking was functional without MCTP tethers, by examining *mctp* mutant responses in conditions known to induce plasmodesmata closure by callose deposition. First, we used brassinolide (Wang et al., 2023) and we confirmed the closure of plasmodesmata upon brassinolide (24h, 200 nM) exposure in wild-type plants, as evidenced by a reduction in DRONPA-s cell-cell diffusion, and callose accumulation (**Figure S6A-B**). In this condition, *mctp* mutants showed an increase in callose identical to the wild-type (**Figure S6B**). However, the impact on cell-cell trafficking was minor and DRONPA-s diffusion remained to the same permissive level in brassinolide-treated *mctp4-1/mctp3-2* mutant than in mock-treated wild-type plants (**Figure S6A)**. Please note that we observed no obvious differences in MCTP4 localisation pattern at cell-cell interface upon brassinosteroid treatments (**Figure S6C).** We next used salt stress (15 min, 20 mM), which triggers short-term callose deposition and plasmodesmata closure (Hunter et al., 2019), as confirmed by quantitative callose immunolocalization and DRONPA-s diffusion (**Figure 3C-D**). Callose deposition was triggered to the same levels in *mctp3-2/mctp4-1, mctp3-2/mctp4-1/mctp6-1* and wild-type roots (**Figure 3C**). However, while wild-type showed sharp decrease in DRONPA-s cell-cell trafficking, plasmodesmata closure was inhibited in *mctp3-2/mctp4-1* mutants (**Figure 3D**).

Collectively these data indicate that without the MCTP-tether machinery, callose cannot efficiently regulate plasmodesmata. To confirm this, we manipulated callose levels genetically by expressing the gain-of-function callose synthase 3 gene (*cals3m)* (Vatén et al., 2011) under an inducible expression system in the epidermal cells in *mctp3-2/mctp4-1* mutant expressing cytosolic DRONPA-s. After 3h of induction with β-estradiol, we observed a clear accumulation of callose in both the wild-type and *mctp3-2/mctp4-1* mutants (**Figure 3E)**. Cals3m-induced callose deposition translated into a drastic reduction in DRONPA-s cell-cell trafficking in the wild-type but not in the *mctp3-2/mctp4-1* mutant (**Figure 3F, Movie S8)**.

Altogether, these data indicate that 1) dynamic regulation of cell-cell molecular flow is lost in the absence of functional ER-PM tethers, and 2) callose regulation is no longer effective in *mctp* mutants.

### PI4P levels control MCTP docking to the PM

MCTPs are anchored into the ER through their C-terminal transmembrane region, and were proposed to dock to the PM by interacting with anionic phospholipids through their C2-lipid binding domains (Brault et al., 2019) akin to E-Syts and SYTs (Giordano et al., 2013; Pérez-Sancho et al., 2015a; Benavente et al., 2021) **(Movie S9**). Anionic phospholipids act as signals in membranes and they are involved in development, reproduction and environmental interactions. They are low-abundant lipids, whose concentration can drastically vary depending on the cell types or during stress episodes (Noack and Jaillais, 2020). We therefore wondered if anionic lipids could regulate plasmodesmata, by controlling MCTP interaction with the PM. We focused on PI4P, which was previously identified as a prime determinant of the PM inner leaflet electrostatic signature in plants (Simon et al., 2016a). Earlier studies on individual MCTP C2 domains suggested that PI4P may indeed promote docking to the PM (Brault et al., 2019). To confirm these data, we purified individual C2 domains of the master regulator MCTP4. While C2C and C2D were insoluble after expression in bacteria, we managed to purify C2B. We then performed liposome sedimentation assays with liposomes of different lipidic composition and confirmed preferential binding of MCTP4-C2B to the lipid bilayer in the presence of PI4P (**Figure 4A**). While PI4P is known to be present at the bulk PM, plasmodesmata, as any MCS, display a specialized membrane environment (Li et al., 2021) and it is not known whether they contain phosphoinositide species. We thus purified plasmodesmata from *A. thaliana* cell culture and ran LC-MS/MS analysis. Both PIP and PIP2 species were indeed detected in plasmodesmata fractions (**Figure 4B**).

**Figure 4.**
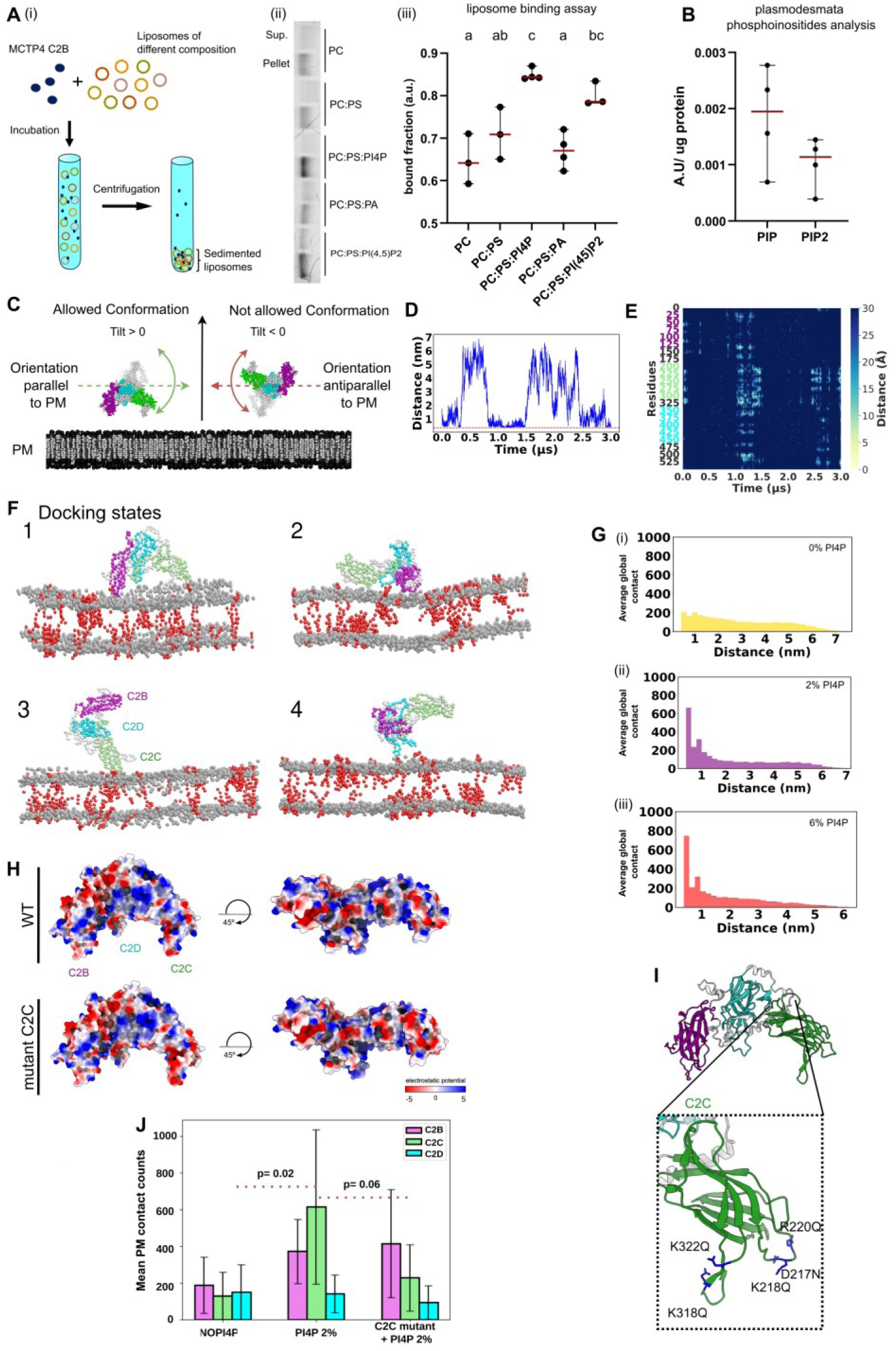
PI(4)P regulates the docking of MCTP4 to the PM by interacting with C2-lipid binding domains. (A) Liposome binding of the purified C2B domain of MCTP4. (i). Schematic representation of the liposome binding assay. Different colors indicate liposomes of different composition. (ii) SDS-PAGE of the supernatant (sup.) and precipitated (pellet) fractions after centrifugation. Proteins were visualized with a Bio-Rad stain-free imager and one single band was observed in each well, corresponding to the C2B domain of MCTP4. (iii) Quantification of the binding of C2B domain of MCTP4 to liposomes of different lipid compositions. Bound fraction was calculated as the ratio between C2B in the pellet and C2B in the supernatant. Line indicates median. Error bars indicate 95% CI. Statistical analysis was done using ANOVA followed by Tukey test. Different letters indicate p<0.05. Experiment was repeated 3-4 times and data were pooled together. PC: phosphatidyl choline; PS: phophatidyl serine; PI4P: phosphatidyl inositol 4-phosphate; PA: phosphatidic acid; PI(4,5)P2: phosphatidyl inositol 4,5-biphosphate. (B) Identification of phosphoinositides in plasmodesmata purified from WT cell cultures. Experiment includes 4 biological replicates. (C) Application of the Colvar Tilt variable to MCTP4. The harmonic potential activates when the tilt angle is below 0, constraining the system to maintain a realistic orientation relative to the membrane in the absence of the TMR/ER. (D) Analysis of the minimum distance between the C2 domains and the lipid bilayer over 3 microseconds, illustrating the dynamics of domain-membrane interactions from a single simulation. (E) Heatmap visualization representing the distances between C2 domain residues and PI4P in the membrane from the same simulation as in (C). The distances are color coded according to the legend in the right. (F) Snapshots from a simulation of the MCTP4 cytoplasmic domain engaging with a PM-like lipid bilayer in the presence of 2% PI4P: (1) Interaction through two C2 domains, (2) All three C2 domains simultaneously interacting with the bilayer, and (3 & 4) interaction with one different C2 domain. The simulations lasted 3 microseconds and were repeated 5 times for each condition. The C2 domains appear in different colors: C2B, magenta; C2C, green; C2D, cyan. (G) Histograms comparing the interaction frequency (average global contact) under three PI4P conditions: without PI4P, with 2% PI4P, and with 6% PI4P, showing increased frequency and stabilization of MCTP4-PM interactions correlating with higher PI4P concentrations. (H) Surface representation of the C2 block with the electrostatic potential ranging from -5 (red) to 5 (blue), created using ChimeraX. Two different angle-views are used to show the differences between WT block and C2 block with mutated amino acids on the C2C that are putatively responsible for binding with PI4P. (I) Schematic representation of the C2block (C2B in magenta, C2D in cyan) with a focus on C2C (in green) highlighting the amino acids that were mutated (in dark blue). (J) Average PM contact counts of the three separate C2blocks in conditions with no added PI4P, 2% PI4P and when putative lipid binding amino acids are mutated on the C2C block in 2% PI4P. Error bars show SD. Data provided by 7 simulations.

We next addressed the molecular mechanisms underlying PI4P’s role in relation to MCTP membrane tethering activity. We turned to molecular modelling and molecular dynamics to obtain near-atomistic structural insights into MCTP4’s interaction with the PM, with spatio-temporal resolution. In previous work, we only assessed individual C2 domains (Brault et al., 2019), here we investigated the behavior of the entire MCTP4 cytosolic domain (the full protein but without the C-terminal ER-anchored transmembrane region) in the presence of a PM-like lipid bilayer. We conducted molecular dynamics simulations using the recently introduced Martini 3 coarse-grained force field (Souza et al., 2021; Borges-Araújo et al., 2023) and we employed Alphafold2 for protein structure prediction (Jumper et al., 2021). MCTPs are anchored into the ER via their TMR domain, restricting their 3D orientation relative to the PM plane. In order to focus solely on functionally relevant protein positioning, we restricted movement in the MCTP4 C2 cytosolic domain, so that the C-terminus of the polypeptide (where the TMR normally is) could not face the PM (**Figure 4C**). The structural model for the cytoplasmic part of MCTP4, comprising the three C2 domains, was extracted from (Sritharan et al., 2023) and embedded in a water box with a lipid bilayer mimicking the PM composition, (∼70% phosphatidylcholine (PC), 20% Sitosterol, 10% phosphatidylserine (PS)) and 2% PI4P. To check the interactions between MCTP4 and the PM, we analyzed the fluctuating distances between the C2 domains and the lipid bilayer over time **(Figure 4D)** and the distribution of distances as a function of sequence (y axis) and time (x axis) (**Figure 4E).** Interactions with the PM involve multiple docking configurations, wherein combinations of C2/membrane interaction patterns differ, involving either one C2 (C2C **Figure 4F** state3 or C2D **Figure 4F** state4), two C2s (C2B and C2C) (**Figure 4F** state1), or all three (**Figure 4F** state2) see **Movie S10**. We next evaluated the interaction of the cytoplasmic part of MCTP4 with a PM with varying concentration of PI4P (0%, 2% or 6%). The data indicate that PI4P boosts the frequency of binding between the C2 domains and the PM, implying a role in stabilizing PM docking (**Figure 4G**).

We then identified the amino acids that were directly interacting with PI4P lipids. Examining the electrostatic profile of the C2 domains revealed several positively charged regions on their surfaces, mostly associated with lysines (shown in Movie S11 where the green border areas represent lysine residues) known to play a significant role in membrane binding of peripheral proteins (Tubiana et al., 2022). In our simulations with 2% PI4P, these residues showed significant interactions with the lipid bilayer (**Figure S7A**). We concentrated on the C2C domain which presented the highest hit targets and identified five residues (D217, K218, R220, K318, K322) which showed high PI4P-interaction frequency (**Figure 4I and S7B**) and mutated them to reduce their electrostatic potential (**Figure 4H and S7B**) and performed additional simulations. We conducted the simulations without PI4P and with 2% PI4P. As before, when C2C was not mutated, we observed increased protein-membrane contacts in the presence of PI4P (**Figure 4J**). However, after introducing mutations to the selected PI4P-interacting residues within the C2C domain, we observed a trend towards reduced membrane contacts of the C2C domain even in the presence of PI4P. **(Figure 4J)**.

Altogether, our data show that MCTP C2 domains conditionally interact with membranes depending on the concentration of PI4P, a lipid present at plasmodesmata. These findings suggest that PI4P, as a signaling lipid, may regulate the interaction of MCTP with the PM.

### MCS modulates cell-cell molecular flow through the tandem action of MCTP tethers and PI4P lipid

We next tested the impact of PI4P on cell-cell trafficking. Our data supports a scenario where MCTPs bind to the PM through interaction with PI4P. Thus, the removal of PI4P should phenocopy *mctp3-2/mctp4-1* in terms of cell-cell transport capacity. To test this hypothesis, we first assessed the effect of reduced PI4P levels by applying the PI4K inhibitor PAO (phenylarsine oxide) at a concentration and exposure time previously described to affect the localization of PI4P sensors in Arabidopsis roots (Simon et al., 2016) (**Figure 5A**). We then subjected PAO-treated wild-type plants expressing cytosolic DRONPA-s to our cell-cell connectivity test. Inhibiting PI4P synthesis in wild-type plants increased cell-cell trafficking to a similar level than non-treated *mctp3-2/mctp4-1* mutant (**Figure 5A**). By contrast, using YM201636 (an inhibitor of PI(3,5)P_2_ synthesis) or Wortmannin (an inhibitor of PI3P synthesis) did not change the cell-cell trafficking properties of plasmodesmata (**Figure S8A-C**). We then subjected *mctp3-2/mctp4-1 p*35S::DRONPA-s to the same PAO treatment. The mutant transport capacity was not affected by PAO (**Figure 5A**), consistent with the notion that MCTP and PI4P act together to regulate plasmodesmata. We also checked that neither PAO nor DMSO differentially affected callose levels in wild-type and *mctp3-2/mctp4-1* mutant backgrounds (**Figure 5B**).

**Figure 5:**
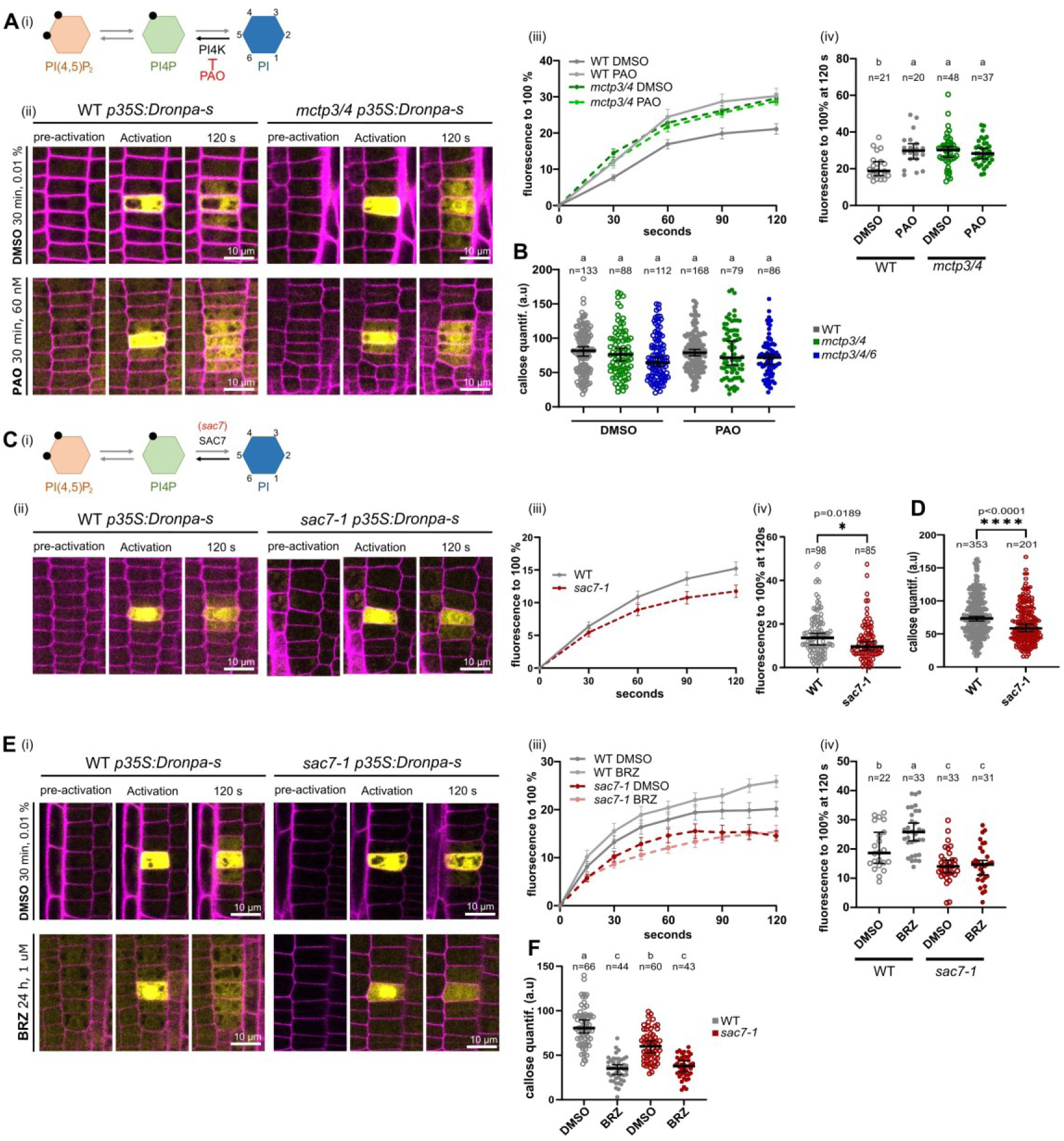
PI4P levels modulate cell-cell molecular flow. (A-B) PAO treatment; DMSO or PAO 60 nM for 30 min. (A) DRONPA-s diffusion assay in treated WT and *mctp3/4*. (i) Schema of phosphoinositide metabolism depicting the role of PAO blocking PI4K. (ii) Representative images of DRONPA-s diffusion. (iii) Normalized DONRPA-s fluorescence over time. Line indicates mean. Bars show SEM. (iv) Percentage of DRONPA-s normalized fluorescence in the N±1 cells at 120 seconds post activation from (iii). (B) Quantification of anti-callose immunostaining in WT, *mctp3/4* and *mctp3/4/6*. (C-D) *sac7-1* mutant. (C) DRONPA-s diffusion assay in WT and *sac7-1* meristematic epidermal cells. (i) Schema of phosphoinositide metabolism depicting the role of SAC7 as PI4P phosphatase. (ii) Representative images of DRONPA-s diffusion. (iii) Normalized DRONPA-s fluorescence over time. Line indicates mean. Bars show SEM. (iv) Percentage of DRONPA-s normalized fluorescence in the N±1 cells at 120 seconds post activation from (iii) Statistical analysis was done using Student t-test. (D) Quantification of anti-callose immunostaining in WT and *sac7-*Statistical analysis was done using Student t-test (E-F) *sac7-1* mutant, BRZ treatment; DMSO or 1 μM BRZ for 24 h (A) DRONPA-s mobility assay in treated WT and *sac7-1*. (i) Representative images of DRONPA-s diffusion. (ii) Normalized DRONPA-s fluorescence over time. Line show mean. Error bars show SEM. (iii) Percentage of DRONPA-s normalized fluorescence in the N±1 cells at 120 s post activation from (ii). (F) Quantification of anti-callose immunostaining in treated WT and *sac7-1*. When not differently stated, statistical analysis was done using ANOVA followed by Tukey’s test. Lines indicate median. Error bars indicate 95% CI. Different letters indicate p<0.05. A minimum of 5 plants per genotype, per condition, per experiment were used. Experiment were repeated 3 times and data were pooled together.

If PI4P levels are indeed instrumental to the regulation of cell-cell trafficking, elevated PI4P should have the opposite effect than low PI4P and induce plasmodesmata closure. Therefore, we next assayed if increased PI4P level modifies cell-cell transport. To do so, we generated a stable line expressing DRONPA-s in the *suppressor of actin7* (*sac7*, also known as *root hair defective4*) mutant. SAC7 is a PI4P 4-phosphatase, whose mutant accumulates PI4P *in vivo* (Thole et al., 2008; Song et al., 2021). The *sac7* mutant presented reduced DRONPA-s cell-to-cell diffusion, strengthening the idea that PI4P contributes to controlling plasmodesmata connectivity (**Figure 5C**). Compared to the wild-type, *sac7* has decreased accumulation of callose, ruling out callose as the cause of plasmodesmata closure **(Figure 5D**). We then asked whether callose removal could still open plasmodesmata with increased PI4P levels. Therefore, we treated *p*35S::DRONPA-s lines with brassinazole (24h, 1 µM), which led to callose removal in both wild-type and *sac7* backgrounds (**Figure 5F**). However, while wild-type roots responded by opening plasmodesmata with faster DRONPA-s movement, *sac7* maintained plasmodesmata closed (**Figure 5E**). Thus, the signaling lipid PI4P can modify cell-to-cell cytoplasmic diffusion bypassing callose accumulation, with reduced PI4P levels enhancing molecular movement, and elevated PI4P levels diminishing molecular flow.

Our data indicate that low PI4P accumulation results in plasmodesmata opening, while high PI4P levels lead to plasmodesmata closure. From these data we concluded that the signaling lipid PI4P may act as a regulator of plasmodesmata connectivity.

### SAC7 PI4P phosphatase is a cortical ER protein that closely associates with plasmodesmata in a cell type specific manner

Our data suggest that graded PI4P levels have the capacity to modulate plasmodesmata connectivity. We next addressed the mechanistic bases of this regulation and whether variations of PI4P levels are relevant to regulate plasmodesmata connectivity in a wild-type situation. To this end, we focused on the SAC7 enzyme, because i) the *sac7* mutant showed impaired plasmodesmata connectivity and ii) we found SAC7 in our plasmodesmata proteome (Brault et al., 2019).

To analyze SAC7 expression pattern and subcellular localization, we transformed the *sac7* mutant with p*SAC7::mCitrine-SAC7* and p*SAC7::2xmCherry-SAC7* constructs. Both fusion proteins complemented the short root hair phenotype of the *sac7* mutant showing that they were functional (**Figure S9 A-B**). Looking at SAC7 localization in root, we immediately noticed a distinct tissue-specific expression pattern, with SAC7 being predominantly expressed in the root hair cell file (trichoblast) while being present at much lower levels in non-root hair cells (atrichoblast) (**Figure 6A-B**). To confirm the strong expression of SAC7 in trichoblast cells, we crossed the *pSAC7::2xmCherry-SAC7* line with the *EXP7 SAND* reporter line, which is exclusively expressed in trichoblast cells (Marquès-Bueno et al., 2016). Colocalization revealed that 2xmCherry-SAC7 was indeed preferentially expressed in trichoblast, and was present in the meristem before the bulging and elongation of root hairs (**Figure 6B**).

**Figure 6:**
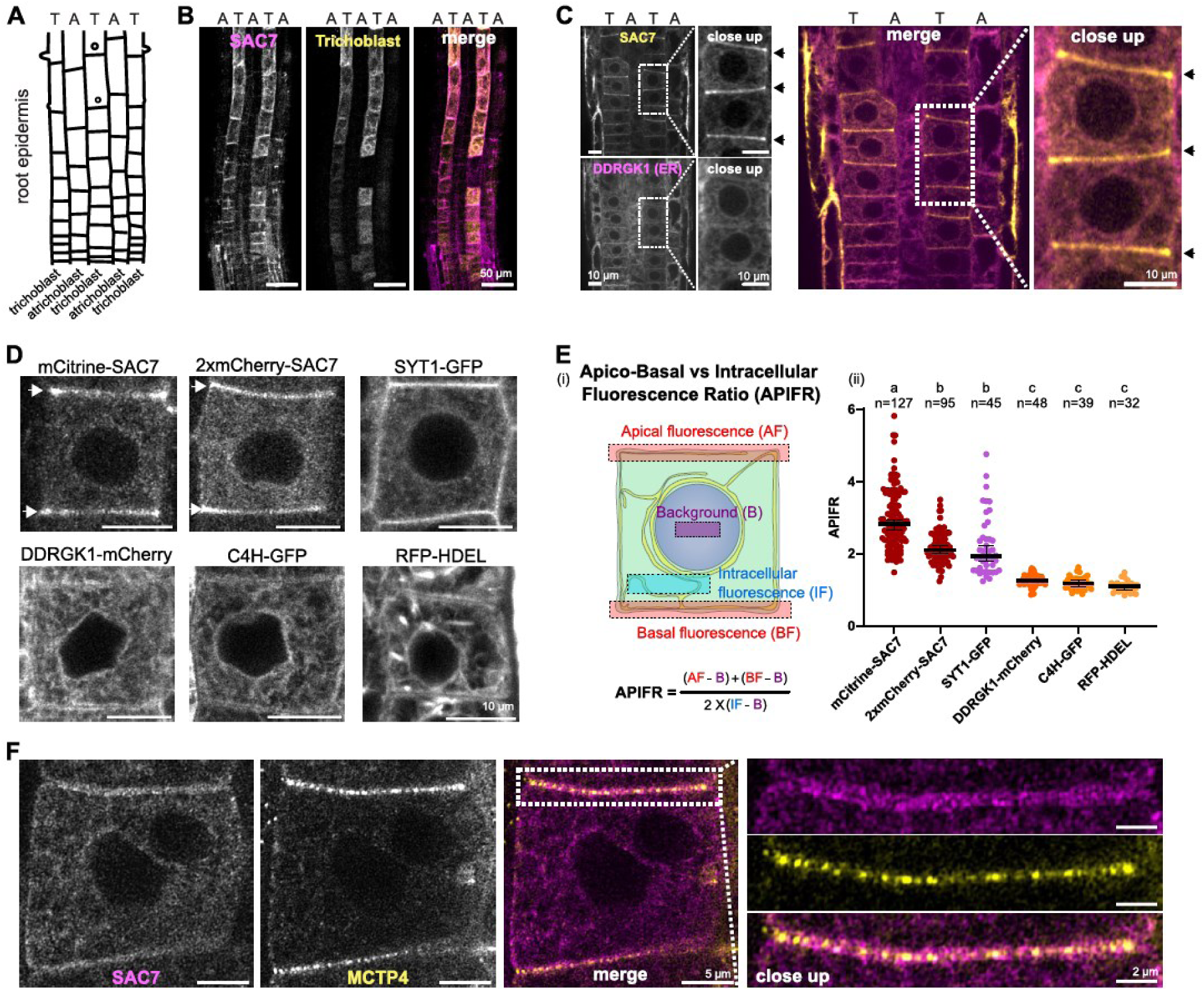
SAC7 is preferentially expressed in trichoblast cells and is polarly localized in the ER in proximity to MCTPs. (A) Schematic representation of the root epidermis showing the alternate pattern of root hair cells (trichoblasts, T) and non-root hair cells (atrichoblast, A). (B) Confocal micrographs showing the overlapping expression of *pSAC7::2xmCherry-SAC7* with *pEXP7::4xYFP* (trichoblast marker) in root-tip epidermis. (C) Confocal images showing the colocalization of *pSAC7::mCitrine-SAC7* (yellow) and *pUBQ10::DDRGK1-mCherry* (bulk ER marker, magenta) in the root meristem. Note the alternate accumulation in trichoblasts (T) but not in atrichoblasts (A). (D) Confocal images of the indicated ER-localized proteins in root meristematic cells. White arrows point toward apico-basal accumulation of SAC7 protein. (E) (i) Calculation of apical-basal vs intracellular fluorescent ratio (APIFR). (ii) Quantification APIFR of the proteins shown in (D). Statistical analysis was done with ANOVA followed by Tukey’s test. Lines indicate median. Error bars indicate 95% CI. (F) Confocal airyscan micrographs of roots coexpressing *pSAC7::2xmCherry-SAC7* (magenta) and *pMCTP4::MCTP4-GFP* (yellow) in root-tip epidermis cells, with a zoom of the apical-basal cell-cell interface.

We next analyzed the subcellular localization of *pSAC7::2xmCherry-SAC7.* The protein accumulated at the tip of growing root hairs, as previously reported (**Figure S9C**) (Thole et al., 2008). We also noticed that SAC7 localized in reticulated structures in root hairs that reminded us of the ER membranes. Because SAC7 was localized in the ER in BY–2 cells, and its yeast and animal orthologs, Sac1p/SAC1, are ER-resident proteins (Dubois and Jaillais, 2021), we explored whether SAC7 could locate in this compartment in Arabidopsis. To this end, we analyzed the localization of SAC7 in cotyledon epidermis, a cell type with a stereotypical reticulated ER, and confirmed that mCitrine-SAC7 colocalized with the ER marker DDRGK Domain-containing protein 1-mCherry (DDRGK1-mCherry) in this cell type (**Figure S9D**) (Stephani et al., 2020). In the root meristem, SAC7 also displayed an ER-localisation pattern while exhibiting a pronounced enrichment at the apico-basal cortical region of the cell (**Figure 6C**).

To gain further insights into this unique localization pattern, we compared the localization of SAC7 (mCitrine-SAC7 and 2xmCherry-SAC7) with various ER markers, including the cortical ER marker SYT1-GFP, the bulk ER membrane proteins DDRGK1-mCherry and Cinnamate-4-hydroxylase-GFP (C4H-GFP) and the ER luminal marker RFP-HDEL (**Figure 6D**). To quantify the cortical enrichment of SAC7, we used the Apico-basal vs Intracellular Fluorescence ratio (APIFR) across all ER proteins (**Figure 6E**). SYT1, an ER-PM tether (Pérez-Sancho et al., 2015b), exhibited an enrichment in the cortical ER similar to SAC7 (**Figure 6D-E**). However, unlike SAC7, SYT1 also accumulated at the lateral sides of atrichoblast cells (**Figure 6D**). All the other ER proteins displayed a more uniform distribution without significant enrichment at the cortical ER and a prominent labeling of the perinuclear ER (**Figure 6D-E**). Together, these results highlight a unique polar localization of SAC7 in the apical-basal cortical ER membranes.

As plasmodesmata were previously reported to be enriched at the apico-basal cell-cell interface as opposed to lateral sides in Arabidopsis root epidermis (Zhu et al., 1998). We therefore checked SAC7 association with MCTP4. Using high resolution airyscan microscopy, we colocalized 2xmCherry-SAC7 with GFP-MCTP4 both expressed under their native promoters. We noticed that 2xmCherry-SAC7 accumulated adjacent to GFP-MCTP4-labeled plasmodesmata (**Figure 6F**). Thus, SAC7 is an ER protein, enriched in the apico-basal cortical ER of trichoblast cells, in close association with plasmodesmata. Together, these data suggest that the SAC7 PI4P phosphatase have the potential to regulate PI4P levels at or near plasmodesmata.

### SAC7 is a cell-type specific regulator of MCTP localization and plasmodesmata connectivity

Our data indicate that PI4P levels regulate MCTP4 interaction with the PM and cell-cell connectivity. SAC7 is a PI4P phosphatase that localizes in proximity to MCTP4, and is differentially expressed in trichoblast/atrichoblast cells with high expression in the trichoblast cells (low PI4P) and low expression in the atrichoblast cells (high PI4P). We thus hypothesized that SAC7 might function as a negative regulator of MCTP and cell-to-cell connectivity. We first analyzed whether the localization of GFP-MCTP4 expressed under its native promoter differed between trichoblast and atrichoblast cells in wild-type plants. To this end, we counter-stained the cell wall using propidium iodine (PI) and automatically segmented the cell’s contour. We then quantified the GFP-MCTP4 signal at cell-cell junctions in trichoblast and atrichoblast cells (see **Figure 6E**). Using this pipeline, we found that MCTP4 accumulated at significant higher levels at the apical-basal interface in the atrichoblast cell files (which display low SAC7 expression levels) compared to trichoblast cells (which display high SAC7 expression levels) (**Figure 7A**).

**Figure 7:**
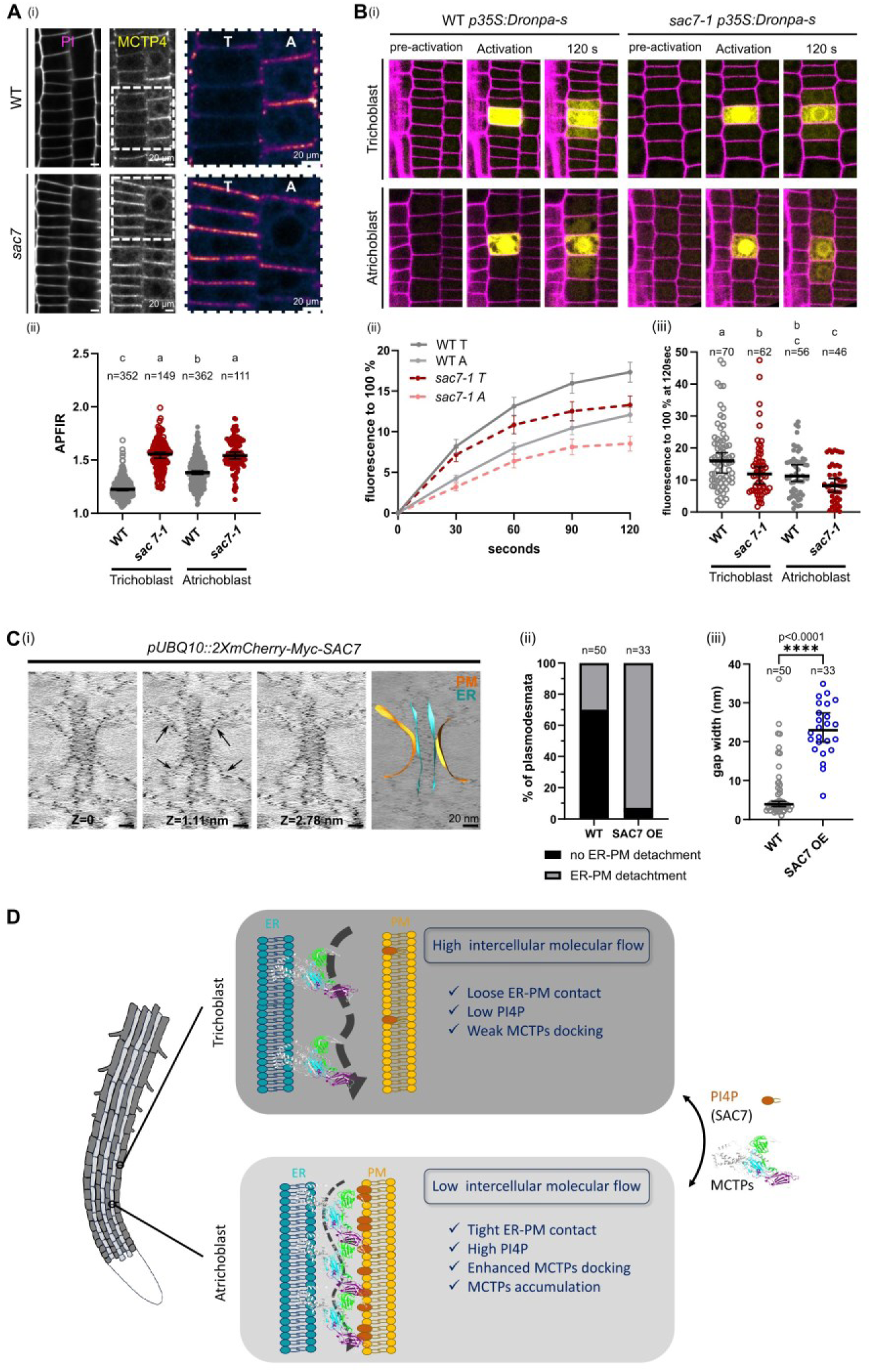
SAC7 regulates MCTP localization and cell-to-cell diffusion in a cell type-specific manner. (A) APIFR of *pMCTP4::MCTP4-GFP* in WT or *sac7-1* trichoblast (T) and atrichoblast (A) cells. (i) Representative confocal images. PI is used to visualized cell peripheries. (ii) APIFR quantification. Statistical analysis was done with ANOVA followed by Tukey’s test. Lines indicate median. Error bars indicate 95% CI. (B) DRONPA-s diffusion assay in WT and *sac7-1* trichoblast and atrichoblast meristematic cells. (i) Representative images of DRONPA-s diffusion. (ii) Normalized DRONPA-s fluorescence over time. Lines indicate mean. Bars show SEM. (iii) Percentage of DRONPA-s normalized fluorescence in the N±1 cells at 120 seconds post activation from (iii). Statistical analysis was done with ANOVA followed by Tukey’s test. Lines indicate median. Error bars indicate 95% CI. (C) (i) Individual tomographic slices and segmented 3D reconstruction of plasmodesmata in WT and *sac7-1*. Arrows pointing to ER-PM detachment. (ii) Contingency plot of the percentage of plasmodesmata showing ER-PM detachment. (iii) Quantification of the distance between ER and PM in WT and *sac7-1* plasmodesmata at the point where the gap is the widest. Lines show median. Error bars show 95 % CI. Please, note that WT values in graphs (ii) and (iii) are the same as in Fig 1C (ii) and (iii). (D) Cell-type dependent dynamic regulation of plasmodesmata MCS and its impact on molecular traffic

We next investigated how the differential localization of GFP-MCTP4 in trichoblast and atrichoblast cells affects cell-to-cell connectivity within these two cell files in wild-type plants. For that, we specifically quantified the dynamics of DRONPA-s diffusion in each cell file. We found that DRONPA-s diffused more rapidly in trichoblast than in atrichoblast cells, correlating with the high expression of SAC7 in this cell type and lower MCTP4 accumulation (**Figure 7B**). We then wondered if eliminating SAC7’s differential cell type expression would impact MCTP4 localization and thereby cell-cell connectivity. To this end, we expressed both GFP-MCTP4 (under its native promoter) and cytosolic DRONPA-s in *sac7-1* mutant background. We observed a strong increase of GFP-MCTP4 signal at the apico-basal walls of trichoblast cells in *sac7*. We also found a slight, but significant, increase of GFP-MCTP4 in *sac7* atrichoblast cells compared to wild-type, likely due to the weaker expression of SAC7 in this cell type. These changes in MCTP4 localization correlated with a modification of cell-cell molecular flow, which was diminished in the *sac7-1* mutant compared to the wild-type with the most significant impact in the trichoblast cell file (**Figure 7B**).

We then asked how low PI4P levels would impact plasmodesmata structure. To investigate this, we returned to electron tomography on high pressure frozen freeze substituted root samples. Because it is difficult to unambiguously distinguish between trichoblast and atrichoblast cells without cell type markers, we generated SAC7 over-expressing lines (p*UBQ10::2XmCherry-Myc-SAC7*). We screened through the apico-basal walls in the epidermis division zone (as for **Figure1 A**). We found that plasmodesmata in SAC7 overexpressing lines presented clear and frequent ER-PM detachment events (∼96 % plasmodesmata; n=23) (**Figure7C; Movie S12)** similar to *mctp* triple mutants (**Figure 1C**).

Our results suggest that the graded expression of SAC7 impacts MCTP4 localization in a cell type dependent manner, thereby directly regulating plasmodesmata connectivity. As a result, at the root tip, two cell types of the wild-type epidermis show different degrees of cell-to-cell molecular diffusion. Trichoblast and atrichoblast cells have different elongation rates, physiological status, interactions with the underlying cortex and overall have specific functions in root biology. Although the physiological impact of the differential connectivity of these two cell files remains to be determined, our data serve as a proof-of-concept that an ER/PM crosstalk, via the developmental modulation of PI4P/MCTP interactions, can regulate plasmodesmata connectivity.

## DISCUSSION

Our results uncover two important findings. Firstly, we reveal that while MCS are typically linked to inter-organellar communication in eukaryotes, they can also operate at cell-cell interface to govern inter-cellular communication. Plasmodesmata bridges are prevalent across the Plantae kingdom. Here, we show that they function as unconventional tubular ER-PM MCS regulating direct molecular exchange between neighboring cells. Similar to other MCS, the regulation of membrane contacts at plasmodesmata involves a complex molecular partnership consisting of phosphoinositides and protein tethers. However, in contrast to the typical MCS, which primarily govern direct molecular exchange between organelles, plasmodesmata MCS serve to control the flow of molecules between adjacent cells (**Figure 7D**). To our knowledge, this marks the first discovery of the functional involvement of MCS in direct intercellular exchange and hints at a new conceptual framework. Recent findings support the notion that MCS may also function in cell-cell communication in animals. A recent preprint reports two ER-PM MCS-associated proteins, TMEM24 and C2CD2, to be involved in cell-cell contact interaction in neurons (Johnson et al., 2023), although experimental support for effective communication is still lacking. Likewise, Bharathan et al. described close contact between the ER and desmosomal intercellular junctions (Bharathan et al., 2023). This work has sparked inquiries into whether an integrated ER-keratin-desmosome contact zone could facilitate the transmission of ER stress signals between cells. Our study brings this concept one step further and demonstrates the functional involvement of multicellular eucaryotic MCS in direct intercellular exchanges. The second important outcome of this study is to propose a novel regulatory mechanism for cell-cell trafficking underscoring the significance of MCS. Right now, the textbook model depicts callose as the master regulator of plasmodesmata. Unquestionably, numerous studies report the involvement of callose in opening or closing plasmodesmata in a wide range of developmental and physiological responses. The prevailing concept behind callose action is that callose build-up would create pressure on the PM, causing it to push against the ER, therefore obstructing cellular traffic by reducing the size of the cytoplasmic sleeve. However, recent research in the field suggests that callose actually functions as a plasticizer, enhancing wall flexibility, rather than forming a rigid donut-shaped structure that presses against the PM (Abou-Saleh et al., 2018). Moreover, extreme proximity between the ER and the PM, which can be as little as 3-4 nm (Nicolas et al., 2017a), is likely to require membrane bridging-complexes to induce and to stabilize the contact. This work suggests that without the molecular machinery acting at the ER-PM interface (*i.e.* PI4P and MCTP tethers) dynamic regulation of plasmodesmata is compromised even if callose regulation is still functional. Collectively our data indicate that MCTP/PI4P regulate plasmodesmata opening and closing, largely independent of callose levels. This finding introduces a new perspective on how intercellular molecular flow is regulated in plants, and opens the door to a more intricate regulatory mechanism than previously thought. How could MCTP/PI4P dynamically regulate molecular flow across plasmodesmata MCS? The most straightforward hypothesis is by modulating the ER-PM gap (**Figure 7D**). Theoretical models suggest that changes in plasmodesmata structure directly affect the rate of molecular flow through these channels (Deinum et al., 2019; Ostermeyer et al., 2022). This is notably influenced by the internal geometry and size of the cytoplasmic sleeve (defined as the space between the two membranes), which determines the resistance to diffusion or convective flow. The overexpression of SAC7 PI4P phosphatase leads to increased ER-PM detachment events within plasmodesmata. The MCTP family comprises 16 members in Arabidopsis, and the deletion of three members, specifically MCTP3, MCTP4, and MCTP6, leads to a noticeable alteration in ER-PM tethering within plasmodesmata. This structural modification is associated with an increase in the global molecular flow between adjacent cells. This rise in cell-cell trafficking occurs in a context where loss-of-function mutants for *mctp3/mctp4* and *mctp3/mctp4/mctp6* exhibit a significant reduction of approximately 30% to nearly 50% in the number of plasmodesmata in the root meristem (Li et al., 2023). This suggests that the transport capacity of a single plasmodesma is considerably greater in *mctp* mutant background compared to wild-type. What are the underlying molecular mechanisms responsible for regulating the ER-PM gap? Our data indicate that MCTPs operate in coordination with the anionic lipid PI4P. We show that MCTP4 establishes a direct interaction with PI4P through its C2 domains as indicated by liposome binding assays and molecular dynamics. Furthermore, molecular dynamics simulations suggest that the levels of PI4P directly influence MCTP’s attachment to the PM. This model is corroborated by the experimental finding that when PI4P production is up- or down-regulated, cell-cell transport is either hindered or facilitated. PI4P is a signaling lipid, whose level and subcellular accumulation are highly regulated during development and environmental interactions (Noack and Jaillais, 2020). Here, we showed that the PI4P phosphatase SAC7 regulates MCTP localization and plasmodesmata connectivity in a tissue dependent manner. This serves as a proof-of-concept that graded PI4P levels in a wild-type context can impact cell-cell connectivity in different cell types. The SAC7-dependent regulation of MCTP localization that we observed at the developmental scale can be considered a relatively slow response that is established during tissue differentiation. In this scenario, PI4P binding sites inside plasmodesmata are likely limiting factors for MCTP docking and ultimately will determine the overall accumulation of MCTP at cell-cell interfaces. However, PI4P regulation may also occur on a shorter timescale, within minutes or even seconds (Marković and Jaillais, 2022). This is notably the case during certain biotic or abiotic stresses. It is also possible that some MCTP C2 domains show increased PI4P-binding upon Ca^2+^ binding (Rizo and Su, 1998; Giordano et al., 2013; Yu et al., 2016). In such a scenario, PI4P and Ca^2+^ variations may regulate plasmodesmata conductance very rapidly. However, this may not directly translate in changes in MCTP levels within plasmodesmata. It is possible that changes in PI4P concentrations might induce conditional docking of MCTPs to the PM, thereby opening or closing the cytoplasmic sleeve on demand. Such scenarios have been observed in the context of other MCS. For example, in human cells, the amount of PI4P at the Golgi determines the conditional establishment of ER/Golgi contacts and can even generate traveling waves of MCS between the two organelles (Mesmin et al., 2013; Mesmin et al., 2017). Interestingly, in this example, membrane contact establishment is regulated by a PI4P phosphatase called Sac1, which is orthologous to the Arabidopsis SAC7 enzyme (Dubois and Jaillais, 2021). In addition to conditional docking, molecular dynamics suggest that MCTP4 C2 domains may interact with the PM and PI4P in different conformations (Bian et al., 2018). Whether such conformational changes exist in vivo remains to be determined but this hypothesis represents an attractive model to regulate cytoplasmic diffusion on a short time scale.

Altogether, our study shows that ER/PM tethering not only maintains plasmodesmata structure but it also suggests that the ER-PM junction can be dynamically regulated in physiological contexts to regulate the connectivity of plant cells within their tissue.

### Limitation of the study

A limitation of our work lies in the reliance on loss-of-function *mctp* alleles. The *mctp3/mctp4/mctp6* loss-of-function mutant exhibits structural defects with looser ER-PM contacts, indicating that MCTP tethers not only act as regulatory elements but also as structural elements. One possibility is that callose may not effectively block plasmodesmata due to a significant disruption in plasmodesmata structure in the *mctp* mutant lines. Moreover, we cannot exclude that callose interaction with the PM or other wall-polymer is affected in *mctp* mutant background modifying its action. While we do not expect MCTPs to affect extracellular wall polymers or wall properties directly, we cannot rule out indirect effects. Nevertheless, our data in the *sac7* mutant suggest that increased PI4P levels stabilize MCTP within plasmodesmata and reduces cell-to-cell cytoplasmic diffusion even when callose deposition is almost completely inhibited (i.e. after BRZ treatment). While we still do not fully understand the functional and structural relationship between callose deposition and ER/PM tethering within plasmodesmata, our results strongly imply that MCTP/PI4P interaction can close plasmodesmata in the absence of callose.

In this study, we focused on the function of PI4P in the regulation of MCTP localization and plasmodesmata activity. However, PI4P is not the only signaling lipid within the PM, which also includes phosphatidic acid, phosphatidylserine and PI(4,5)P_2_ (Platre et al., 2018). Because each MCTP protein has multiple C2 domains, which all have the potential to bind to one or several anionic lipids, we need to be cautious and acknowledge that PI4P is likely not the only lipid regulator at plasmodesmata. In fact, we found both PI4P and PI(4,5)P_2_ in our purified plasmodesmata fraction. It is possible that each MCTP protein will have specific lipid preferences, thereby regulating ER/PM contact in a differential manner according to the tissue analyzed and/or to environmental stresses. In addition, the interaction between MCTP C2 domains and different anionic lipid species may also be modulated by calcium. Altogether, we believe that MCTP/lipid interactions have immense regulatory potential to fine tune plasmodesmata connectivity across a broad range of time scales and with exquisite spatial and temporal specificity.

## Supporting information

Supplemental data Perez Sancho-Smokvarska-Dubois

## Acknowledgments

We would like to thank Paulo Telles de Souza for sharing Martini3 coarse grain parameter for sitosterol. Fabrice Cordelières for live-imaging data display. Jérome Hénin for assistance with colvars protocol. Christel Poujol, Jeremie Teillon and Sebastien Marais for assistance with SP8-STED-FLIM and SP5-2P. We would also like to thank Alexandre Martinière, Yohann Boutté, and Sébastien Mongrand for reading and commenting on the article. Light and electron imaging were done at the Bordeaux Imaging Center, member of the national infrastructure France-BioImaging supported by the French National Research Agency (ANR-10-INBS-04) or at the RDP lab (ENS Lyon, France). We acknowledge the contribution of the SFR Biosciences (UMS3444/CNRS, US8/Inserm, ENS de Lyon, UCBL) E. Chatre, and J. Brocard at LBIPLATIM-MICROSCOPY for assistance with imaging, and Claire Lionnet for assistance with imaging at the RDP. The lipid analysis was done at the Bordeaux Metabolome platform for lipid analysis (https://www.biomemb.cnrs.fr/en/lipidomic-plateform/) supported by Bordeaux Metabolome Facility-MetaboHUB (grant no. ANR–11–INBS–0010 to LF). We thank Yohann Boutté for his help in anionic lipid extraction, Lothar Kalmbach for the pLOK180 plasmid gift, Yasin Dagdas for the DDRGK1-mCHERRY line and Thomas Eekhout (VIB) for the scRNA seq data mining.

## Author contributions

Conceptualization: J.P-S., M. S., Y.J., E.M.B

Methodology: J.P-S., M. S., G.D., M.G., S.S., G.D., L.F., F.I., J.T., A.T., Y.J., E.M.B

Validation: J.P-S., M. S., G.D., A.T., Y.J., E.M.B

Investigation: J. P-S, M.S., G.D., M.G., S.S., G.D. V.D., M.P.P., Z. P.L., H.M., A.P., Y.L., L. F., M.S.G., P. C-Q, T. S.M., F.I., V.W., M.D., L.B., P.R., V.B., L.L.L., C.C., S.C., M.Z., Y. L., M.Z.

Supervision: Y.J., E.M.B.

Funding acquisition: A.T., Y.J., E.M.B,

Writing – original draft: J.P-S, M.S., Y.J., E.M.B

Writing – review & editing: J.P-S, M.S., G.D., V.D., A.P., P. C-Q, M.P.P., J.T., E.R. M.B., Y.H., A.T., Y.J., E.M.B

## Data and materials availability

All data are available in the main text or the supplementary materials

## METHODS

### Plant material

*Arabidopsis thaliana*, accession Columbia (Col-0), (which we are referring to as wild-type-WT) and transgenic lines (all in Columbia background) were surface sterilized, vernalized at 4C during 2 days and grown vertically on solid half-strength Murashige and Skoog media supplemented with vitamins (2.15 g/L), MES (0.5 g/L), sucrose (10 g/L) and plant agar (7 g/L), pH 5.7 for 6 days at 22^○^C in a 16-h light/8-h dark cycle with 70% relative humidity and a light intensity of 200 µmol x m^—2^ x s^—1^ prior to use. Recently harvested, synchronized seeds were used for all phenotypical analysis. *Nicotiana benthamiana* plants were grown in an insect proof greenhouse with 18/25°C night/day.

The following lines were previously published and characterized *rhd4-1/sac7-1* (EMS) and *rhd4-3/sac7-3* (SALK_079231) (Thole et al., 2008), *EXP7pro::4xYFP* (Marquès-Bueno et al., 2016), DDRGK1-mCHERRY (Stephani et al., 2020), *SYT1pro:SYT1-GFP* (Pérez-Sancho et al., 2015b), C4H-GFP (Lee et al., 2019b), RFP-HDEL (Pain et al., 2019), *mctp3-2/mctp4-1* mutant and *MCTP4pro::*GFP-MCTP4 (Brault et al., 2019), *35Spro::DRONPA-s* (Gerlitz et al., 2018)

### Cell culture

*Arabidopsis* (*Landsberg erecta*) culture cells were cultivated as described in Bayer et al. (Bayer et al., 2004) under constant light (20 μE/m/s) at 22°C. Cells were used for plasmodesmata extraction at 7 days old.

### Cloning and plasmid construction

In this study, the constructs employed were derived from Col-0 cDNA/genomic DNA, or were synthesized. The promoter sequences were integrated into pDONR-p4RP1, while the genes and fluorescent tags were integrated into either pDONR221 or pDONR-P2RP3, utilizing the MultiSite-Gateway cloning system. For the development of an inducible genome editing vector, plasmids containing the inducible RPS5A promoter, codon-optimized and RFP-tagged Cas9, along with two sgRNAs, were assembled using the Gateway system. The assembly of all constructs was carried out through multi-Gateway reactions, wherein three segments were cloned into the destination vector, pLOK180, which express a red seed-coat selection marker (a gift from L. Kalmbach).

The expression vectors were introduced into *Agrobacterium tumefaciens* strain GVG3101, which was then used to transform Arabidopsis Col-0 or *mctp* mutants (as indicated in the figure legends) through the floral dip method, as previously described (Clough and Bent, 1998). Transformed seeds were selected based on the presence of either red seed coat or BASTA resistance, depending on the selection marker used. We employed a similar selection scheme as described in (Lebecq et al., 2022), which ensures that for each transgenic lines selected, several independent transformants, with a single insertion, show similar results. Briefly, between 20 and 24 independent T1 (Basta or Green/Red seed coat) were selected and propagated. In T2, all lines were screened using confocal microscopy for fluorescence signal and localization. Between 3 and 5 independent lines with a mono-insertion and showing a consistent, representative expression level and localization were selected and grown to the next generation. Each selected line was reanalyzed in T3 by confocal microscopy to confirm the results obtained in T2 and to select homozygous plants. At this stage, we selected one representative line for in-depth analysis of the localization.

Cloning of SAC7-related material: The SAC7 promoter (Sac7pro, 1963bp upstream of AT3G51460 START codon) was amplified from Col-0 DNA and inserted in the pENTR-5’-TOPO Gateway vector using TOPO cloning (Invitrogen™). The coding sequence (CDS) of SAC7 (ATG to STOP codon) was amplified by PCR from Col-0 cDNA and recombined by BP reaction into pDONR-P2R-P3. The mCITRINE/pDONR 221, and 2xmCHERRY-1xmyc/pDONR P221 and UBQ10pro/pDONRP4RP1 were published before (Jaillais et al., 2011; Simon et al., 2014). The LR clonase-based three-fragment recombination system (Invitrogen®) was used to produce the pSAC7::mCitrine-SAC7 into pLOK180_pR7m34g (a gift from L. Kalmbach, red seed selection) and pSAC7::2xmCherry-Myc-SAC7; pUBQ10::2xmCHERRY-Myc-SAC7into pLOK292_pFG7m34G (a gift from L. Kalmbach, green seed selection).

To generate the inducible *PDR9xve::cals3m,* a 2kb DNA fragment corresponding to *PDR9* promoter, active in the epidermis (Robe et al., 2021) was cloned into p1R4-ML:XVE (Siligato et al., 2016) and subsequently assembled by multi-site gateway technology into the destination plasmid pFG7m34GW (Kalmbach et al., 2023) together with *casl3m* (Vaténet al., 2011) in pDONR221 and the terminator tNOS in pDONRP2R-P3 (Karimi et al., 20007)

### Transient expression in *Nicotiana benthamiana*

The abaxial side of leaves of 3- to 4-week-old *Nicotiana benthamiana* plants were pressure-infiltrated with GV3101 agrobacterium strains carrying the different constructs. Prior to infiltration, the agrobacterium cultures were grown overnight in Luria and Bertani medium supplemented with rifampicin (50 mg/mL), gentamycin (25 mg/mL), and the construct-specific antibiotic at 28°C. Subsequently, the cultures were diluted to 1/10 in the same growing medium (without Rifampicin) and further cultivated until the optical density at 600 nm (OD600) reached approximately 0.6-0.8. The bacterial cells were then collected through centrifugation at 3000g for 5 min, followed by resuspension in agroinfiltration solution (10 mM MES, pH 5.6, 10 mM MgCl2, and 1 mM acetosyringone) and incubated for 2h at room temperature and in the dark. For infiltration of individual constructs, a final OD600 of 0.3 was used. For co-infiltration of two constructs, and OD600 of 0.2 each was used. Additionally, the ectopic silencing suppressor p19 was co-infiltrated in all the cases at an OD600 of 0.05. Post-infiltration, plants were kept in the same growing conditions as before for 2 more days; then leaf-discs were collected from the N. benthamiana leaves and subjected to imaging at room temperature.

### Lattice light sheet

The Lattice Light Sheet Microscope (LLSM) was built according to the technical information provided by the group of E. Betzig at Janelia Research Campus, Howard Hughes Medical Institute (HHMI), USA. The lattice light sheet was focused by a custom excitation objective 28.6× 0.66 numerical aperture 3.74-mm (Special Optics). Fluorescence was collected with a CFI Apo LWD 1.1 numerical aperture 25× 2.0-mm detection objective (Nikon) and imaged on a scientific complementary metal-oxide semiconductor (sCMOS) ORCA-Flash4.0 V2 camera (Hamamatsu). The annular mask minimum and maximum numerical apertures were 0.35 and 0.4, respectively, thus creating a light sheet with a uniform thickness over length > 30µm. All images were acquired in dithered square lattice mode. We characterized the LLSM optical resolution at l_em_=510nm using 170nm diameter beads. We found 275 +/- 5 nm and 690 +/- 10 nm, laterally and axially respectively. Images were acquired at ∼10 to 20 µm below the surface of the root. Fast sample translation with a piezo stage was used to acquire Z stacks with a step of 300 nm. Typical laser power incident on the excitation objective is ∼ 0.4mW. The raw data are 3D deconvolved with an open-source Richardson Lucy (RL) algorithm (https://github.com/dmilkie/cudaDecon) using an experimental PSF. The RL software is bundled into LLSpy, a python LLSM data processing and visualization toolbox (https://github.com/tlambert03/LLSpy). Deconvolution runs on a CUDA compatible GPU graphics card. A 3×3 median filter (noisy pixel removal) followed by background subtraction is applied before RL deconvolution with 10 iterations.

### Laser Scanning Confocal Microscopy

In the conducted live-imaging experiments, a Zeiss LSM 880 confocal microscope equipped with 40x and 63x oil-immersion objectives, as well as 40x water immersion objectives, was employed. The microscope was controlled using ZEN Black 2011 software. The experiments were carried out at a controlled temperature ranging from 22 to 24 C.

To detect GFP in the confocal mode, a filter with an excitation (Ex) wavelength of 488 nm and an emission (Em) range of 505-550 nm was utilized. For the detection of tagRFP or propidium iodide (PI) fluorescence, a filter with an Ex wavelength of 556 nm and an Em range of 570-625 nm was employed. To detect YFP fluorescence, a filter with an excitation (Ex) wavelength of 514 nm and an emission (Em) range of 520-570 nm was utilized when imaging YFP alone. When YFP was imaged together with a red reporter, we used the settings already described for GFP.

For Airyscan imaging, GFP or YFP fluorescence was detected using an excitation (EX) wavelength of 488 nm and an emission bandpass (Em BP) range of 420-480 nm in combination with 495-550 nm. On the other hand, for the detection of PI an excitation (EX) wavelength of 488 nm and an emission (Em) range of 620-700 nm in combination with a long pass (LP) filter at 645 nm were used. For callose imaging, atto550 excitation was done with 561 nm power excitation and fluorescence collected between 566 and 700 nm. Callose deposition at plasmodesmata was quantified with Fiji software and signal from the forming cell plae was excluded as indicated in Figure 3. Calcofluor aquisition was done by 405nm excitation and collected at 410-480nm. For SAC7/MCTP colocalization (Figure 6F), Airyscan imaging was conducted on 7-day-old roots using an inverted LSM 980 Axio Observer Z1/7 (Carl Zeiss group, http://www.zeiss.com/) equiped with an AiryScan 2 module, GaAsP-PMT detectors and a C Plan-Apochromat 40X objective (numerical aperture 1.3, oil immersion).

To prevent signal cross-talk, image acquisition occurred by frame when two colors were imaged simultaneously.

### Spinning Disk Microscopy

Spinning disk microscopy was used in Figure 6A-D and 7A. Observations of roots of 7-day-old seedlings were conducted using a Zeiss microscope (AxioObserver Z1, Carl Zeiss Group, http://www.zeiss.com/) equipped with a spinning disk module (CSU-W1-T3, Yokogawa, www.yokogawa.com) and a Prime 95B camera (Photometrics, https://www.photometrics.com/). We used a 40x objective (Plan-Apochromat, numerical aperture 1.1, water immersion) in Figure 7A or a 63x objective in Figure 6A-D (Plan-Apochromat, numerical aperture 1.4, oil immersion). Green Fluorescent Protein (GFP) was excited with a 488 nm laser (150mW), and the ensuing fluorescence emission was filtered via a 525/50 nm BrightLine® single-band bandpass filter (Semrock, http://www.semrock.com/). The mCITRINE marker was excited with a 515nm laser (60mW), and the resulting fluorescence emission was directed through a 578/105 nm BrightLine® single-band bandpass filter (Semrock, http://www.semrock.com/). For mCHERRY or PI labeling, an excitation laser at 561nm (80mW) was applied, and the fluorescence emission was filtered using a 609/54 nm BrightLine® single-band bandpass filter (Semrock, http://www.semrock.com/).

### Cryofixation and freeze -substitution

Five-day old seedlings of WT, *mctp3/4* and *mctp3/4/6* were grown vertically MS ½. Roots were taken and cryofixed in 20% BSA filled copper platelets (100 nm deep and 1.5 mm wide) with EM PACT1 high-pressure freezer (Leica). Subsequently, the root tissues were harvested and subjected to cryopreservation by rapidly freezing them using a high-pressure freezer (Leica EM PACT1) equipped with copper platelets (100 nm deep and 1.5 mm wide) filled with 20% bovine serum albumin (BSA) solution. The cryofixed samples were then subjected to freeze-substitution within an AFS2 system (Leica) maintained at -90 °C. The cryosubstitution process involved immersing the samples in a cryosubstitution mixture containing 0.36% uranyl acetate in pure acetone for a duration of 24 hours. Subsequently, the temperature was gradually increased at a rate of 3 °C per hour until it reached -50 °C and held steady for 3 hours. To ensure thorough substitution, the cryosubstitution mixture was replaced with pure acetone and later pure ethanol, with each solvent being applied for three 10-minute washes. The copper platelets were intentionally retained to prevent any potential loss of the samples. Infiltration of the specimens was carried out using a series of increasing concentrations of HM20 Lowicryl resin (Electron Microscopy Science), including 25% and 50% (1 hour each), 75% (2 hours), and finally 100% (overnight, 4 hours, 48 hours, with fresh resin for each bath). Following infiltration, the samples were polymerized under ultraviolet light for 24 hours at -50 °C, after which the temperature was incrementally raised at a rate of 3 °C per hour until it reached 20 °C, where it was maintained for an additional 6 hours.

### Tomography and Transmission Electron Microscopy

Fixed root embedded in HM20 resin blocks were sectioned longitudinally into slices with a thickness of 80 nm using an EM UC7 ultramicrotome (Leica) These sections were then delicately positioned onto 200 mesh copper grids. Subsequent observations were conducted employing a FEI TECNAI Spirit 120 kV electron microscope. For plasmodesmata structure we acquired tomograms in 3 to 4 roots over two technical replicates.

For the purposes of tomographic analysis, a protocol in accordance with the approach outlined by Nicolas et al., 2017a was employed. To facilitate image alignment during the tomography process, sections were coated with 5 nm colloidal fiducials. A series of tilt images spanning an angular range from -65° to 65° was acquired with a 1° angular increment using the FEI 3D explore tomography software. Subsequently, tomographic reconstructions were generated utilizing the eTomo software (http://bio3d.colorado.edu/imod/). Segmentation tasks were carried out employing the 3dMOD software (https://bio3d.colorado.edu/imod/doc/3dmodguide.html).

### Forester Resonance Energy Transfer-Fluorescence Lifetime Imaging (FRET-FLIM)

The confocal microscope used for FRET FLIM was a Leica SP8 WLL2 on an inverted stand DMI6000 (Leica Microsystems, Mannheim, Germany), using objectives HC Plan Apo CS2 40X oil NA 1.30. The confocal microscope was equipped of a pulsed white light laser 2 (WLL2) with freely tuneable excitation from 470 to 670 nm (1 nm steps) and a pulsed diode at 440 nm and also a diode laser at 405 nm.

The scanning was done using either a conventional scanner (10Hz to 1800 Hz) or a resonant scanner (8000Hz). The microscope was composed of 2 internal photomultiplier tubes (PMT), 2 internal hybrid detectors and 1 external PMT for transmission. For excitation of the donor *pMCTP4::e-YFP-MCTP4*, 516nm laser was used. The confocal microscope was equipped with the FALCON module for FLIM measurements. We used 5 iterations in order to acquire enough photon counts (10-4). Monoexponential fit was applied for both donor alone and donor with the acceptor (*pMCTP3::mCherryMCTP3*) and values were extracted only from the plasmodesmata signal.

### Two-photon microscopy and cell diffusion assay with DRONPA-s

1-day-old 35S::DRONPA-s seedlings was cultivated on ½ MS plates under long-day conditions at 21 °C with 70% humidity. Subsequently, these seedlings were transferred to either DMSO or 200nM Brassinolide plates for an additional 24 hours. The root tips of the seedlings were subjected to analysis using a Leica TCS SP5-Multi photon confocal microscope setup. Prior to imaging, the plants underwent a 2-minute incubation in a 1/500 propidium iodide (PI) solution in water. PI was excited at 488 nm and its emissions were detected at 620-700 nm. Leica LAS AF software was employed for the quantification of fluorescence intensities.

The movement of DRONPA-s was captured as follows:

1. DRONPA-s deactivation: Root tips were illuminated with 488 nm light for 45 seconds at 70% of the total laser intensity using an argon laser (20mW, Leica Microsystems).
2. DRONPA-s activation: Three Regions of Interest (ROIs), each corresponding to a single cell per root, were selected at the cell center to avoid activating adjacent cells. These ROIs were exposed to 5 seconds of 800 nm light using a two-photon IR (Titane-Saphir pulsating laser). The laser power used was approximately 2.6W with a 20% gain.
3. DRONPA-s acquisition: Detection was conducted 2 seconds after the single cell activation and continued for 120 seconds, using a 20% 488 nm argon laser at an emission range of 500-575 nm.

DRONPA-s movement analysis involved measuring mean fluorescence intensities from the activated cell and the two adjacent cells using ImageJ. DRONPA movement was calculated based on the fluorescent signal transferred from the activated cell to the adjacent cells. Any X-Y drifts were corrected with the StackReg plugin. Following normalization and background subtraction, the values were converted into a percentage of the average fluorescence from the two adjacent cells. The activated cell was considered to have 100% DRONPA-s molecules activated in the ROI. A total of more than 20 roots were analyzed, with 2-3 ROIs per root.

### Fluorescence Recovery After Photobleaching

PI permeability assessments were made using FRAP on six day-old Arabidopsis root tips co-stained with CFDA (50μg/mL) and Propidium iodide.Roots were incubated in an aqueous CFDA solution for 5 minutes, then successively washed out in 3 water baths and mounted with propidium iodide in water for imaging FRAP experiments were conducted using a Zeiss LSM 880 confocal microscope, which was equipped with a Zeiss CPL APO x 40 oil-immersion objective (numerical aperture 1.3). To excite CFDA, a 488 nm wavelength was utilized with 100% of argon laser power, and the fluorescence emission was collected within the 505–550 nm range using a GaAsp detector. Photobleaching was performed on rectangular regions of interest (ROIs) encompassing three epidermal cells, with the exciting laser wavelengths set to 100%. The FRAP procedure involved capturing 10 pre-bleach images, followed by 15 iterations of bleaching, with a pixel dwell time of 1.31 µs. Subsequently, 90 post-bleach images were acquired. The recovery profiles were subjected to background subtraction and then double normalization through the FRAP analysis website https://easyfrap.vmnet.upatras.gr/ (Koulouras et al., 2018).

### Quantification of the Apico Basal Vs Intracellular Fluorescence Ratio (APFIR)

Two methods were used to obtain the APFIR, a “manual” approach with a draw-by-hand fluorescence extraction ROI method on Fiji (Figure 6) and an “automated” pipeline with a Fiji macro (Figure 7). In figure 6, we could not use the automated approach because the signals from the various ER markers were too different (perinuclear and diffuse vs subcortical) leading to error in the segmentation.

#### Fluorescence extraction

*Manual method*: the intracellular fluorescence was extracted on each cells by drawing an ROI close to the nucleus to capture most of the perinuclear ER, the apico-basal fluorescence was extracted by drawing two ROI at the apical and basal membranes of each cells, the background fluorescence was extracted by drawing an ROI inside the nucleus. *Automated method*: This method requires a segmentation of all the cells, which is done with the red channel (propidium iodine staining of the root). Subsequently to segmentation, the fluorescence of interest (green channel) is extracted at different ROI automatically determined by the macro for each cells: the intracellular fluorescence (the entire fluorescence inside the cell); the apical membrane fluorescence, and the basal membrane fluorescence. The macro can be found at: https://github.com/RDP-vbayle/SiCE_FIJI_Macro/blob/main/misc/MacroAPFIR_Perez-Sancho_et_al_221020.ijm

#### Calculation of the APFIR

No background substraction is done on the ROI fluorescence values obtained via the automated method. For the fluorescence values obtained via the manual method, the values of the ROI fluorescence (apical, basal, intracellular) is subtracted by the value of the background of the same cell. For each cell, the APFIR is calculated as the mean apico-basal fluorescence divided by the intracellular fluorescence.

### Co-Immunoprecipitation in *Nicotiana benthamiana*

The appropriate constructs were transiently expressed in Nicotiana bentamiana (see Transient expression in *Nicotiana benthamiana* section). Two plants per construct combination and two whole leaves per plant were agroinfiltrated. Using a puncher, 10 leaf-discs were collected for each leaf, so 40 leaf-discs per construct combination, and immediately frozen in liquid nitrogen (aprox 0.5 g of tissue). Frozen tissue was grinded using mortar and pestle and total proteins were extracted with 1 mL of extraction buffer (150 mM Tris-HCl, pH 7.5; 150 mM NaCl; 10 % glycerol; 10 mM EDTA, pH 8; 1mM NaF; 1 mM Na2MoO4; 10 mM DTT; 0.5 mM PMSF; 1% (v/v) P9599 protease inhibitor cocktail (Sigma); 1 % (v/v) Igepal) by incubating during 40 min at 4 C with continuous mixing in an end-over-end rocker. Samples were centrifuged for 20 min at 4C and 9,000 g. Supernatants were filtered by gravity through Poly-PrepChromatography Columns (#731-1550 Bio-Rad). For each sample, 50 uL of the filtered supernatant were collected and mixed with 50 uL of Laemmli buffer (Tris-HCl pH 6.8 125 mM; 4% SDS; 20 % (v/v) glycerol; 2 % (v/v) beta-mercaptoethanol; 0.01 % bromophenol blue) for immunoblot analysis as input. The remaining filtered supernatants were mixed in a 1:1 proportion with wash buffer (extraction buffer without Igepal) to reach a final concentration of 0.5 % Igepal in the samples before adding the GFP-Trap agarose beads (as recommended by the manufacturer to avoid nonspecific binding). 15 uL of equilibrated GFP-Trap agarose beads were added to each sample and incubated during 2 h at 4 C with continuous mixing. After incubation, samples were centrifuged 30 s at 500 g to precipitate the beads Then beads were washed 3 times with a wash buffer. After the third wash, a final centrifugation during 30 s at 2000 g was performed and the supernatant was discarded. Finally, immunoprecipitated proteins were eluted by incubating the beads with 50 uL of Laemmli buffer at 70 C for 20 min followed by 2 min centrifugation at 2500 g and recovery of the supernatant. Input and immunoprecipitated samples were analyzed by western blot.

### Lipid inhibitor treatments

PAO treatment was used to inhibit PI4kinases (Simon et al., 2016b). 25-30 minutes of 60µM PAO was used to 5 days old *p35S::DRONPA-s* in WT and *mctp4/3* mutants background then imaged. Wortmanin was used to inhibit PI3 kinase (Gomez et al., 2022) at a concentration of 10µM for 40 minutes and YM 201653 was used to inhibit PI3,5kinase at a concentration of 2µM and treated for 1 hour.

### Environmental and hormonal triggers with brasinosteroids and NaCL

For Brasinolide (BL) and Brassinazole (BRZ) treatment 4 days old roots of *p35S::DRONPA-s* in WT and mctp4/3 background were transferred in ½ MS plate containing 200nM BL or 1µM BRZ and grown for 24h before imaging. DMSO was used as a mock solution/ Salt treatment was performed by growing the same lines for 5 days and 20mM NaCl was applied 15 minutes before imaging. Water was used as a control solution.

### Callose immunostaining

Arabidopsis seedlings were vertically cultivated on ½MS agar plates for a duration of 4 days. Subsequently, they were transferred to fresh media and subjected to various treatments for an additional 24 hours. The treatments included 200 nM BR, 1 µM BRZ, DMSO mock, and a 15-minute exposure to 20mM NaCl.

The immunolocalization procedure was performed following a previously published protocol (Pendle and Benitez-Alfonso, 2015). All solutions were prepared in microtubule stabilization buffer (MTSB), comprising 50 mM PIPES, 5 mM EGTA, 5 mM MgSO4, and adjusted to pH 7 with KOH. Seedlings were fixed in 4% (v/v) paraformaldehyde for 1h followed by 4 washes with MTSB. Root tips were then excised and mounted on microscopy slides coated with poly-lysine.

Immunolabelling was performed using the Immunorobot InSituPro VSI from Protigene. Briefly, the procedure includes cell-wall permeabilization with driselase 2% during 15 min and plasma membrane permeabilization by 10 % DMSO and 3 % Igepal during 1 h. To prevent nonspecific binding, the samples were subjected to a blocking step using 5% neutral donkey serum before incubation with the primary antibody. The callose antibody (obtained from Australia Biosupplies) was appropriately diluted to 1/500 in MTSB supplemented with 5% (v/v) neutral donkey serum and incubated with the samples at room temperature for 4 hours.

For the subsequent step, the secondary anti-mouse antibody, coupled to atto550 (Merk) or alexa fluor 594, was diluted to 1/1000 in MTSB buffer containing 5% (v/v) neutral donkey serum. This antibody solution was then applied to the samples and allowed to incubate for 1 hour.

### Root phenotyping

#### Phenotyping of the root growth rate

(Schindelin et al., 2012; Slovak et al., 2014; Platre et al., 2023). Wild-type, single (*mctp3-2, mctp3-1, mctp4-1, mctp4-2, mctp6-1*), double (*mctp3-2/mctp4-1, mctp3-1/mctp4-1, mctp3-2/mctp6-1*) and triple (*mctp3-1/mctp4-1/mctp6-1*) mutant seeds were sowed in 12-cm x 12-cm square plates filled with 1/2 MS medium containing 0.8% of agar and 0.5% of sucrose and were stratified for 2-3 days at 4°C before being transfer to a growth chamber. Six days after planting, about 20 seedlings were transferred using forceps to an imaging chamber (Lab-Tek, Chambers #1.0 Borosilicate Coverglass System, catalog number: 155361) filled with the identical medium described above. Note that the transfer took about 45-60 seconds. Images were acquired every minute for 6 hours, in brightfield conditions using a Keyence® microscope model BZ-X810 with a BZ NIKON Objective Lens 2X CFI Plan-Apo Lambda (Platre et al., 2021). The same plants were then transferred to 12-cm x 12-cm square plates filled with 1/2 MS medium containing 0.8% of agar and 0.5% of sucrose and grown to three more days and scanned to calculate the root growth rate per day.

#### Quantification of the root growth rate

For time lapse analysis of the root growth rate the script was used “ SCRIPT_MCTP” and is available on https://github.com/mplatre/SCRIPT_MCTP.git. To calculate the root growth rate per day the plates were scanned using BRAT software (Slovak et al., 2014). We then calculated the root length for three days and divided by three to evaluate the root growth rate per day using Fiji (Schindelin et al., 2012).

#### Root hair characterization and complementation analysis

For quantitative root hair analysis, seedling were grown in vitro, in a RH-media (2.15 g/L of MS (Duchefa, "0.5 MS"), 20 g/L of sucrose, 100 mg/L of myo-inositol, 0.5 mg/L of nicotinic acid, 0.5 mg/L of Vitamin B6, 0.1 mg/L of Vitamin B1, 2 mg/L of Glycine, 6 g/L Phytagel (Sigma-Aldrich)). Root hair length was assessed in 5-day-old seedlings, specifically focusing on homozygous T3 lines. Root hairs falling within the focal plane, spanning from 2000 μm to 4000 μm distance from the root apex, were subjected to measurement. A total of around 400 to 600 root hairs were measured for each transgenic line across two separate repetitions (approximately 200 to 300 root hairs per repetition, encompassing roughly 10 roots per repetition).

### Liposome binding assays

Experiment was performed as in Pérez-Sancho et al. (Pérez-sancho et al., 2016), with small modifications:

#### Protein production and purification

The C2B domain of MCTP4 was expressed as His-tagged protein using the expression vector pET28 and transformed in E. Coli BL21 (DE3). A 5 mL culture of transformed bacteria in LB medium with 50 µg/mL of Kanamycin (Kan) was grown overnight at 37 °C with continuous rotation at 200 rpm. 50 mL LB medium + Kan was inoculated with 2 mL of the overnight grown culture and incubated at 37 °C for 2-3 h, until the OD600 was between 0.6 and 0.8. Then, the expression of MCTP4-C2B was induced by adding 100μM IPTG to the culture and incubating overnight at 18 °C. A sample of the culture before and after protein induction was collected to assay protein production by SDS-PAGE. Cell culture was centrifuged at 1600 g during 15 min and the pellet (1 g) was resuspended in 3 mL of buffer A (HEPES NaOH 50 mM pH 7.6, NaCl 150 mM, Glycerol 10%, DTT 1 mM). Cells were disrupted by sonication (three cycles of 2 min, with 20 s on and 40 s off and 60 % amplitude, keeping the tube on ice) and the suspension was centrifuged at 100 000 g and 4 °C for 30 min. The supernatant was collected, mixed with 600 µL of Ni-NTA beads slurry (300 µL beads) and 125 µL of Benzoase (250 U/ µL), and incubated for 1 h at 4°C wit rotation to allow proteins to bind to the beads. The mixture was centrifuged at 5000 g and 4 °C for 5 min and the pelleted beads were washed 5 times with 1.5 mL of buffer A. Finally, proteins were eluted with buffer A containing 500 mM imidazole and their concentration was determined by Nanodrop (Mw 21278 and Ƹ 32430).

#### Liposome preparation

Liposomes of different composition were prepared by mixing the corresponding lipids stocks in chloroform. Lipid stocks were: PC at 10 mg/mL, PS at 10 mg/mL, PI(4)P at 5 mg/mL, PA at 10 mg/mL, PI(4,5)P2 at 5 mg/mL, and DOPE-Rho (1,2-dioleoyl-sn-glycero-3-phosphoethanolamine conjugated to rhodamine) at 1 mg/mL (used to visualize the lipid fraction during the preparation); and were mixed as indicated in the table below:

| | PC ($\mu\text{L}$) | PS ($\mu\text{L}$) | PI(4)P ($\mu\text{L}$) | PA ($\mu\text{L}$) | PI(4,5)P2 ($\mu\text{L}$) | DOPE-Rho ($\mu\text{L}$) |
| --- | --- | --- | --- | --- | --- | --- |
| PC | 78 | 0 | 0 | 0 | 0 | 13 |
| PC:PS | 62 | 20 | 0 | 0 | 0 | 13 |
| PC:PS:PI(4)P | 42 | 20 | 40 | 0 | 0 | 13 |
| **PC:PS:PA** | 50 | 20 | 0 | 7 | 0 | 13 |
| **PC:PS:PI(4,5)P2** | 42 | 20 | 0 | 0 | 44 | 13 |

Lipid mixtures were dried as a thin layer using a nitrogen stream and the lipid layers were resuspended in 1 mL of buffer A by vortexing at room temperature during 20 min. Then, they were subjected to 5 cycles of freezing in liquid nitrogen and thawing in a water bath at 37 °C and extruded 10 times through an 800 nm pore size polycarbonate membrane.

#### Liposome binding and sedimentation

10 µg of purified MCTP4-C2B and 40 µL of liposomes were mixed in buffer A containing 10 mM CaCl2 (1 mix for each different liposome composition) until a final volume of 200 µL and incubated at room temperature for 15 min without agitation. Large multilamellar vesicles (and the proteins bound to them) were collected by centrifugation at 300 000 g and 4°C for 45 min. 195 µL of supernatants were recovered and the pellets were resuspended in 195 µL of buffer A. Supernatant and resuspended-pellet fractions were mix with Coomassie blue buffer and run into SDS-PAGE for protein separation and quantification. Only one band corresponding to the C2B domain of MCTP4 was observed for each of the samples and each of the repetitions of the experiment. Gels were imaged using a Bio-rad’s stain-free imager on TGX-Stain-Free acrylamide gels and proteins quantified using Fiji.

### Plasmodesmata purification and lipid analysis via liquid chromatography-tandem mass spectrometry (LC-MS/MS)

The cell wall fractions were prepared from 6-day-old Arabidopsis suspension cultured cells, following the methodology outlined by Bayer et al. (Bayer et al., 2004). However, in this study, the cell disruption process was repeated five consecutive times using the N2 cell disruption vessel, in contrast to the standard three repetitions. To obtain the plasmodesmata-enriched membrane fraction, purified wall fragments were utilized, employing the method described by (Fouillen et al., 2022). Briefly, purified cell walls were subjected to digestion with 0.7% (w/v) cellulase R10 (Karlan) in a digestion buffer comprising 10 mM MES (pH 5.5) and 4.4% mannitol, supplemented with 1 μM phenylmethylsulfonyl fluoride and a complete protease inhibitor cocktail (Roche Diagnostic). The digestion process was carried out at 37°C for 1.5 hours with continuous shaking at 50 to 100 rpm. Following the digestion step, centrifugation was performed at 5850g for 5 minutes at 4°C to separate the supernatant and pellet fractions. The pellet underwent additional washing in an excess volume of digestion buffer before undergoing a second centrifugation step. The two supernatants were combined and subjected to further centrifugation at 110,000g for 40 minutes at 4°C. The resulting pellet, which contained plasmodesmata-derived membranes, was washed with an excess volume of cold Tris-buffered saline (TBS; 20 mM Tris-HCl, 0.14 M NaCl, and 2.5 mM KCl, pH 7.4). Finally, the pellet was resuspended in cold 1× TBS containing protease inhibitors (Roche Diagnostic). It is noteworthy that approximately 600 mL of cultured cells was employed to yield 100 μg of plasmodesmata-enriched membrane fraction. The protein amount in the samples was determined using a bicinchoninic acid protein assay with BSA as the standard. For the purification of plasmodesmata-enriched membrane fractions from cells subjected to sterol inhibitor treatment, fen was added to 5-day-old liquid cultured Arabidopsis cells at a final concentration of 250 μg/mL (stock solution 500 μg/μL in DMSO) and incubated as previously described for 48 hours before the purification process.

For anionic lipid analysis we followed the protocol described in (Genva et al., 2023). In brief, plasmodesmata fraction was incubated with 725 µL of a MeOH/CHCl_3_/1M HCl (2/1/0.1 v/v/v) solution and 150 µL water in presence of 10 ng of 17:0-20:4 PI(4)P standard (Avanti). 750 µl of chloroform and 170 µl HCl 2M were then added. The samples were vigorously shaken and centrifuged. The lower phase was washed with 700 µl of the upper phase of a mix of methanol:chloroform:HCl 0.01M 1:2:0.75, vortexed and centrifuged. Then, samples were kept overnight at -20 °C.

The organic phase was transferred to a new Eppendorf and the methylation reaction was carried out. For this purpose, 50 µL of TMS-diazomethane (2M in hexane) were added to each sample. After 10 min, the reaction was stopped by adding 6 µL of glacial acetic acid. 700 µL of the upper phase of a mix of MeOH/CHCl_3_/H_2_O (1/2/0.75 v/v/v) was added to each sample which was then vortexed and centrifuged (1500 x *g*, 3 min). The upper phase was removed and the washing step was repeated once. Finally, the lower organic phases were transferred to a new Eppendorf. Following the addition of 100 µL MeOH/H2O (9:1 v/v), the samples were concentrated under a gentle flow of air until only a drop remained. 80 µL of methanol were added to the samples, which were submitted to ultrasounds for 1 minute, before adding 20 µL water, and were submitted to 1 more-minute ultrasound. The samples were finally transferred to HPLC vials for analysis.

Analysis of methylated anionic phospholipids (Genva et al., 2023) were performed using a liquid chromatography system (1290 Infinity II, Agilent) coupled to a QTRAP 6500 mass spectrometer (ABSciex). The chromatographic separation of anionic phospholipid species was performed on a reverse phase C18 column (SUPELCOSIL ABZ PLUS; 10 cm x 2.1 mm, 3µm, Merck) using methanol/water (3/2) as solvent A and isopropanol/methanol (4/1) as solvent B at a flow rate of 0.2 mL/min. All solvents are supplemented with 0.1% HCCOOH and 10 mM ammonium formate. 10 µL of samples were injected and the percentage of solvent B during the gradient elution was the following: 0–20 min, 45%; 40 min, 60%; 50 min, 80%. The column temperature was kept at 40 °C. Mass spectrometry analysis was performed in the positive ionization mode. Mass spectrometry data were treated using the MultiQuant software (ABSciex). Nitrogen was used for the curtain gas (set to 35), gas 1 (set to 40), and gas 2 (set to 40). Needle voltage was at +5500 V with needle heating at 350 °C; the declustering potential was +10 V. The collision gas was also nitrogen; it was set between 26 to 45 eV according to the lipid classes.

### Identification of peptides by mass spectrometry

Each sample was deposited on a 10% acrylamide SDS-PAGE. After a short migration, gels were stained with colloidal blue overnight. The bands of interest from the SDS-PAGE gel were cut and destained. Reduction of disulfide bonds and alkylation of cysteine were performed, and proteins were finally digested overnight with trypsin. Resulting peptides were extracted from the gel, acidified and were injected on a Ultimate 3000 nanoLC system (Dionex, Amsterdam, The Netherlands) coupled to a Electrospray Orbitrap Fusion™ Lumos™ Tribrid™ Mass Spectrometer (Thermo Fisher Scientific, San Jose, CA). The spectra obtained were analyzed by Proteome Discoverer 3.1 with Sequest HT as the search algorithm, using an *Arabidopsis thaliana* protein database (Araport_20220103; 40,668 entries), including 2 missed cleavage sites by trypsin, carbamidomethylation of cysteine (+57 Da) as static modification) and oxidation of methionine (+16 Da), acetylation of protein N-term (+42 Da), Methionine loss (-131 Da) methionine loss/acetylation (+89 Da) as dynamic modification. Peptide validation was performed using Percolator algorithm and only “high confidence” peptides were retained corresponding to a 1% False Positive Rate at peptide level. Peaks were detected and integrated using the Minora algorithm embedded in Proteome Discoverer.

Protein abundances ware calculated as the sum of the abundance of corresponding unique peptides. Normalization was performed based on total *Arabidopsis thaliana* protein amount. Protein ratios were calculated as the median of all possible pairwise peptide ratios. A t-test was calculated based on background population of peptides or proteins. Quantitative data were considered for proteins quantified by a minimum of two peptides and a statistical p-value lower than 0.05.

### Molecular dynamics

#### Membrane Creation

We used the Insane program (Jumper et al., 2021) to generate three membranes with a composition representative of the plasma membrane: (70%,68%,64%) phosphatidylcholine (PC), 20% Sitosterol, 10% phosphatidylserine (PS), (0%, 2%, 6%) PI(4)P. Two lipids present in our modeled membrane were not present in the Martini 3 topology library: PS of plants that have a longer tail (18:2 24:1) and sitosterol. The topology for PS (18:2 24:1) was created by analogy, using the topology files of PIPS and DNPS. The Sitosterol model was obtained from Paulo C. T. Souza and Luís Borges Araújo following the recent parameterization of cholesterol in MARTINI 3 (Borges-Araújo et al., 2023).

#### Protein Setup

Three prediction tools, AlphaFold Monomer (Jumper et al., 2021), AlphaFold Multimer (Souza et al., 2021), and RosettaFold (Baek et al., 2021), were employed to determine the 3D structure of MCTP4. These methods converged on a single solution for the C2 domains.

All simulations were conducted using Gromacs 2021.5 (https://manual.gromacs.org/2021.5/release-notes/index.html), supplemented with the Colvar plugin (Fiorin et al., 2013). The Martini 3 force field (Souza et al., 2021) was utilized. We applied a constraint network to the protein to stabilize the 3D structure of the cytoplasmic domain. The protein’s center of mass was positioned 10 nm from the membrane. We replicated each condition five times. Each system underwent minimization and equilibration at 300K. The protocol included an initial phase of energy minimization, followed by multi-stage equilibration, and concluded with a production simulation. Energy minimization was performed in two iterations of 5,000 steps each, using the steepest descent method. The simulations were then equilibrated in five stages, using time steps of 2, 5, 10, 15, and 20 fs. A target temperature of 300 K was maintained using the v-rescale thermostat, with a coupling constant of 1 ps. A semi-isotropic pressure of 1 bar was maintained using the Parrinello-Rahman barostat, with a compressibility of 4.5 × 10^-5 bar^-1 and a relaxation time constant of 12 ps. Long-range interactions were handled with a cutoff radius of 1.1 nm for both van der Waals and Coulombic interactions, implementing a switching function from 1.0 nm for van der Waals. The production simulations were executed in an NPT ensemble with a time step of 20 fs for a total simulation time of 3 microseconds.

#### Simulation Condition

##### Colvars

Two collective variables (colvars) were used to maintain a realistic orientation of the protein relative to the membrane. The colvar tilt was chosen to control the protein’s tilt angle, and the colvar distanceZ was implemented to prevent the protein from passing through the periodic box.

##### Protein Variants

Specific mutations were introduced to study their impact on the functionality and interaction of proteins with membranes. These mutations were identified by analyzing contacts in our simulations with PI4P and using a prediction tool for membrane-interacting residues, hydrophobic protrusions (https://reuter-group.github.io/peprmint/pepr2vis). The following mutations were explored:

Five residues on the C2C domain (D217N, K218Q, R220Q, K318Q, K322Q). To observe the effect of the presence or absence of PI4P on the interaction of the protein with the membrane we performed simulations with various PI4P concentrations (without PI4P or with a 2% concentration of PI4P). Then to assess the impact of these modifications on the domain of the protein, we perform simulations C2C variant. In each case, all three domains of the protein were retained in the mutants to ensure a comprehensive analysis.

##### Calculation of Minimum Distances

To evaluate interactions between mutated proteins and PI4P molecules, the minimum distance between the mutated regions of the proteins and PI4P was calculated using the gmx mindist tool from GROMACS. In the condition without PI4P we compute the distance with the membrane.

##### Distance Analysis

The minimum distances obtained were used to generate a histogram representing the distribution of observed distances. The distance histogram was divided into several bins (distance intervals). The first bin, representing the shortest distances (>0.5nm), was specifically analyzed to identify the closest contacts between proteins and PI4P. The contact distances calculated with gmx for each domain (C2B, C2C, and C2D) were obtained as previously described in the minimum distance section. We performed an ANOVA test to statistically evaluate these contacts, ensuring all conditions such as homogeneity of variances and normality were met.

### · ● Quantification and Statistical Analysis

Statistical analyses were performed in GraphPad Prism (GraphPad Software). All experiments were independently repeated a minimum of 2–3 times (indicated in figure legends). Data are expressed as median unless specified in the figure legends.

## References

Abou-Saleh RH, Hernandez-Gomez MC, Amsbury S, Paniagua C, Bourdon M, Miyashima S, Helariutta Y, Fuller M, Budtova T, Connell SD, et al (2018) Interactions between callose and cellulose revealed through the analysis of biopolymer mixtures. Nat Commun 9: 4538

Amari K, Boutant E, Hofmann C, Schmitt-Keichinger C, Fernandez-Calvino L, Didier P, Lerich A, Mutterer J, Thomas CL, Heinlein M, et al (2010) A family of plasmodesmal proteins with receptor-like properties for plant viral movement proteins. PLoS Pathog 6: 1–10

Amsbury S, Kirk P, Benitez-Alfonso Y (2017) Emerging models on the regulation of intercellular transport by plasmodesmata-associated callose. J Exp Bot 69: 105–115

Baek M, DiMaio F, Anishchenko I, Dauparas J, Ovchinnikov S, Lee GR, Wang J, Cong Q, Kinch LN, Dustin Schaeffer R, et al (2021) Accurate prediction of protein structures and interactions using a three-track neural network. Science (80-) 373: 871–876

Balla T (2018) Ca2+ and lipid signals hold hands at endoplasmic reticulum–plasma membrane contact sites. J Physiol 596: 2709–2716

Bayer E, Thomas CLL, Maule AJJ (2004) Plasmodesmata in Arabidopsis thaliana suspension cells. Protoplasma 223: 93–102

Benavente JL, Siliqi D, Infantes L, Lagartera L, Mills A, Gago F, Ruiz-López N, Botella MA, Sánchez-Barrena MJ, Albert A (2021) The structure and flexibility analysis of the Arabidopsis synaptotagmin 1 reveal the basis of its regulation at membrane contact sites. Life Sci Alliance. doi: 10.26508/LSA.202101152

Benitez-Alfonso Y, Faulkner C, Pendle A, Miyashima S, Helariutta Y, Maule A (2013) Symplastic intercellular connectivity regulates Llteral root patterning. Dev Cell 26: 136–147

Bharathan NK, Giang W, Hoffman CL, Aaron JS, Khuon S, Chew TL, Preibisch S, Trautman ET, Heinrich L, Bogovic J, et al (2023) Architecture and dynamics of a desmosome–endoplasmic reticulum complex. Nat Cell Biol 25: 823–835

Bian X, Saheki Y, De Camilli P (2018) Ca 2+ releases E-Syt1 autoinhibition to couple ER -plasma membrane tethering with lipid transport. EMBO J 37: 219–234

Borges-Araújo L, Borges-Araújo AC, Ozturk TN, Ramirez-Echemendia DP, Fábián B, Carpenter TS, Thallmair S, Barnoud J, Ingólfsson HI, Hummer G, et al (2023) Martini 3 Coarse-Grained Force Field for Cholesterol. J Chem Theory Comput. doi: 10.1021/acs.jctc.3c00547

Brault ML, Petit JD, Immel F, Nicolas WJ, Glavier M, Brocard L, Gaston A, Fouché M, Hawkins TJ, Crowet J-M, et al (2019) Multiple C2 domains and transmembrane region proteins (MCTPs) tether membranes at plasmodesmata. EMBO Rep e47182

Cheval C, Faulkner C (2018) Plasmodesmal regulation during plant–pathogen interactions. New Phytol 217: 62–67

Chung J yun, Torta F, Masai K, Lucast L, Czapla H, Tanner LB, Narayanaswamy P, Wenk MR, Nakatsu F, De Camilli P, et al (2015) PI4P/phosphatidylserine countertransport at ORP5- and ORP8-mediated ER – plasma membrane contacts. Science (80-) 349: 428–432

Clough SJ, Bent AF (1998) Floral dip : a simplified method for Agrobacterium-mediated transformation of Arabidopsis thaliana. Plant J 16: 735–743

Collado J, Kalemanov M, Martinez-Sanchez A, Campelo F, Baumeister W, Stefan CJ, Fernandez-Busnadiego R (2019) Tricalbin-Mediated Contact Sites Control ER Curvature to Maintain Plasma Membrane Integrity. SSRN Electron J 1–61

Crawford KM, Zambryski PC (2001) Non-targeted and targeted protein movement through plasmodesmata in leaves in different developmental and physiological states. Plant Physiol 125: 1802–1812

Cui W, Lee J-YY (2016) Arabidopsis callose synthases CalS1/8 regulate plasmodesmal permeability during stress. Nat Plants 2: 16034

Daum G, Medzihradszky A, Suzaki T, Lohmann JU (2014) A mechanistic framework for non-cell autonomous stem cell induction in Arabidopsis. Proc Natl Acad Sci U S A 111: 14619–24

Deinum EE, Mulder BM, Benitez-Alfonso Y (2019) From plasmodesma geometry to effective symplasmic permeability through biophysical modelling. Elife 8: 1–40

Ding B, Turgeon R, Parthasarathy M V. (1992) Substructure of freeze-substituted plasmodesmata. Protoplasma 169: 28–41

Dong R, Saheki Y, Swarup S, Lucast L, Harper JW, De Camilli P (2016) Endosome-ER contacts control actin nucleation and retromer function through VAP-dependent regulation of PI4P. Cell 166: 408–423

Dubois GA, Jaillais Y (2021) Anionic phospholipid gradients: An uncharacterized frontier of the plant endomembrane network. Plant Physiol 185: 577–592

Eden ER, Sanchez-Heras E, Tsapara A, Sobota A, Levine TP, Futter CE (2016) Annexin A1 tethers membrane contact sites that mediate ER to endosome cholesterol transport. Dev Cell 37: 473–483

Eden ER, White IJ, Tsapara A, Futter CE (2010) Membrane contacts between endosomes and ER provide sites for PTP1B-epidermal growth factor receptor interaction. Nat Cell Biol 12: 267–72

Faulkner C, Petutschnig E, Benitez-Alfonso Y, Beck M, Robatzek S, Lipka V, Maule AJ (2013) LYM2-dependent chitin perception limits molecular flux via plasmodesmata. Proc Natl Acad Sci U S A 110: 9166–70

Fernandez-Calvino L, Faulkner C, Walshaw J, Saalbach G, Bayer E, Benitez-Alfonso Y, Maule A (2011) Arabidopsis plasmodesmal proteome. PLoS One 6: e18880

von Filseck JM, Vanni S, Mesmin B, Antonny B, Drin G (2015) A phosphatidylinositol-4-phosphate powered exchange mechanism to create a lipid gradient between membranes. Nat Commun 6: 1–12

Fiorin G, Klein ML, Hénin J (2013) Using collective variables to drive molecular dynamics simulations. Mol Phys 111: 3345–3362

Fouillen L, Claverol S, Bayer E, Grison SM (2022) Isolation of plasmodesmata membranes for lipidomic and proteomic analysis. Method Mol Biol 2457: 189–207

Friedman JR, Lackner LL, West M, DiBenedetto JR, Nunnari J, Voeltz GK (2011) ER tubules mark sites of mitochondrial division. Science (80-) 334: 358–362

Gao C, Liu X, Storme N De, Martens HJ, Schulz A, Liesche J, Gao C, Liu X, Storme N De, Jensen KH, et al (2020) Directionality of Plasmodesmata-Mediated Transport in Arabidopsis Leaves Supports Auxin Report Directionality of Plasmodesmata-Mediated Transport in Arabidopsis Leaves Supports Auxin Channeling. Curr Biol 30: 1970–1977.e4

Gaudioso-Pedraza R, Beck M, Frances L, Kirk P, Ripodas C, Niebel A, Oldroyd GEDD, Benitez-alfonso Y, de Carvalho-Niebel F, Carvalho-niebel F De, et al (2018) Callose-regulated symplastic communication coordinates symbiotic root nodule development. Curr Biol 28: 3562–3577.e6

Genva M, Fougère L, Bahammou D, Mongrand S, Boutté Y, Fouillen L (2023) A global LC–MS2-based methodology to identify and quantify anionic phospholipids in plant samples. Plant J. doi: 10.1111/tpj.16525

Gerlitz N, Gerum R, Sauer N, Stadler R (2018) Photoinducible DRONPA-s : a new tool for investigating cell – cell connectivity. Plant Physiol 94: 751–766

Giordano F, Saheki Y, Idevall-Hagren O, Colombo SF, Pirruccello M, Milosevic I, Gracheva EO, Bagriantsev SN, Borgese N, De Camilli P (2013) PI(4,5)P2-dependent and Ca2+-regulated ER-PM interactions mediated by the extended synaptotagmins. Cell 153: 1494

Gisel A, Barella S, Hempel FD, Zambryski PC (1999) Temporal and spatial regulation of symplastic trafficking during development in Arabidopsis thaliana apices. Development 126: 1879–1889

Gombos S, Miras M, Howe V, Xi L, Pottier M, Kazemein Jasemi NS, Schladt M, Ejike JO, Neumann U, Hänsch S, et al (2023) A high-confidence Physcomitrium patens plasmodesmata proteome by iterative scoring and validation reveals diversification of cell wall proteins during evolution. New Phytol 238: 637–653

Grison MS, Kirk P, Brault ML, Wu XN, Schulze WX, Benitez-Alfonso Y, Immel F, Bayer EM (2019) Plasma membrane-associated receptor-like kinases relocalize to plasmodesmata in response to osmotic stress. Plant Physiol. doi: 10.1104/pp.19.00473

Guenoune-Gelbart D, Elbaum M, Sagi G, Levy A, Epel BL (2008) Tobacco mosaic virus (TMV) replicase and movement protein function synergistically in facilitating TMV spread by lateral diffusion in the plasmodesmal desmotubule of Nicotiana benthamiana. Mol Plant Microbe Interact 21: 335–345

Guillén-Samander A, De Camilli P (2023) Endoplasmic reticulum membrane contact sites, lipid transport, and neurodegeneration. Cold Spring Harb Perspect Biol. doi: 10.1101/cshperspect.a041257

Guseman JM, Lee JS, Bogenschutz NL, Peterson KM, Virata RE, Xie B, Kanaoka MM, Hong Z, Torii KU (2010) Dysregulation of cell-to-cell connectivity and stomatal patterning by loss-of-function mutation in Arabidopsis CHORUS (GLUCAN SYNTHASE-LIKE 8). Development 137: 1731–1741

Hamasaki M, Furuta N, Matsuda A, Nezu A, Yamamoto A, Fujita N, Oomori H, Noda T, Haraguchi T, Hiraoka Y, et al (2013) Autophagosomes form at ER-mitochondria contact sites. Nature 495: 389–393

Han X, Hyun T, Zhang M, Kumar R, Koh EJ, Kang BH, Lucas W, Kim JY (2014) Auxin-callose-mediated plasmodesmal gating is essential for tropic auxin gradient formation and signaling. Dev Cell 28: 132–146

Hernández-Alvarez MI, Sebastián D, Vives S, Ivanova S, Bartoccioni P, Kakimoto P, Plana N, Veiga SR, Hernández V, Vasconcelos N, et al (2019) Deficient endoplasmic reticulum-mitochondrial phosphatidylserine transfer causes liver disease. Cell 177: 881–895.e17

Hirabayashi Y, Kwon SK, Paek H, Pernice WM, Paul MA, Lee J, Erfani P, Raczkowski A, Petrey DS, Pon LA, et al (2017) ER-mitochondria tethering by PDZD8 regulates Ca2+ dynamics in mammalian neurons. Science (80-) 358: 623–630

Hoffmann PC, Bharat TAM, Wozny MR, Boulanger J, Miller EA, Kukulski W (2019) Tricalbins Contribute to Cellular Lipid Flux and Form Curved ER-PM Contacts that Are Bridged by Rod-Shaped Structures. Dev Cell 51: 488–502.e8

Hunter K, Kimura S, Rokka A, Tran HC, Toyota M, Kukkonen JP, Wrzaczek M (2019) CRK2 enhances salt tolerance by regulating callose deposition in connection with PLDα1. Plant Physiol. doi: 10.1104/pp.19.00560

Jaillais Y, Belkhadir Y, Balsemão-Pires E, Dangl JL, Chory J (2011) Extracellular leucine-rich repeats as a platform for receptor/coreceptor complex formation. Proc Natl Acad Sci U S A 1: 1–5

Johnson B, Iuliano M, Lam T, Biederer T, De Camilli P (2023) A complex of the lipid transport ER proteins TMEM24 and C2CD2 with band 4.1 at cell-cell contacts. An ER-plasma membrane complex at cell contacts. bioRxiv. doi: 10.1101/2023.12.06.570396

Johnston MG, Breakspear A, Samwald S, Zhang D, Papp D, Faulkner C, De Keijzer J (2023) Comparative phyloproteomics identifies conserved plasmodesmal proteins. J Exp Bot 74: 1821– 1835

Jumper J, Evans R, Pritzel A, Green T, Figurnov M, Ronneberger O, Tunyasuvunakool K, Bates R, Žídek A, Potapenko A, et al (2021) Highly accurate protein structure prediction with AlphaFold. Nature 596: 583

Kalmbach, L., Bourdon, M., Belevich, I., Safran, J., Lemaire, A., Heo, J.-o., Otero, S., Blob, B., Pelloux, J., Jokitalo, E., and Helariutta, Y. (2023). Putative pectate lyase PLL12 and callose deposition through polar CALS7 are necessary for long-distance phloem transport in Arabidopsis. Current Biology 33, 926–939.e929.

Karimi, M., Bleys, A., Vanderhaeghen, R., and Hilson, P. (2007). Building blocks for plant gene assembly. Plant Physiol 145, 1183–1191. 10.1104/pp.107.110411.

Kirk P, Amsbury S, German L, Gaudioso-Pedraza R, Benitez-Alfonso Y (2022) A comparative meta-proteomic pipeline for the identification of plasmodesmata proteins and regulatory conditions in diverse plant species. BMC Biol 20: 1–21

Kitagawa M, Wu P, Balkunde R, Cunniff P, Jackson D (2022) An RNA exosome subunit mediates cell-to-cell trafficking of a homeobox mRNA via plasmodesmata. Science (80-) 375: 177–182

Koulouras G, Panagopoulos A, Rapsomaniki MA, Giakoumakis NN, Taraviras S, Lygerou Z (2018) EasyFRAP-web: A web-based tool for the analysis of fluorescence recovery after photobleaching data. Nucleic Acids Res 46: W467–W472

Kraner ME, Müller C, Sonnewald U (2017) Comparative proteomic profiling of the choline transporter-like1 (CHER1) mutant provides insights into plasmodesmata composition of fully developed Arabidopsis thaliana leaves. Plant J 92: 696–709

Kumagai K, Hanada K (2019) Structure, functions and regulation of CERT, a lipid-transfer protein for the delivery of ceramide at the ER–Golgi membrane contact sites. FEBS Lett 593: 2366–2377

Lebecq A, Doumane M, Fangain A, Bayle V, Leong JX, Rozier F, Del Marques-Bueno M, Armengot L, Boisseau R, Simon ML, et al (2022) The Arabidopsis SAC9 enzyme is enriched in a cortical population of early endosomes and restricts PI(4,5)P2 at the plasma membrane. Elife. doi: 10.7554/eLife.73837

Lee E, Vanneste S, Pérez-Sancho J, Benitez-Fuente F, Strelau M, Macho AP, Botella MA, Friml J, Rosado A (2019a) Ionic stress enhances ER–PM connectivity via phosphoinositide-associated SYT1 contact site expansion in Arabidopsis. Proc Natl Acad Sci U S A 116: 1420–1429

Lee E, Vanneste S, Pérez-Sancho J, Benitez-Fuente F, Strelau M, Macho AP, Botella MA, Friml J, Rosado A (2019b) Ionic stress enhances ER–PM connectivity via phosphoinositide-associated SYT1 contact site expansion in Arabidopsis. Proc Natl Acad Sci U S A 116: 1420–1429

Lee EK, Santana BVN, Samuels E, Benitez-Fuente F, Corsi E, Botella MA, Perez-Sancho J, Vanneste S, Friml J, Macho A, et al (2020) Rare earth elements induce cytoskeleton-dependent and PI4P-associated rearrangement of SYT1/SYT5 endoplasmic reticulum-plasma membrane contact site complexes in Arabidopsis. J Exp Bot 71: 3986–3998

Lees JA, Messa M, Sun EW, Wheeler H, Torta F, Wenk MR, De Camilli P, Reinisch KM (2017) Lipid transport by TMEM24 at ER-plasma membrane contacts regulates pulsatile insulin secretion. Science (80-). doi: 10.1126/science.aah6171

Lexy RO, Kasai K, Clark N, Fujiwara T, Sozzani R, Gallagher KL (2018) Exposure to heavy metal stress triggers changes in plasmodesmatal permeability via deposition and breakdown of callose. 69: 3715–3728

Li ZP, Moreau H, Petit JD, Souza-Moraes T, Smokvarska M, Perez-Sancho J, Petrel M, Decoeur F, Brocard L, Chambaud C, et al (2023) Plant plasmodesmata bridges form through ER-driven incomplete cytokinesis. bioRxiv

Li ZP, Paterlini A, Glavier M, Bayer EM (2021) Intercellular trafficking via plasmodesmata: molecular layers of complexity. Cell Mol Life Sci 78: 799–816

Marković V, Jaillais Y (2022) Phosphatidylinositol 4-phosphate: a key determinant of plasma membrane identity and function in plants. New Phytol 235: 867–874

Marquès-Bueno MM, Morao AK, Cayrel A, Platre MP, Barberon M, Caillieux E, Colot V, Jaillais Y, Roudier F, Vert G (2016) A versatile Multisite Gateway-compatible promoter and transgenic line collection for cell type-specific functional genomics in Arabidopsis. Plant J 85: 320–333

Mehra P, Pandey BK, Melebari D, Banda J, Leftley N, Couvreur V, Rowe J, Anfang M, De Gernier H, Morris E, et al (2022) Hydraulic flux–responsive hormone redistribution determines root branching. Science (80-) 378: 762–768

Mellor NL, Voß U, Janes G, Bennett MJ, Wells DM, Band LR (2020) Auxin fluxes through plasmodesmata modify root-tip auxin distribution. Development. doi: 10.1242/dev.181669

Mesmin B, Bigay J, Von Filseck JM, Lacas-Gervais S, Drin G, Antonny B (2013) A four-step cycle driven by PI(4)P hydrolysis directs sterol/PI(4)P exchange by the ER-Golgi Tether OSBP. Cell 155: 830–843

Mesmin B, Bigay J, Polidori J, Jamecna D, Lacas-Gervais S, Antonny B (2017) Sterol transfer, PI4P consumption, and control of membrane lipid order by endogenous OSBP. EMBO J 36: e201796687

Miras M, Pottier M, Schladt TM, Ejike JO, Redzich L, Frommer WB, Kim JY (2022a) Plasmodesmata and their role in assimilate translocation. J Plant Physiol 270: 153633

Miras M, Pottier M, Schladt TM, Ejike JO, Redzich L, Frommer WB, Kim JY (2022b) Plasmodesmata and their role in assimilate translocation. J Plant Physiol 270: 153633

Miyashima S, Roszak P, Sevilem I, Toyokura K, Blob B, Heo J, Mellor N, Help-rinta-rahko H, Otero S, Smet W, et al (2019) Mobile PEAR transcription factors integrate positional cues to prime cambial growth. Nature 565: 490–494

Nakajima K, Sena G, Nawy T, Benfey PN (2001) Intercellular movement of the putative transcription factor SHR in root patterning. Nature 413: 307–311

Naón D, Hernández-Alvarez MI, Shinjo S, Wieczor M, Ivanova S, de Brito OM, Quintana A, Hidalgo J, Palacín M, Aparicio P, et al (2023) Splice variants of mitofusin 2 shape the endoplasmic reticulum and tether it to mitochondria. Science (80-). doi: 10.1126/science.adh9351

Nicolas W, Grison M, Trépout S, Gaston A, Fouché M, Cordelières F, Oparka K, Tilsner J, Brocard L, Bayer E (2017a) Architecture and permeability of post-cytokinesis plasmodesmata lacking cytoplasmic sleeve. Nat Plants 3: 17082

Nicolas W, Grison MS, Bayer EMF (2017b) Shaping intercellular channels of plasmodesmata: the structure-to-function missing link. J Exp Bot 69: 91–103

Noack LC, Jaillais Y (2020) Functions of anionic lipids in plants. Annu Rev Plant Biol 71: 71–102

Ostermeyer GP, Jensen KH, Franzen AR, Peters WS, Knoblauch M (2022) Diversity of funnel plasmodesmata in angiosperms: the impact of geometry on plasmodesmal resistance. Plant J 110: 707–719

Otero S, Helariutta Y, Benitez-Alfonso Y (2016) Symplastic communication in organ formation and tissue patterning. Curr Opin Plant Biol 29: 21–28

Pain C, Kriechbaumer V, Kittelmann M, Hawes C, Fricker M (2019) Quantitative analysis of plant ER architecture and dynamics. Nat Commun 10: 1–15

Pérez-sancho J, Schapire AL, Botella MA, Rosado A (2016) Analysis of protein-Lipid interactions using purified C2 domains. Methods Mol Biol 1363: 175–187

Pérez-Sancho J, Vanneste S, Lee E, McFarlane H, Esteban del Valle A, Valpuesta V, Friml J, Botella MA, Rosado A (2015a) The Arabidopsis SYT1 is enriched in ER-PM contact sites and confers cellular resistance to mechanical stresses. Plant Physiol 168: 132–143

Pérez-Sancho J, Vanneste S, Lee E, McFarlane HE, Esteban Del Valle A, Valpuesta V, Friml J, Botella MA, Rosado A (2015b) The Arabidopsis synaptotagmin1 is enriched in endoplasmic reticulum-plasma membrane contact sites and confers cellular resistance to mechanical stresses. Plant Physiol 168: 132–43

Petit JD, Glavier M, Brocard L, Bayer EM Plasmodesmata ultrastructure determination using electron tomography. Method Mol. Biol.

Platre MP, Mehta P, Halvorson Z, Zhang L, Brent L, Gleason F. M, Faizi K, Goulding C, Busch W (2023) Root Walker: an automated pipeline for large scale quantification of early root growth responses at high spatial and temporal resolution. Plant J. doi: 10.1111/tpj.16493

Platre MP, Noack LC, Doumane M, Bayle V, Simon MLA, Maneta-Peyret L, Fouillen L, Stanislas T, Armengot L, Pejchar P, et al (2018) A Combinatorial lipid code shapes the electrostatic landscape of plant endomembranes. Dev Cell 45: 465–480.e11

Platre MP, Satbhai SB, Brent L, Gleason MF, Cao M, Grison M, Glavier M, Zhang L, Gaillochet C, Goeschl C, et al (2021) The receptor kinase SRF3 coordinates iron-level and flagellin dependent defense and growth responses in plants. Nat Com 13: 4445

Prinz WA, Toulmay A, Balla T (2020) The functional universe of membrane contact sites. Nat Rev Mol Cell Biol 21: 7–24

Qian T, Li C, Liu F, Xu K, Wan C, Liu Y, Yu H (2022) Arabidopsis synaptotagmin 1 mediates lipid transport in a lipid composition-dependent manner. Traffic 23: 346–356

Radulovic M, Wenzel EM, Gilani S, Holland LK, Lystad AH, Phuyal S, Olkkonen VM, Brech A, Jäättelä M, Maeda K, et al (2022) Cholesterol transfer via endoplasmic reticulum contacts mediates lysosome damage repair. EMBO J. doi: 10.15252/embj.2022112677

Rizo J, Su TC (1998) C2-domains, Structure and Function of a universal Ca2+ binding Domain. J Biol Chem 273: 15879–15882

Robe, K., Conejero, G., Gao, F., Lefebvre-Legendre, L., Sylvestre-Gonon, E., Rofidal, V., Hem, S., Rouhier, N., Barberon, M., Hecker, A., et al. (2021). Coumarin accumulation and trafficking in Arabidopsis thaliana: a complex and dynamic process. The New phytologist 229, 2062–2079. 10.1111/nph.17090.

Rowland AA, Chitwood PJ, Phillips MJ, Voeltz GK (2014) ER contact sites define the position and timing of endosome fission. Cell 159: 1027–1041

Ruiz-Lopez N, Pérez-Sancho J, del Valle AE, Haslam RP, Vanneste S, Catalá R, Perea-Resa C, van Damme D, García-Hernández S, Albert A, et al (2021) Synaptotagmins at the endoplasmic reticulum–plasma membrane contact sites maintain diacylglycerol homeostasis during abiotic stress. Plant Cell 33: 2431–2453

Sager R, Wang X, Hill K, Yoo BC, Caplan J, Nedo A, Tran T, Bennett MJ, Lee JY (2020) Auxin-dependent control of a plasmodesmal regulator creates a negative feedback loop modulating lateral root emergence. Nat Commun 11: 1–10

Saheki Y, Bian X, Schauder CM, Sawaki Y, Surma MA, Klose C, Pincet F, Reinisch KM, De Camilli P (2016) Control of plasma membrane lipid homeostasis by the extended synaptotagmins. Nat Cell Biol 18: 504–15

Salvador-Gallego R, Hoyer MJ, Voeltz GK (2017) SnapShot: Functions of endoplasmic reticulum membrane contact sites. Cell 171: 1224.e1–1224.e1

Schauder CM, Wu X, Saheki Y, Narayanaswamy P, Torta F, Wenk MR, De Camilli P, Reinisch KM (2014) Structure of a lipid-bound extended synaptotagmin indicates a role in lipid transfer. Nature 510: 552–555

Schindelin J, Arganda-Carreras I, Frise E, Kaynig V, Longair M, Pietzsch T, Preibisch S, Rueden C, Saalfeld S, Schmid B, et al (2012) Fiji: An open-source platform for biological-image analysis. Nat Methods 9: 676–682

Scorrano L, De Matteis MA, Emr S, Giordano F, Hajnóczky G, Kornmann B, Lackner LL, Levine TP, Pellegrini L, Reinisch K, et al (2019) Coming together to define membrane contact sites. Nat Commun 10: 1287

Siligato, R., Wang, X., Yadav, S.R., Lehesranta, S., Ma, G., Ursache, R., Sevilem, I., Zhang, J., Gorte, M., Prasad, K., et al. (2016). MultiSite Gateway-Compatible Cell Type-Specific Gene-Inducible System for Plants. Plant Physiol 170, 627–641.

Simon MLA, Platre MP, Assil S, Van Wijk R, Chen WY, Chory J, Dreux M, Munnik T, Jaillais Y (2014) A multi-colour/multi-affinity marker set to visualize phosphoinositide dynamics in Arabidopsis. Plant J 77: 322–337

Simon MLA, Platre MP, Marquès-bueno, Maria M, Armengot L, Stanislas T, Bayle V, Caillaud M, Jaillais Y (2016) APtdIns(4)P-driven electrostatic field controls cell membrane identity and signalling in plants. Nat Plants 20;2: 16089

Slovak R, Göschl C, Su X, Shimotani K, Shiina T, Busch W (2014) A scalable open-source pipeline for large-scale root phenotyping of Arabidopsis. Plant Cell 26: 2390–2403

Soboloff J, Spassova M a., Dziadek M a., Gill DL (2006) Calcium signals mediated by STIM and Orai proteins-A new paradigm in inter-organelle communication. Biochim Biophys Acta - Mol Cell Res 1763: 1161–1168

Song JH, Kwak S, Nam KH, Schiefelbein J, Lee MM (2019) QUIRKY regukates root epidermal cell patterning through stabilizing SCRAMBLED to control CAPRICE movement in Arabidopsis. Nat Commun 10: 1–12

Song L, Wang Y, Guo Z, Lam SM, Shui G, Cheng Y (2021) NCP2/RHD4/SAC7, SAC6 and SAC8 phosphoinositide phosphatases are required for PtdIns4P and PtdIns(4,5)P2 homeostasis and Arabidopsis development. New Phytol 231: 713–725

Souza PCT, Alessandri R, Barnoud J, Thallmair S, Faustino I, Grünewald F, Patmanidis I, Abdizadeh H, Bruininks BMH, Wassenaar TA, et al (2021) Martini 3: a general purpose force field for coarse-grained molecular dynamics. Nat Methods 2021 184 18: 382–388

Sritharan S, Versini R, Petit J, Bayer E, Taly A (2023) Deep Learning-Based Prediction of A. thaliana’s MCTP4 Structure and Exploration of Transmembrane Dynamics using Coarse-Grained Molecular Dynamics Simulations. bioRxiv 2023.08.04.552001

Stahl Y, Faulkner C (2016) Receptor complex mediated regulation of symplastic traffic. Trends Plant Sci 21: 450–459

Stahl Y, Grabowski S, Bleckmann A, Kühnemuth R, Weidtkamp-Peters S, Pinto KG, Kirschner GK, Schmid JB, Wink RH, Hülsewede A, et al (2013) Moderation of arabidopsis root stemness by CLAVATA1 and ARABIDOPSIS CRINKLY4 receptor kinase complexes. Curr Biol 23: 362– 371

Stephani M, Picchianti L, Gajic A, Beveridge R, Skarwan E, Hernandez VS de M, Mohseni A, Clavel M, Zeng Y, Naumann C, et al (2020) A cross-kingdom conserved er-phagy receptor maintains endoplasmic reticulum homeostasis during stress. Elife 9: 1–105

Tee EE, Johnston MG, Papp D, Faulkner C (2022) A PDLP-NHL3 complex integrates plasmodesmal immune signaling cascades. PNAS 120: e2216397120

Thole JM, Vermeer JEM, Zhang Y, Gadella TWJ, Nielsen E (2008) Root Hair Defective4 encodes a phosphatidylinositol-4-phosphate phosphatase required for proper root hair development in Arabidopsis thaliana. Plant Cell 20: 381–395

Tilsner J, Linnik O, Louveaux M, Roberts IM, Chapman SN, Oparka KJ (2013) Replication and trafficking of a plant virus are coupled at the entrances of plasmodesmata. J Cell Biol 201: 981–995

Tilsner J, Nicolas W, Rosado A, Bayer EM (2016) Staying Tight: Plasmodesmal Membrane Contact Sites and the Control of Cell-to-Cell Connectivity in Plants. Annu Rev Plant Biol 67: 1–28

Tilsner J, Taliansky M, Torrance L (2014) Plant Virus Movement. Encycl life Sci 1–12

Tran TM, McCubbin TJ, Bihmidine S, Julius BT, Baker RF, Schauflinger M, Weil C, Springer N, Chomet P, Wagner R, et al (2019) Maize Carbohydrate Partitioning Defective33 Encodes an MCTP Protein and Functions in Sucrose Export from Leaves. Mol Plant 12: 1278–1293

Tubiana T, Sillitoe I, Orengo C, Reuter N (2022) Dissecting peripheral protein-membrane interfaces. PLoS Comput Biol. doi: 10.1371/journal.pcbi.1010346

Tylewicz S, Bhalerao RP, Petterle A, Marttila S, Miskolczi P, Azeez A, Singh RK, Immanen J, Mähler N, Hvidsten TR, et al (2018) Photoperiodic control of seasonal growth is mediated by ABA acting on cell-cell communication. Science (80-) 360: 212–215

Vaddepalli P, Herrmann A, Fulton L, Oelschner M, Hillmer S, Stratil TF, Fastner A, Hammes UZ, Ott T, Robinson DG, et al (2014) The C2-domain protein QUIRKY and the receptor-like kinase STRUBBELIG localize to plasmodesmata and mediate tissue morphogenesis in Arabidopsis thaliana. Development 141: 4139–4148

Vatén A, Dettmer J, Wu S, Stierhof YD, Miyashima S, Yadav SR, Roberts CJ, Campilho A, Bulone V, Lichtenberger R, et al (2011) Callose biosynthesis regulates symplastic trafficking during root development. Dev Cell 21: 1144–1155

Wang P, Pleskot R, Zang J, Winkler J, Wang J, Yperman K, Zhang T, Wang K, Gong J, Guan Y, et al (2019) Plant AtEH/Pan1 proteins drive autophagosome formation at ER-PM contact sites with actin and endocytic machinery. Nat Commun. doi: 10.1038/s41467-019-12782-6

Wang X, Sager R, Cui W, Zhang C, Lu H, Lee J-Y (2013) Salicylic acid regulates Plasmodesmata closure during innate immune responses in Arabidopsis. Plant Cell 25: 2315–29

Wang Y, Perez-Sancho J, Platre MP, Callebaut B, Smokvarska M, Ferrer K, Luo Y, Nolan TM, Sato T, Busch W, et al (2023) Plasmodesmata mediate cell-to-cell transport of brassinosteroid hormones. Nat Chem Biol June: 1–11

Wilhelm LP, Wendling C, Védie B, Kobayashi T, Chenard M, Tomasetto C, Drin G, Alpy F (2017) STARD 3 mediates endoplasmic reticulum-to-endosome cholesterol transport at membrane contact sites. EMBO J 36: 1412–1433

Wong LH, Levine TP, Wirtz KW, Zilversmit DB, Pagano RE, Vance JE, Bernhard W, Rouiller C, Prinz WA, Vihtelic TS, et al (2016) Lipid transfer proteins do their thing anchored at membrane contact sites… but what is their thing? Biochem Soc Trans 44: 517–27

Wozny MR, Di Luca A, Morado DR, Picco A, Khaddaj R, Campomanes P, Ivanović L, Hoffmann PC, Miller EA, Vanni S, et al (2023) In situ architecture of the ER–mitochondria encounter structure. Nature 618: 188–192

Wu H, Carvalho P, Voeltz GK (2018a) Here, there and everywhere: The importance of ER membrane contact sites. Science (80-) 361: 466

Wu S-W, Kumar R, Bagus A, Iswanto B, Kim J-Y (2018b) Callose balancing at plasmodesmata. J Exp Bot 69: 5325–5339

Yan D, Yadav SR, Paterlini A, Nicolas WJ, Petit JD, Brocard L, Belevich I, Grison MS, Vaten A, Karami L, et al (2019) Sphingolipid biosynthesis modulates plasmodesmal ultrastructure and phloem unloading. Nat Plants 5: 604–615

Yu H, Liu Y, Gulbranson DR, Paine A, Rathore SS, Shen J (2016) Extended synaptotagmins are Ca2+-dependent lipid transfer proteins at membrane contact sites. Proc Natl Acad Sci U S A 113: 4362–4367

Zavaliev R, Ueki S, Epel BL, Citovsky V (2011) Biology of callose (β-1,3-glucan) turnover at plasmodesmata. Protoplasma 248: 117–130

Zhu T, Lucas WJ, Rost TL (1998) Directional cell-to-cell communication in the Arabidopsis root apical meristem I. An ultrastructural and functional analysis. Protoplasma. doi: 10.1007/BF01280585

