## Supplemental data Perez Sancho-Smokvarska-Dubois for "Plasmodesmata act as unconventional membrane contact sites regulating inter-cellular molecular exchange in plants"

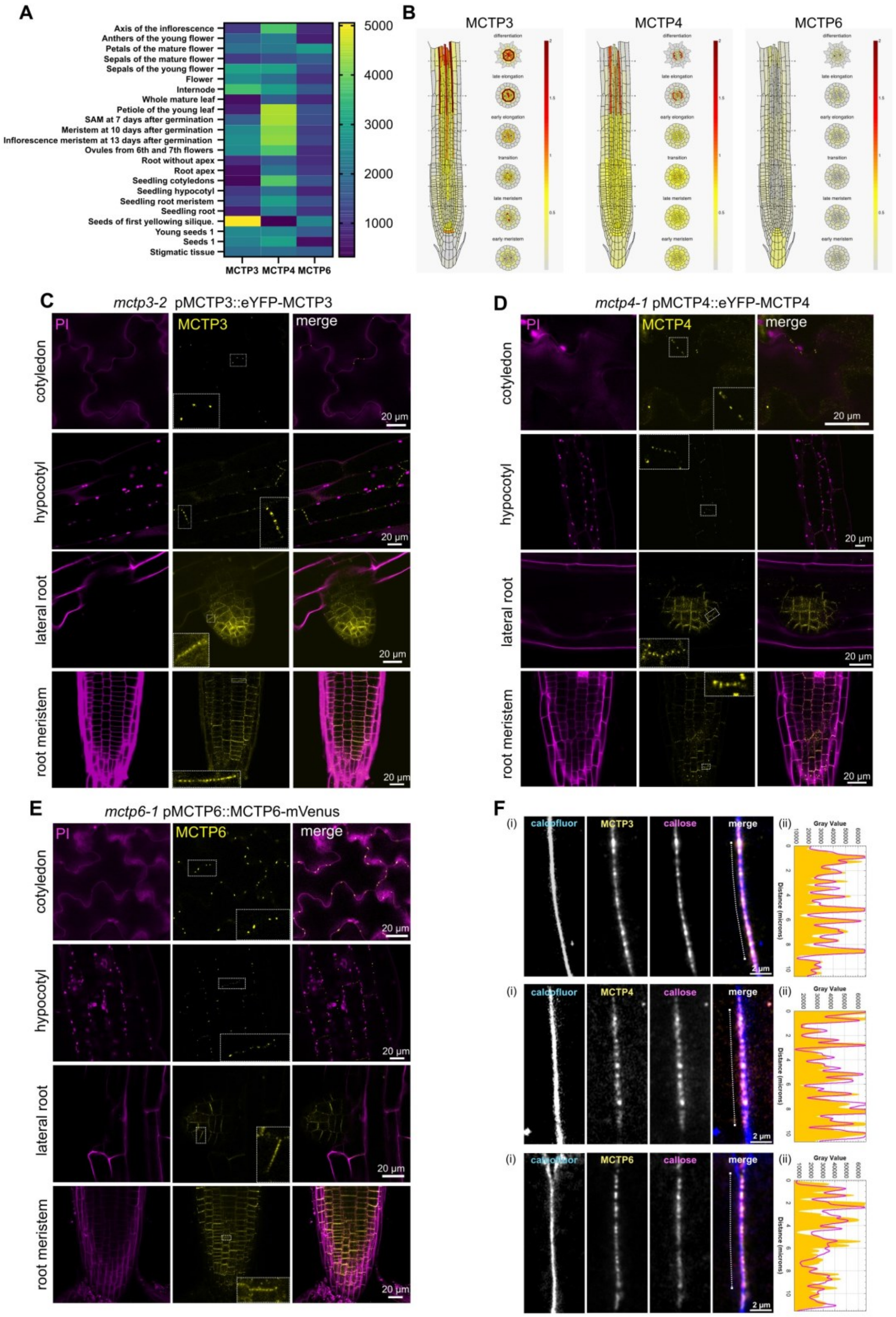

A
Axis of the inflorescence
Anthers of the young flower
Petals of the mature flower
Sepals of the mature flower
Sepals of the young flower
Flower
Internode
Whole mature leaf
Petiole of the young leaf
SAM at 7 days after germination
Meristem at 10 days after germination
Inflorescence meristem at 13 days after germination
Ovules from 6th and 7th flowers
Root without apex
Root apex
Seedling cotyledons
Seedling hypocotyl
Seedling root meristem
Seedling root
Seeds of first yellowing silique.
Young seeds 1
Seeds 1
Stigmatic tissue
MCTP3 MCTP4 MCTP6
5000
4000
3000
2000
1000
B
MCTP3
MCTP4
MCTP6
differentiation
late elongation
early elongation
transition
late meristem
early meristem
C
mctp3-2 pMCTP3::eYFP-MCTP3
PI
MCTP3
merge
cotyledon
hypocotyl
lateral root
root meristem
20 µm
D
mctp4-1 pMCTP4::eYFP-MCTP4
PI
MCTP4
merge
cotyledon
hypocotyl
lateral root
root meristem
20 µm
E
mctp6-1 pMCTP6::MCTP6-mVenus
PI
MCTP6
merge
cotyledon
hypocotyl
lateral root
root meristem
20 µm
F
(i)
(ii)
calcofluor
MCTP3
MCTP4
MCTP6
callose
merge
2 µm
Gray Value
Distance (microns)

**Figure S1. MCTP3, MCTP4 and MCTP6 are ubiquitously expressed and plasmodesmata localized.**

(A) Heat map of absolute read counts of MCTP3, 4 and 6 in different tissues and stages of plant development (data obtained from fP browser).

(B) Single-cell gene expression profile of MCTP3, 4 and 6 in the root obtained from the Root Cell Atlas (rootcellatlas.org).

(C-E) Representative confocal images of fluorescently tagged MCTP3, MCTP4 and MCTP6 under their native promoters in their respective mutant backgrounds. Propidium Iodide (PI) was used to mark cell peripheries. Color codes are indicated in the images. Insets show close up views of the regions indicated by the white dashed-line rectangles.

(F) (i) Confocal micrographs of *mctp3 pMCTP3::eYFP-MCTP3*, *mctp4 pMCTP4::eYFP-MCTP4* and *mctp6 pMCTP6::MCTP6-mVenus* co-immunolocalized with callose by anti-callose immunostaining. Calcofluor is used to visualize cell walls. (ii) Intensity plots showing YFP/mVenus (yellow) and atto594 (callose, magenta) intensity values along the walls in (i).

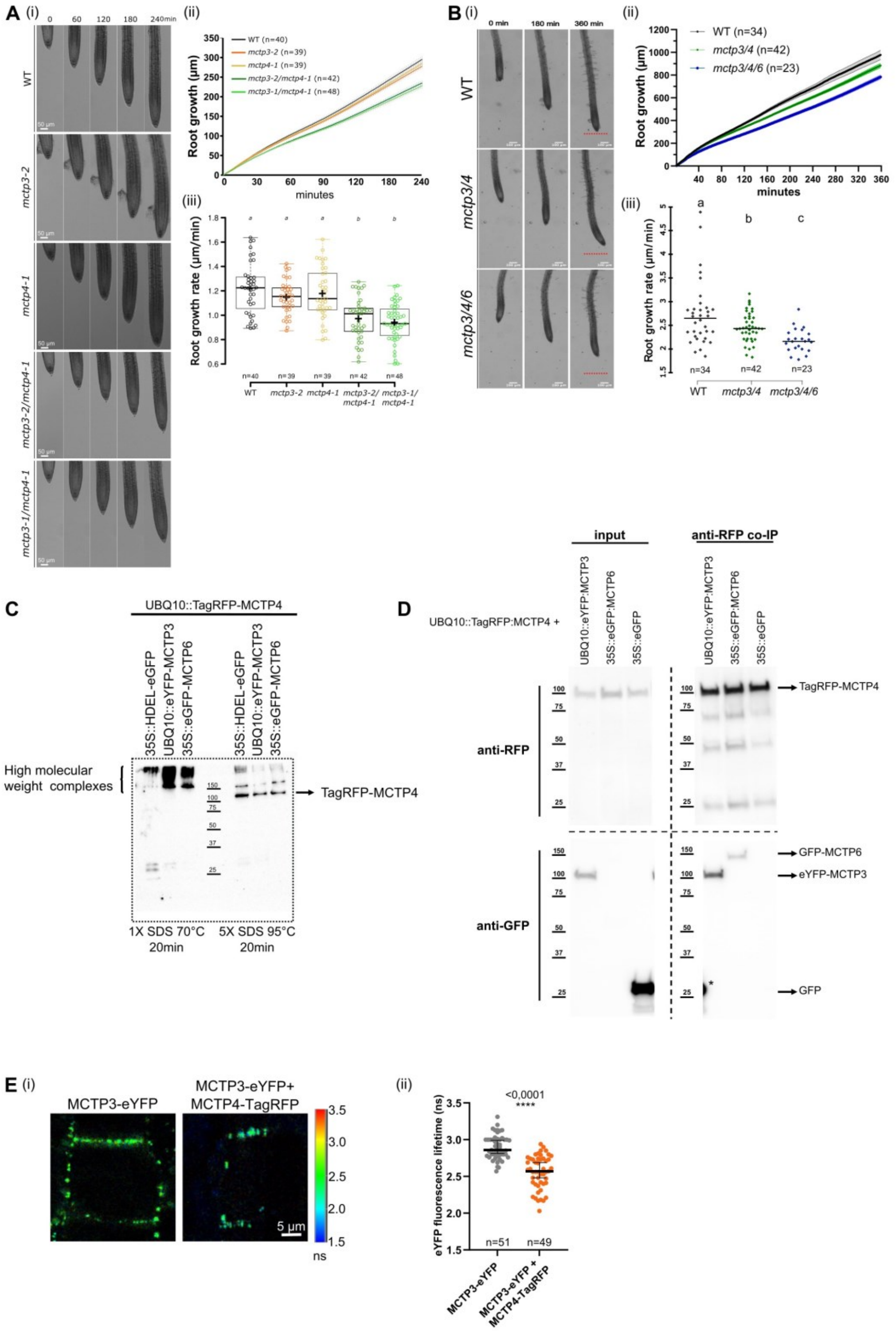
A
(i)
0
60
120
180
240min
WT
mctp3-2
mctp4-1
mctp3-2/mctp4-1
mctp3-1/mctp4-1
50 µm
(ii)
WT (n=40)
mctp3-2 (n=39)
mctp4-1 (n=39)
mctp3-2/mctp4-1 (n=42)
mctp3-1/mctp4-1 (n=48)
Root growth (µm)
minutes
(iii)
Root growth rate (µm/min)
a
a
a
b
b
n=40
n=39
n=39
n=42
n=48
WT
mctp3-2
mctp4-1
mctp3-2/mctp4-1
mctp3-1/mctp4-1
B
(i)
0 min
180 min
360 min
WT
mctp3/4
mctp3/4/6
(ii)
WT (n=34)
mctp3/4 (n=42)
mctp3/4/6 (n=23)
Root growth (µm)
minutes
(iii)
Root growth rate (µm/min)
a
b
c
n=34
n=42
n=23
WT
mctp3/4
mctp3/4/6
C
UBQ10::TagRFP-MCTP4
35S::HDEL-eGFP
UBQ10::eYFP-MCTP3
35S::eGFP-MCTP6
35S::HDEL-eGFP
UBQ10::eYFP-MCTP3
35S::eGFP-MCTP6
High molecular weight complexes
TagRFP-MCTP4
1X SDS 70°C 20min
5X SDS 95°C 20min
D
UBQ10::TagRFP:MCTP4 +
input
anti-RFP co-IP
UBQ10::eYFP:MCTP3
35S::eGFP:MCTP6
35S::eGFP
UBQ10::eYFP:MCTP3
35S::eGFP:MCTP6
35S::eGFP
anti-RFP
anti-GFP
TagRFP-MCTP4
GFP-MCTP6
eYFP-MCTP3
GFP
E
(i)
MCTP3-eYFP
MCTP3-eYFP+ MCTP4-TagRFP
5 µm
ns
(ii)
<0,0001
****
eYFP fluorescence lifetime (ns)
n=51
n=49
MCTP3-eYFP
MCTP3-eYFP + MCTP4-TagRFP

**Figure S2. MCTP3, MCTP4 and MCTP6 work together in maintaining plant fitness.**

A) Root growth assay of 5 days old WT, *mctp3-2, mctp4-1, mctp3-2/mctp4-1, mctp3-1/mctp4-1* continuously imaged for 240 min. (i) Representative images at the indicated time points after transferring for imaging. (ii) Quantification of the root length over time. (iii) Quantification of the root growth rate in µm/minutes. Number of plants per condition indicated in the figure. Statistical analysis was done with ANOVA followed by Tukey's test. Line indicates median. Cross indicates mean. Root growth assays were repeated 3 times with similar results.

(B) Root growth assay of WT, *mctp3/4* and *mctp3/4/6* continuously imaged for 360 minutes. (i). Representative images at the indicated time points. The red line marks the length of the WT root, for comparison with the mutants (ii). Quantification of root growth over time. (iii). Quantification of the root growth rate in µm/minutes. Number of plants per condition indicated in the figure. Statistical analysis was done with ANOVA followed by Tukey's test. Line indicates median. Root growth assays were repeated 3 times with similar results.

(C) Western blot of proteins transiently expressed in *N. Benthamiana* and extracted with 2 different buffers as indicated.

(D) Western blot showing co-immunoprecipitation with RFP-Trap® beads in *N. benthamiana* plants transiently expressing *pUBQ10::TagRFP-MCTP4* as bait and *p35S::GFP*, *pUBQ10::eYFP-MCTP3* or *p35S::eGFP-MCTP6* as pray proteins. Each sample consists of 40 leaf-discs extracted from 3 plants, 2 leaves per plant. Experiment was repeated 3 times with similar results. * Indicates that the black spot next to it correspond the input of 35S::GFP, which was run in the same gels (and, as it very abundant, the band extends into neighbor wells).

(E) Fluorescence lifetime of *mctp3/4/6* plants expressing *pMCTP3::eYFP-MCTP3* as donor alone or with *pMCTP4::tagRFP-MCTP4* as acceptor. (i). Representative images with color-coded fluorescent lifetime per pixel in nanoseconds (ns). (ii). Quantification of fluorescent lifetime. 3-4 plants were used per condition, 2-3 images were taken per plant and several ROIs (total indicated in the figure) were measured per image. Statistical analysis was done with Student t-test. Line indicates median, error bars indicate 95% confidence interval (CI). Experiment was repeated twice with similar results and data were pooled together.

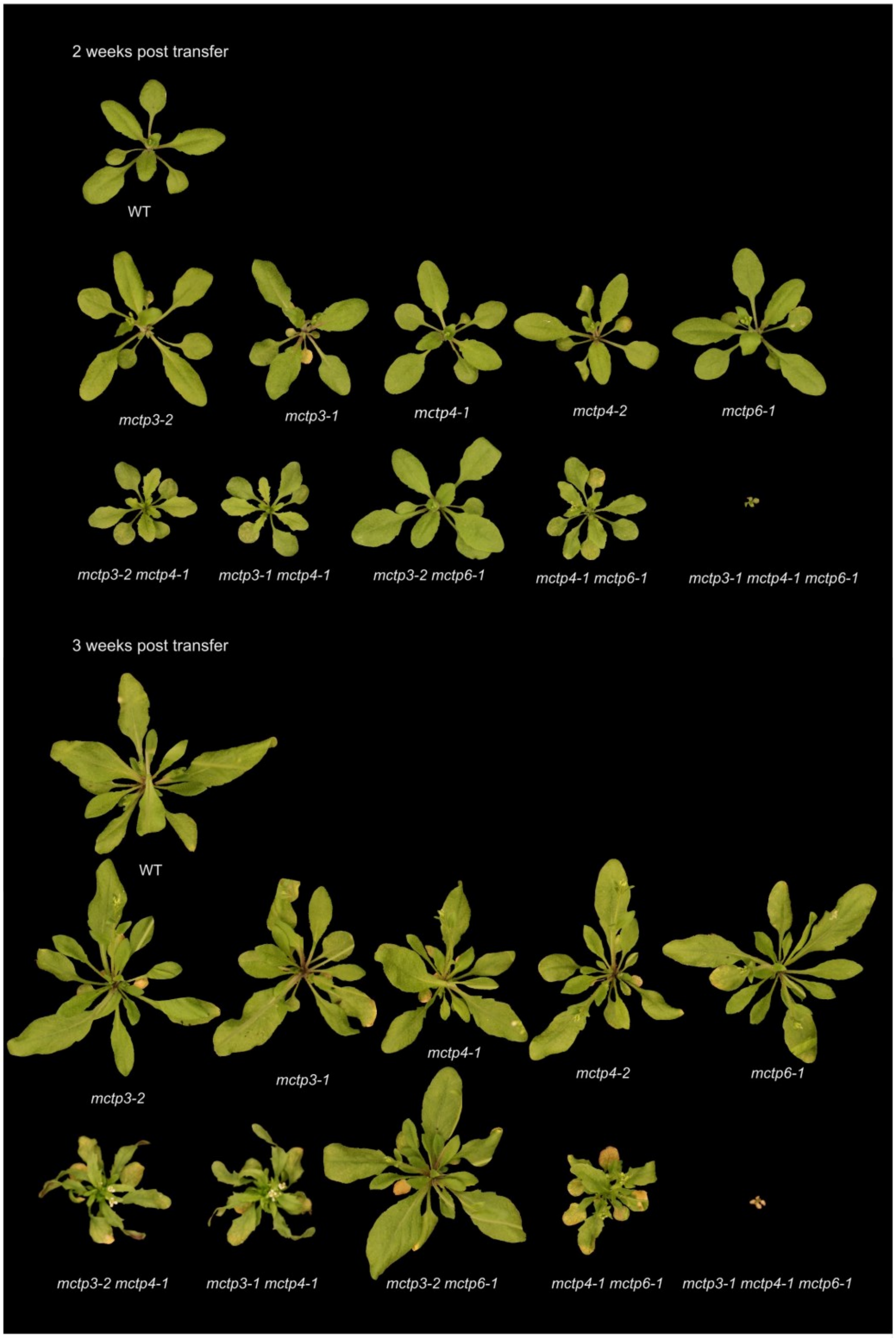
2 weeks post transfer
WT
mctp3-2
mctp3-1
mctp4-1
mctp4-2
mctp6-1
mctp3-2 mctp4-1
mctp3-1 mctp4-1
mctp3-2 mctp6-1
mctp4-1 mctp6-1
mctp3-1 mctp4-1 mctp6-1
3 weeks post transfer
WT
mctp3-2
mctp3-1
mctp4-1
mctp4-2
mctp6-1
mctp3-2 mctp4-1
mctp3-1 mctp4-1
mctp3-2 mctp6-1
mctp4-1 mctp6-1
mctp3-1 mctp4-1 mctp6-1

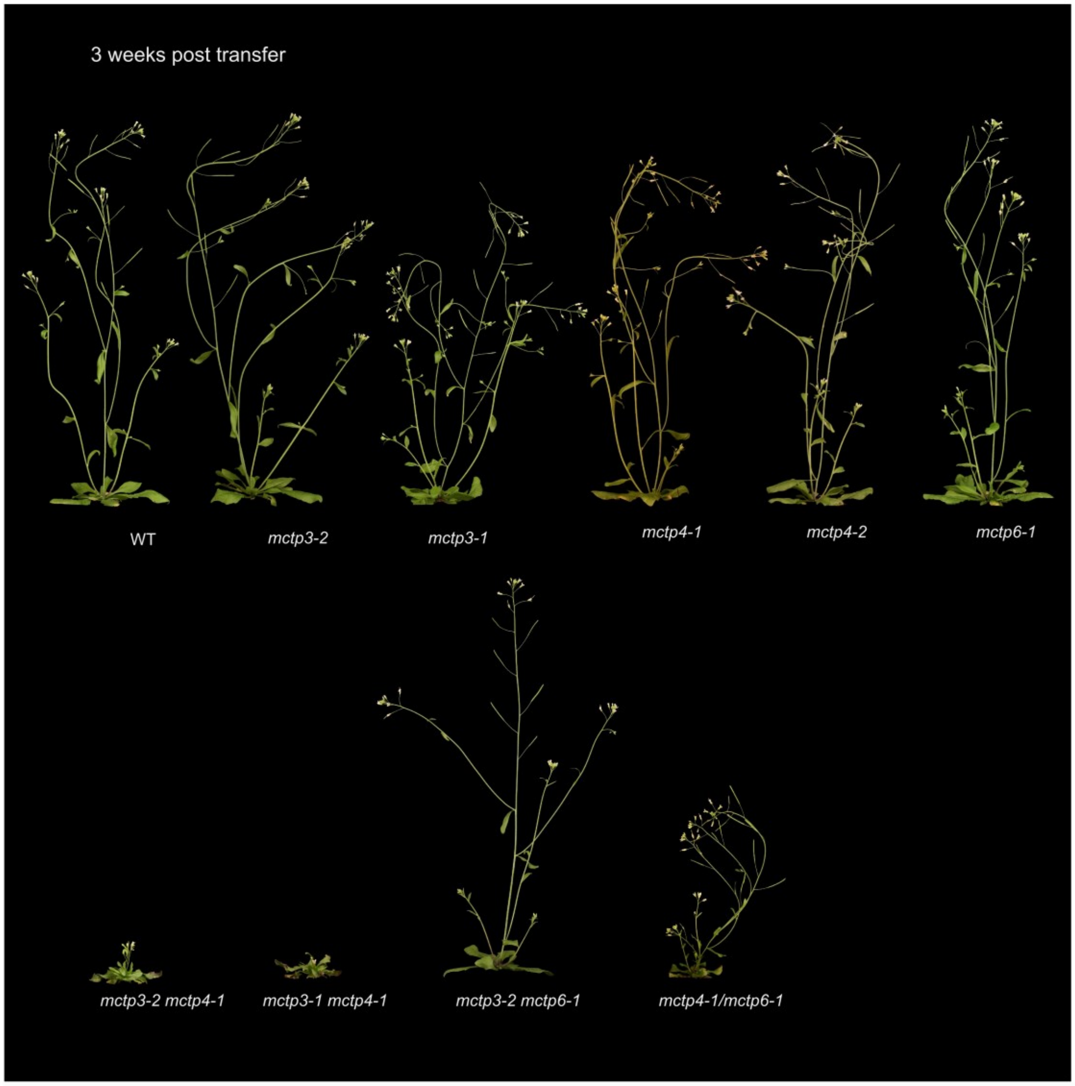


**Figure S3. Shoot growth of different MCTP mutant alleles.**

Zenithal and lateral photographs of WT, *mctp3-2, mctp3-1, mctp4-1, mctp4-2, mctp6-1, mctp3-2/mctp4-1, mctp3-1/mctp4-1, mctp3-2/mctp6-1, mctp4-1/mctp6-1 and mctp3-2/mctp4-1/mctp6-1,* 2- and 3-weeks post transfer from in-vitro plate to pots. Experiment was repeated 3 times with at least 5 plants per genotype per experiment. Representative plants were selected.

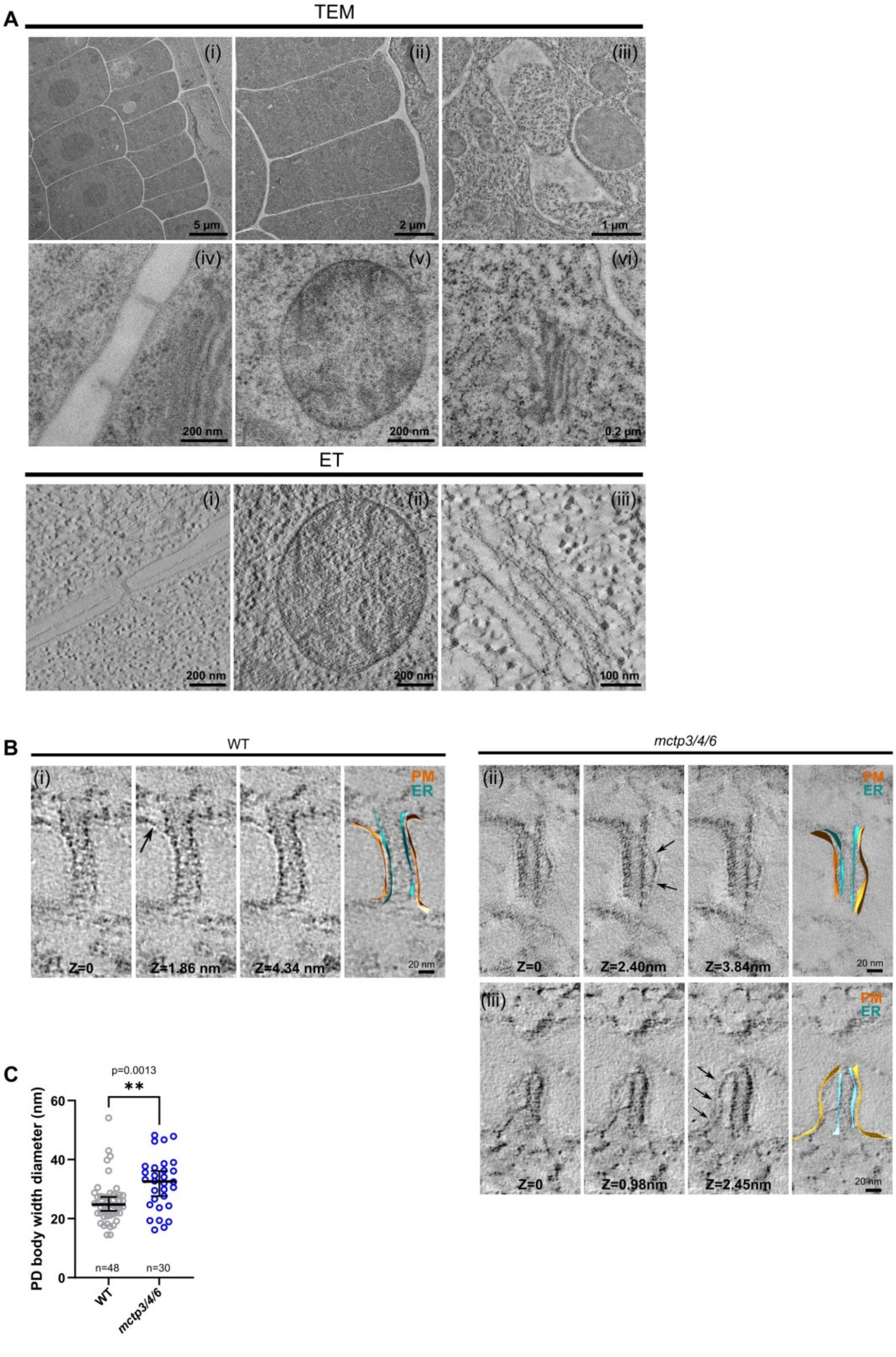
A
TEM
(i)
5 µm
(ii)
2 µm
(iii)
1 µm
(iv)
200 nm
(v)
200 nm
(vi)
0.2 µm
ET
(i)
200 nm
(ii)
200 nm
(iii)
100 nm
B
WT
(i)
Z=0
Z=1.86 nm
Z=4.34 nm
PM
ER
20 nm
mctp3/4/6
(ii)
Z=0
Z=2.40nm
Z=3.84nm
PM
ER
20 nm
(iii)
Z=0
Z=0.98nm
Z=2.45nm
PM
ER
20 nm
C
p=0.0013
**
PD body width diameter (nm)
60
40
20
0
n=48
n=30
WT
mctp3/4/6

**Figure S4. Transmission electron micrograph of an epidermal cell wall.**

(A) 2D transmission electron microscopy (TEM) and electron tomography (ET) images of high-pressure frozen, freeze-substituted WT root cells. TEM: (i) Cells overview. (ii) Magnified cells overview. (iii) Vacuole. (iv) Plasmodesmata. (v) Mitochondria. (vi) Golgi. ET: (i) Plasmodesmata. (ii) Mitochondria. (iii) Golgi.

(B) Individual tomographic slices and segmented 3D reconstructions of plasmodesmata in WT and *mctp3/4/6*. Arrows pointing to ER-PM detachment. (i) and (ii) are the same plasmodesmata as in Fig 1C, but more tomographic slices are shown here.

(C) Quantification of plasmodesmata body width in WT and *mctp3/4/6*. Linen indicate median. Error bars indicate 95 % CI.

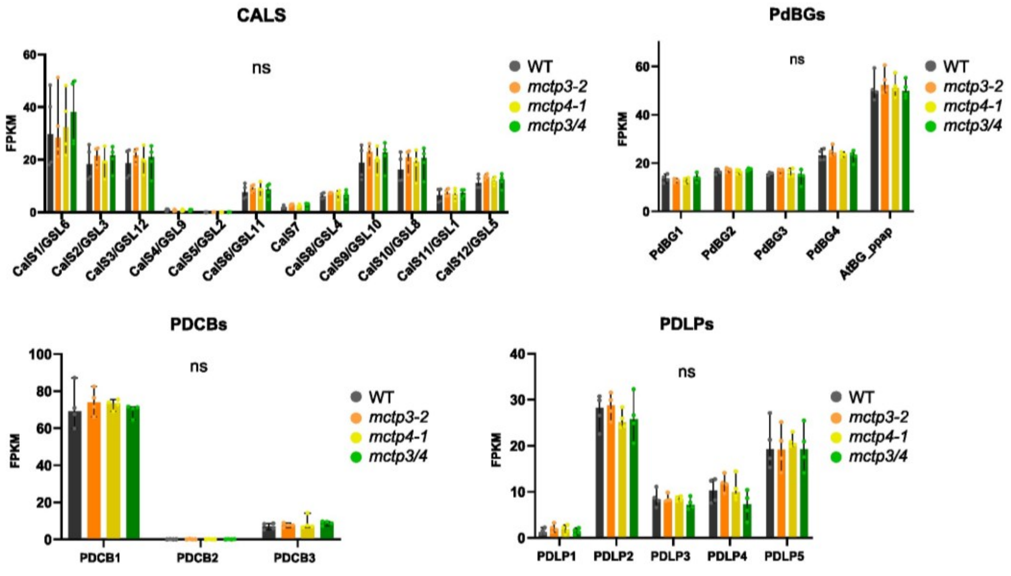


| | gene_ID |
|---|---|
| PDBC1 | AT5G61130 |
| PDBC2 | AT5G08000 |
| PDBC3 | AT1G18650 |
| PdBG1 | AT3G13560 |
| PdBG2 | AT2G01630 |
| PdBG3 | AT1G66250 |
| PdBG4 | AT1G11820 |
| AtBG_ppap | AT5G42100 |
| CalS1 | AT1G05570 |
| CalS2 | AT2G31960 |
| CalS3 | AT5G13000 |
| CalS4 | AT5G36870 |
| CalS5 | AT2G13680 |
| CalS6 | AT3G59100 |
| CalS7 | AT1G06490 |
| CalS8 | AT3G14570 |
| CalS9 | AT3G07160 |
| CalS10 | AT2G36850 |
| CalS11 | AT4G04970 |
| CalS12 | AT4G03550 |
| PDLP1 | AT5G43980 |
| PDLP2 | AT1G04520 |
| PDLP3 | AT2G33330 |
| PDLP4 | AT3G04370 |
| PDLP5 | AT1G70690 |

**Figure S5. Relative expression of proteins involved in callose deposition or degradation in WT, *mctp3-2*, *mctp4-1* and *mctp3-2/mctp4-1.***

FPKM, Fragments per kilobase mapped. Bars reach the value of the median; error bars represent 95 % CI. Experiment includes 4 biological replicates, each of them being a pool of 40 mg of 5 days old roots cut at 1 cm from the tip. Data for WT and *mctp3/mctp4* are the same as in figure 3A. Statistical analysis was done with ANOVA followed by Tukey's test.

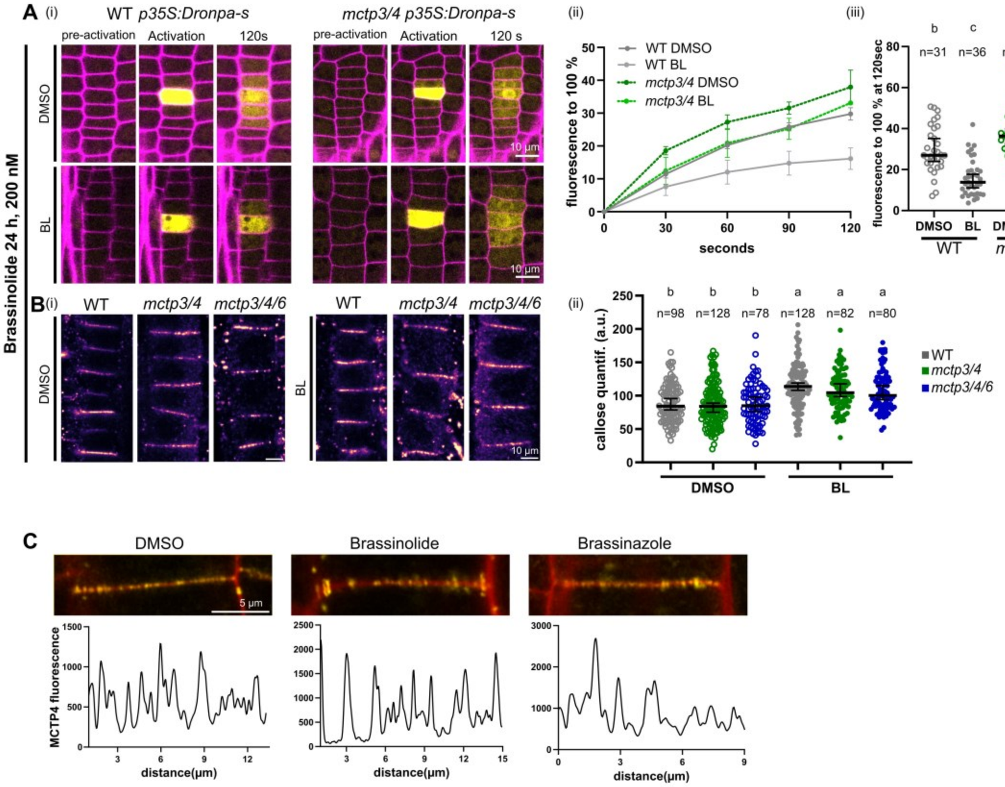


**Figure S6. Effect of brassinosteroids in cell-cell connectivity and MCTP4 accumulation**

(A-B) Brassinolide treatment; DMSO or 200 nm Brassinolide (BL) for 24 h. (A) DRONPA-s mobility assay in treated WT and *mctp3/4*. (i) Representative confocal images before activation, at activation or 120 s after two-photon activation in a single cell. (ii) Normalized fluorescence over time of DRONPA-s averaged from N±1 imaged every 15 s. Line show mean. Bars show SEM. (iii) Percentage of DRONPA-s normalized fluorescence in the N±1 cells at 120 s post activation from (ii). (B) Anti-callose immunostaining in treated WT, *mctp3/4* and *mctp3/4/6*. (i). Representative confocal images of callose immunostaining. (ii) Quantification of callose. Lines indicate median. Error bars show 95% CI. Statistical analysis was done with ANOVA followed by Tukey's test.

(C) Localization pattern at cell-cell interface of *mctp4 pMCTP4::eYFP-MCTP4* after incubation for 24h with DMSO, 200 nm BL or 1 µM BRZ. The intensity plots along a line in the apico-basal cell wall are shown below each image.

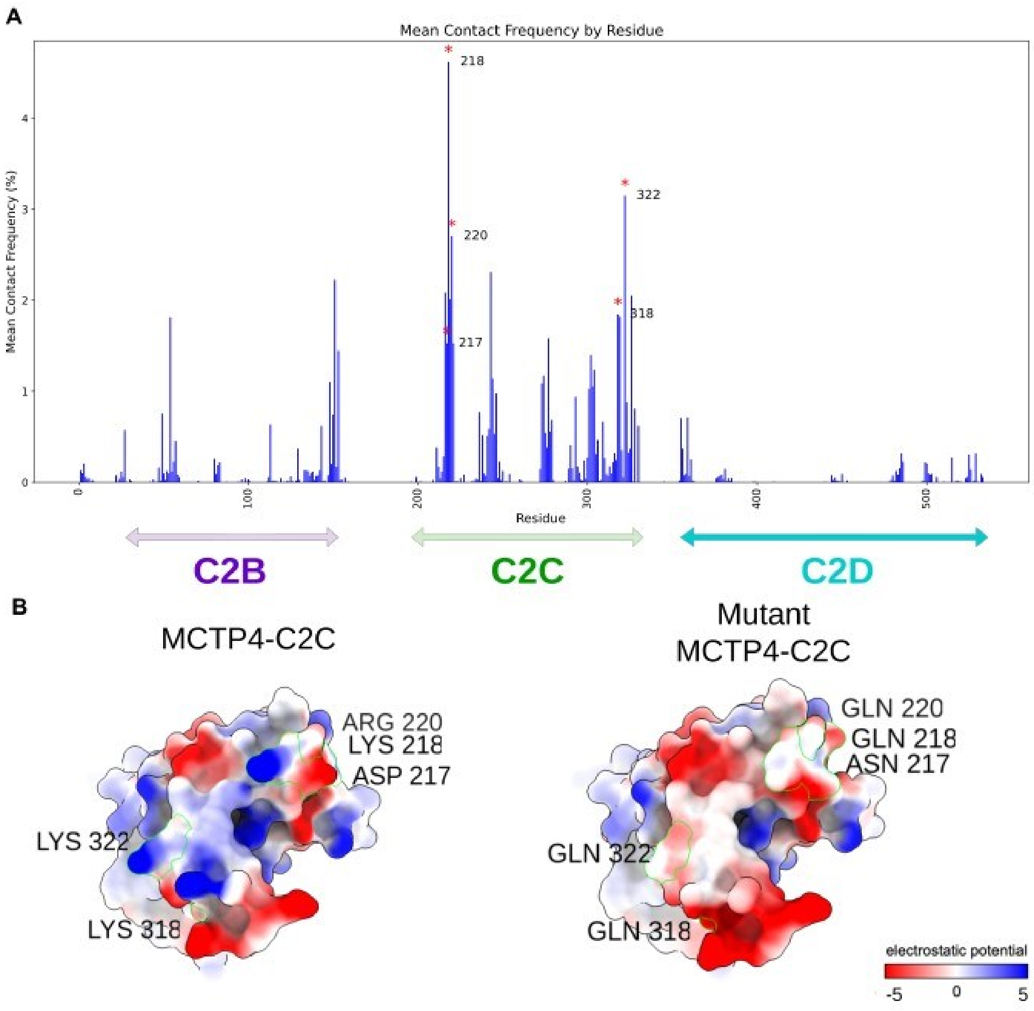


**Figure S7. Identification of putative amino acids on the C2C block responsible for PI4P binding.**

(A) Histogram of the mean contact frequency (%) for each residue across five simulations of the C2 domain of MCTP4 with a membrane composed of 2% PI4P. Two groups of contacts with higher frequency are identified, involving residues 218, 217, 220, 318, and 322.

(B) Surface representation of the C2C domain with the electrostatic potential ranging from -5 (red) to 5 (blue), created using ChimeraX. Differences between WT C2C and mutanted amino acids are shown on the electrostatic map.

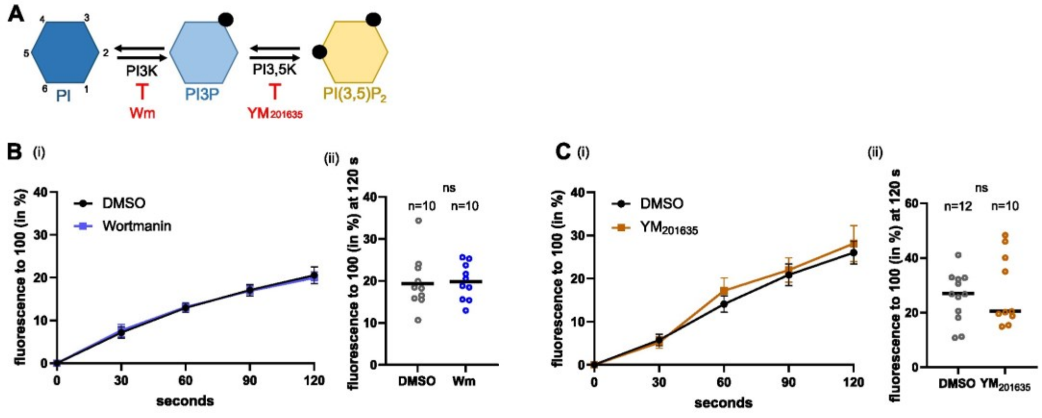


**Figure S8. Inhibition of PI3K and PI3,5K has no effect on cell-cell diffusion.**

(A) Schematic representation of the metabolism of different lipid species from phosphatidylinositol (PI) to phosphatidylinositol 3,5 bisphosphate (PI3,5P2).

(B) (i) Quantification of DRONPA-s diffusion in adjacent cell to the activated cell upon 10 µM Wortmanin (inhibitor of PI3K) for 45 minutes and DMSO treated WT plants. Lines indicate mean. Error bars indicate SEM (ii). Percentage of DRONPA-s diffusion in the adjacent cells at 120 seconds post activation between DMSO and Wortmanin treated plants.

(C) (i) Quantification of DRONPA-s diffusion in adjacent cell to the activated cell upon 2µM YM201635 (inhibitor of PI3,5K) for 1hour and DMSO treated WT plants. Lines indicate mean. Error bars indicate SEM (ii). Percentage of DRONPA-s diffusion in the adjacent cells at 120 seconds post activation between DMSO and YM201635 treated plants. Error bars show SEM. Lines indicate median. Experiment was repeated twice and data were pooled together.

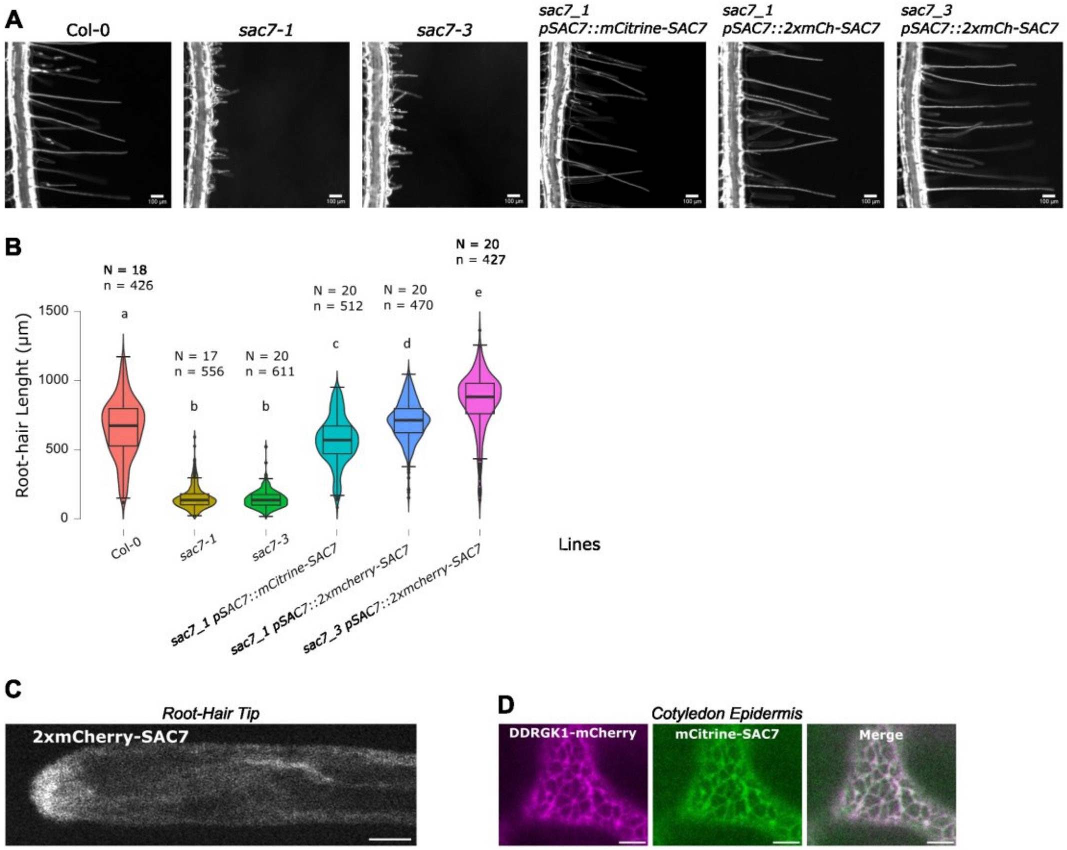


**Figure S9: SAC7 reporter lines complement *sac7* mutant root hair phenotypes**

(A) Root-hairs of WT, *sac7-1, sac7-3* and complemented lines *pSAC7::mCitrine-SAC7* in *sac7-1, pSAC7::2xmCherry-MycSAC7* in *sac7-1* and *pSAC7::2xmCherry-Myc-SAC7* in *sac7-3.*

(B) Quantification of the root-hair lenghts (µm) of the lines in (A). N= number of roots observed, n= number of root-hairs measured. Two independent experiments were done and data were pooled together.

(C) *pSAC7: :2xmCherry-Myc-SAC7* subcellular localization in root hair.

(D) Confocal images showing colocalization of *pUBQ10::DDRGK1-mCherry* and *pUBQ10::mCitrine-SAC7* after transient expression in *N. benthamiana* leaves.
